# Protein-novelty, sequence-similarity, and negative-sampling controls reveal benchmark-specific discrimination in protein–protein interaction prediction

**DOI:** 10.64898/2026.10.06.757125

**Authors:** Moruf A. Adeagbo, Halimah A. Agbaje, Sabrina A. Silveira

## Abstract

Protein–protein interaction (PPI) predictors are often evaluated with random pair splits that permit proteins to recur across training and test sets, while constructed negative pairs can introduce a separate source of benchmark-specific signal. We reassessed the Guo yeast benchmark using a matched protein-novelty curve, independently generated MMseqs2 sequence-cluster-controlled R3 variants, topology-only controls, degree- and length-matched putative negatives, handcrafted and frozen protein-language-model representations, multiple model configurations, seed analyses, and connected-component-aware inference. In the matched R1→R2→R3 comparison, AAC+CTD–MLP ROC-AUC declined from 0.697 for seen–seen pairs to 0.626 for one-unseen pairs and 0.587 for both-unseen pairs. Across independently regenerated cluster-disjoint R3 variants at 50%, 40%, 30%, and 20% MMseqs2 clustering thresholds, performance showed no consistent monotonic trend. Guo original-negative endpoints had lower positive-network degree than positive endpoints (mean 3.34 versus 6.72), and a degree-only logistic model reached ROC-AUC 0.962 on the original negatives but 0.502 on degree-matched negatives. We used degree-matched negatives consistently for R3 training and testing; six sequence-based model configurations yielded ROC-AUC 0.493–0.547; none showed corrected connected-component-aware evidence of above-chance discrimination. A PIPR multispecies analysis reproduced degree-associated label separation but also exposed cross-species pairing as an additional pooled cue. These results show that protein novelty, operational sequence-cluster separation, and negative-set topology are distinct dimensions of PPI benchmark evaluation. The Guo result is benchmark-specific and should not be generalized to sequence-based PPI prediction or protein language models overall. The dataset and source code for the experiments are publicly available at https://github.com/morufwork/guoAudit.git.

**Key Points:**

- Random pair splits, protein-disjoint splits, sequence-cluster controls, and negative-set controls answer different evaluation questions and should be reported separately.
- In Guo, AAC+CTD–MLP ROC-AUC declined from 0.697 at R1 to 0.587 at R3, while independently generated R3-50/40/30/20 variants showed no monotonic threshold trend.
- Guo original negatives carried strong positive-network-degree imbalance; a degree-only logistic model achieved ROC-AUC 0.962 on N0 but 0.502 on degree-matched N2.
- Under consistent R3/N2 evaluation, six sequence-model configurations had ROC-AUC 0.493–0.547, and the secondary PIPR analysis showed that pooled degree matching can introduce an additional taxonomic-pairing cue.

## Introduction

Protein–protein interactions (PPIs) organize signaling, metabolism, transport, macromolecular assembly, and other cellular processes [25, 18, 7]. Because experimental mapping remains incomplete and costly, researchers have developed sequence-based predictors as scalable complements to laboratory assays [16, 26, 27]. Their reported performance, however, depends on what information is allowed to recur between model development and evaluation and on how non-interacting examples are constructed. The Guo Saccharomyces cerevisiae benchmark is influential in this literature: it supplied a balanced sequence-based PPI dataset that has been reused across classical and deep-learning studies [8, 3, 1, 11, 10, 19, 13]. A first evaluation concern is protein recurrence. Randomly splitting interaction pairs can place the same protein in both training and test pairs even when no exact pair is duplicated. [21] formalized this problem for pair-input prediction through C1, C2, and C3 test cases containing two previously represented entities, one new entity, or two new entities. [9] later generalized the framework using sequence similarity rather than exact identity. Related PPI studies have also applied sequence-identity controls: DPPI used a 40% redundancy criterion and a stricter ¡25% test subset; PIPR and later DeepTrio and DPEG reported multispecies evaluations across multiple identity-filtered datasets; and INTREPPPID combined strict C3 splitting with explicit cross-split identity control [11, 6, 12, 20, 24]. Thus, neither sequence-aware C3 evaluation nor multi-threshold identity filtering is new in isolation.

A second concern is negative-set construction. Most large PPI benchmarks do not contain experimentally established non-interactions at the scale required for balanced supervised learning, so unobserved pairs are typically treated as putative negatives. If positive and constructed-negative pairs differ systematically in network degree or other endpoint properties, models may distinguish the data-construction process rather than transferable molecular compatibility. A related benchmark-construction effect has been demonstrated in antimicrobial-peptide prediction: across multiple architectures and negative-sampling procedures, [22] found that predictive performance depended substantially on how negative training and benchmark examples were selected, particularly when the two sets were generated using similar sampling schemes. Recent PPI studies have emphasized data leakage, node-degree effects, imbalances in endpoint connectivity, and topology-aware or degree-balanced negative sampling [3, 14, 15, 5]. [4] likewise highlighted the importance of rigorous dataset curation and model assessment for sequence-based PPI prediction.

These two concerns are related but not interchangeable. A protein-disjoint split can still retain a biased negative set, and a degree-balanced negative set can still permit protein recurrence across folds. Representation and architecture comparisons are therefore most interpretable when protein novelty, sequence-cluster separation, and negative-set construction are audited within the same realized benchmark framework.

Here, we integrate these controls in the Guo benchmark. We construct a matched exact-protein novelty curve with one shared training set, independently generate cluster-disjoint R3 variants at four MMseqs2 thresholds, and reconstruct alternative negative sets controlling for degree and sequence length. Handcrafted and frozen protein-language-model representations are then compared across baseline and neural models, while graph dependence is quantified using connected-component-aware inference. A PIPR multispecies analysis provides a secondary scope check. The study is intended as an integrated benchmark audit rather than a claim that any one control is novel by itself. Table 1 positions this integrated design relative to representative PPI benchmarking studies, showing which prior studies addressed protein novelty, sequence control, and negative-set or topology control and where the present audit combines those dimensions.

**Table 1.** Position of the present benchmark audit relative to representative PPI evaluation studies.

| Study | Protein novelty | Sequence control | Negative/topology control | Relevance to this study |
| --- | --- | --- | --- | --- |
| Park & Marcotte (2012); Hamp & Rost (2015) | C1/C2/C3; later generalized by sequence similarity | Exact identity and operational similarity | Random negatives / not primary focus | Foundational pair-input and sequence-aware novelty frameworks. |
| DPPI; PIPR; DeepTrio; DPEG | Varies; threshold experiments generally conventional CV | 40/;25% or Any/40/25/10/1% filtering | Not degree-matched as primary control | Establishes prior sequence-identity and multi-threshold benchmarking. |
| INTERPPIID (2024) | Strict C3 | Explicit cross-split identity control | Not primary focus | Shows strict C3 plus identity control predates the present study. |
| Bernett et al. (2024); Lannelongue & Inouye (2024) | Leakage-reduced or topology-aware evaluations | Sequence similarity controlled/minimized in relevant analyses | Node-degree/topology baselines | Directly motivates integrated leakage and topology auditing. |
| Li et al. (2025); Chatterjee et al. (2025) | Varies | Not central | Degree-balanced or topology-driven negative sampling | Shows negative construction can materially change apparent performance. |
| **Present study** | **Matched R1–R3 curve; R3 both unseen** | **Independent MMseqs2 cluster-disjoint R3–50/40/30/20** | **Degree/length matching, topology-only controls, component-aware inference** | **Integrates these controls within one Guo audit and a secondary PIPR scope analysis.** |

*Note:* C1, C2, and C3 denote the corresponding protein-pair novelty controls used in prior studies. R1–R3 denote the matched sequence-identity regimes used in the present benchmark. MMseqs2 denotes the sequence-clustering procedure used to enforce cluster-disjoint evaluation.

## Methods

### Datasets, canonicalization, and evaluation regimes

The primary analysis used the supplied Guo yeast benchmark: 2,497 protein sequences and 11,188 labeled pairs, with 5,594 positive and 5,594 benchmark-negative pairs [8]. We audited identifiers, sequences, labels, self-pairs, exact and reversed duplicates, and label conflicts, then canonicalized each undirected pair to lexical endpoint order. Identical amino-acid sequences were grouped before strict splitting. Detailed audit and canonicalization procedures are provided in Supplementary Methods M1.

We evaluated four exact-protein overlap regimes. R0 was an independently sampled stratified 80/20 random pair split and permitted protein recurrence. R1 contained test pairs for which both endpoints occurred in training, R2 exactly one unseen endpoint, and R3 two unseen endpoints. R1–R3 were generated from one partition of exact-sequence-equivalence groups and shared one training set, forming the matched protein-novelty curve. The split-validation procedure checked for and excluded forbidden protein, exact-sequence-group, exact-pair, and reverse-pair overlap. R0 is therefore a conventional reference rather than a matched fourth point on the R1→R2→R3 trajectory. Split construction, realized class/protein counts, and sequence-similarity characterization are detailed in Supplementary Methods M2.

To formalize these regimes, let *V*_train_ denote the set of protein identifiers in the training set. For any test pair (*u, v*), we define the number of unseen endpoints relative to that training set as

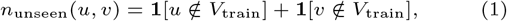

where **1**[*·*] denotes the indicator function. Accordingly, Under this definition, the three matched protein-novelty strata correspond to

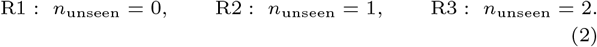

The strict both-unseen condition additionally requires complete protein disjointness between the training and test protein sets:

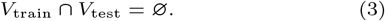

To extend exact-protein novelty with an operational sequence control, all supplied sequences were clustered independently with MMseqs2 at minimum-sequence-identity thresholds of 50%, 40%, 30%, and 20% [23]. At each threshold, MMseqs2 first generated sequence clusters using greedy set-cover clustering, 80% bidirectional coverage, sensitivity 4.0, and identity normalized by alignment length. Each resulting cluster was then assigned in its entirety to either the training or test protein pool, thereby preventing a cluster from being represented in both pools; pairs spanning the two pools were discarded. Clustering and subsequent pool assignment were rerun independently at each threshold, so R3-50, R3-40, R3-30, and R3-20 are non-nested sensitivity analyses rather than progressively filtered versions of one fixed test set. Differences among them reflect both the operational clustering criterion and realized sample composition and should not be interpreted as an isolated dose-response effect of decreasing identity. Supplementary Methods M2 reports the exact command, software build, and reproducibility checks. The complete analysis workflow is summarized in Supplementary Figure S1.

To distinguish exact protein novelty from sequence-cluster separation formally, let *c*_*τ*_ (*v*) denote the MMseqs2 cluster identifier assigned to protein *v* at clustering threshold *τ*, and let *C*_*τ*_ (*V*) denote the set of cluster identifiers represented by a protein set *V*:

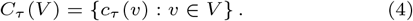

For each R3-*τ* evaluation, it is required for the training and test protein sets to contain disjoint MMseqs2 cluster sets under the corresponding operational threshold:

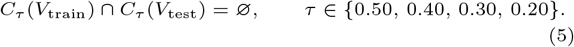

This constraint is operational under the stated MMseqs2 search, coverage, and clustering parameters; it does not imply an exhaustive pairwise identity ceiling or evolutionary non-homology. Figure 1 provides a schematic overview of these evaluation regimes, highlighting how endpoint novelty and sequence-cluster separation progressively constrain train–test overlap.

**Fig. 1.**
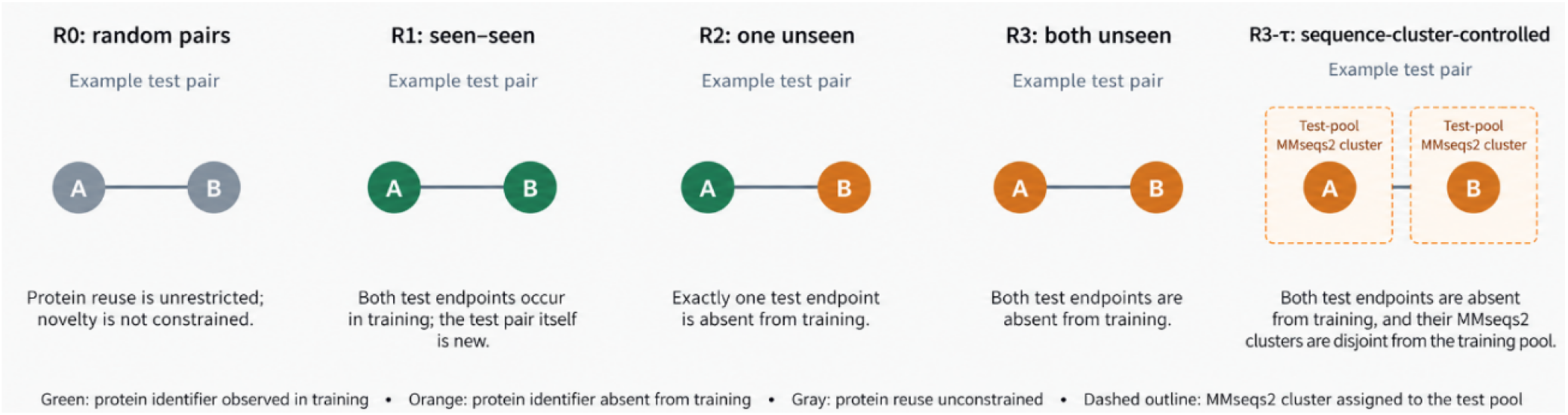
R0 permits protein recurrence; R1, R2, and R3 contain zero, one, and two unseen test endpoints, respectively. R3-*τ* additionally assigns whole MMseqs2 clusters to disjoint training and test pools under the specified operational clustering threshold.

### Representations and models

We represented each protein independently before pair construction—the principal handcrafted representation combined 20-dimensional amino-acid composition with 147 CTD descriptors. We froze the principal pretrained representation, ESM-2 (`facebook/esm2 t12 35M UR50D`; 480 dimensions), and evaluated it with mean and CLS pooling [17]. We deterministically truncated sequences longer than 1,022 residues for the main embedding pipeline, with dedicated truncation sensitivity analyses. We report a limited frozen ProtT5 cross-check only in the Supplementary Information. Undirected pair features used symmetric operators based on sums, absolute differences, elementwise products, or their combination; endpoint-swap tests verified order invariance. Representation construction, pair fusion, truncation handling, and sensitivity procedures are detailed in Supplementary Methods M3.

Because PPIs are undirected, we made the principal pair representation explicitly symmetric. For protein vectors **h**_*A*_ and **h**_*B*_, the combined operator concatenated the sum, absolute difference, and element-wise (Hadamard) product:

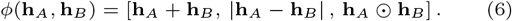

This operator is invariant to exchanging endpoints *A* and *B*. Alternative fixed operators, plain concatenation, and learned symmetric fusion are reported in Supplementary Methods M3. Baseline models were logistic regression, random forest, XGBoost, and a multilayer perceptron (MLP). Neural comparisons added a shared-encoder Siamese model, attention fusion, and a low-rank cross-interaction module. We fixed model settings across conditions and did not perform per-condition hyperparameter searches. Internal validation used training data only, and we excluded external test data from fitting transformations or calibrators. We used five model-training seeds to assess algorithmic robustness. Supplementary Methods M3 contains architecture and training details.

### Negative sets and topology controls

N0 denotes the benchmark’s original constructed negatives. We created alternative sets to expose properties that N0 might encode: N1 random unobserved pairs, N2 degree-matched pairs, N3 length-matched pairs, N5 joint degree–length-matched pairs, N6 two-sided degree substitution, and N7 joint degree–length matching with within-pair 20%-cluster exclusion. These are putative negatives, not experimentally established non-interactions. Negative-set construction and topology diagnostics are specified in Supplementary Methods M4.

N2, N3, and N5 used one-sided matched endpoint substitution. For each positive pair, the matching procedure selected one endpoint for replacement while leaving the other fixed. It drew candidates from the replaced protein’s degree decile, length decile, or joint degree–length stratum. We rejected candidates that created a self-pair, a known positive, or a duplicate negative. Failed substitutions were omitted rather than relaxing the matching rule. For R3, we generated train and test negatives within their respective protein pools and rechecked protein disjointness.

Positive-network degree was the number of observed positive partners in the deduplicated Guo positive graph. Topology-only classifiers used sum, absolute difference, minimum, and maximum endpoint degree without sequence information; length-only controls used the analogous sequence-length summaries. Because full-graph degree is label-derived and retrospective, we treat it as a benchmark diagnostic rather than a prospective feature for an unseen protein. Negative-generation robustness used 30 N2 draws with the model-training seed fixed. Exact- and nearest-degree matching, two-sided substitution, and N7 supplied additional sensitivity analyses (Supplementary Methods M4).

Let *G*^+^ = (*V, E*^+^) denote the deduplicated positive-interaction graph. The positive-network degree of protein *i*, used for matching and topology-only controls, was defined as

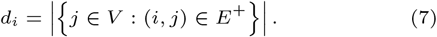

Thus, *d*_*i*_ counts protein *i*’s observed positive partners in the benchmark graph. Because this quantity comes from the positive labels, we use it as a retrospective audit variable rather than a prospective feature for an unseen protein.

### Metrics and graph-aware inference

Primary discrimination metrics were ROC-AUC and PR-AUC (average precision). We reported MCC and other threshold metrics at a fixed probability cutoff of 0.5; we fixed decision thresholds before test-set evaluation. Calibration analyses used Platt and isotonic mappings fitted on validation data from training. Score complementation was used only as an algebraic diagnostic of below-chance ordering. Supplementary Methods M5 specifies the metric, score-ordering, calibration, and operating-point procedures.

To make the score-complementation diagnostic explicit, complementing a prediction score p reverses the ROC ranking according to

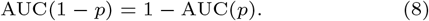

Accordingly, we used score complementation only to diagnose reversed class ordering, not as an independently validated correction.

Because PPI pairs that share endpoints are dependent, we used connected components of the undirected test-pair graph as resampling units in uncertainty analyses. A percentile connected-component bootstrap used 2,000 replicates, and a delete-one-component jackknife provided a sensitivity analysis for small component counts. We used paired contrasts when models shared an identical test set. N0–N2 comparisons resampled their different component distributions separately; because the sets share positive pairs and arise from the same biological graph, statistical independence is not established. We corrected two-sided P values with Benjamini–Hochberg within the predefined 44-test family separately for bootstrap and jackknife inference [2]. Repeated seeds are algorithmic robustness analyses, not independent biological replication. Supplementary Methods M6 provides the full hypothesis family, null values, and resampling definitions.

### Secondary PIPR benchmark

The secondary analysis used the published PIPR multispecies benchmark comprising C. elegans, D. melanogaster, and E. coli [6]. After retrieving the benchmark, protein accessions were resolved taxonomically and the original PIPR labels were retained as the N0 reference. Alternative degree-matched negative sets were then generated for both pooled and species-specific analyses. For the within-species analyses, positive-network degree and matching strata were recalculated separately for each species, replacement proteins were restricted to the same species, and new 80/20 random pair splits were generated using seed 42. In the pooled analysis, a same-species indicator was used to quantify discrimination associated with taxonomic pairing. These analyses were used to assess whether the degree-associated patterns observed in Guo were also present in an independent benchmark; however, the random pair splits still permit protein recurrence across training and test sets. Taxonomy resolution, pooled and within-species negative-set generation, and matching procedures are detailed in Supplementary Methods M7, with reproducibility and provenance described in M8.

## Results

### Protein novelty reduces discrimination; sequence-cluster sensitivity is non-monotonic

Canonicalization removed 24 duplicate pairs, leaving 11,164 unique pairs (5,581 positive; 5,583 constructed negative). The positive graph contained 2,217 observed nodes, 5,581 edges, 98 connected components, a mean degree of 5.03, and a maximum degree of 15. The matched R1–R3 evaluations shared 5,703 training pairs and included 1,426, 3,570, and 462 test pairs, respectively; R0 included 8,931 training and 2,233 test pairs. Supporting tables covering dataset/network integrity, R0–R3 split composition, cluster-controlled split integrity, and post-audit counts and sequence lengths are provided in Supplementary Tables S1–S4. Figure 2 summarizes the post-audit class, length, and degree distributions. Degree zero denotes absence from the observed positive graph, not evidence of biological isolation.

**Fig. 2.**
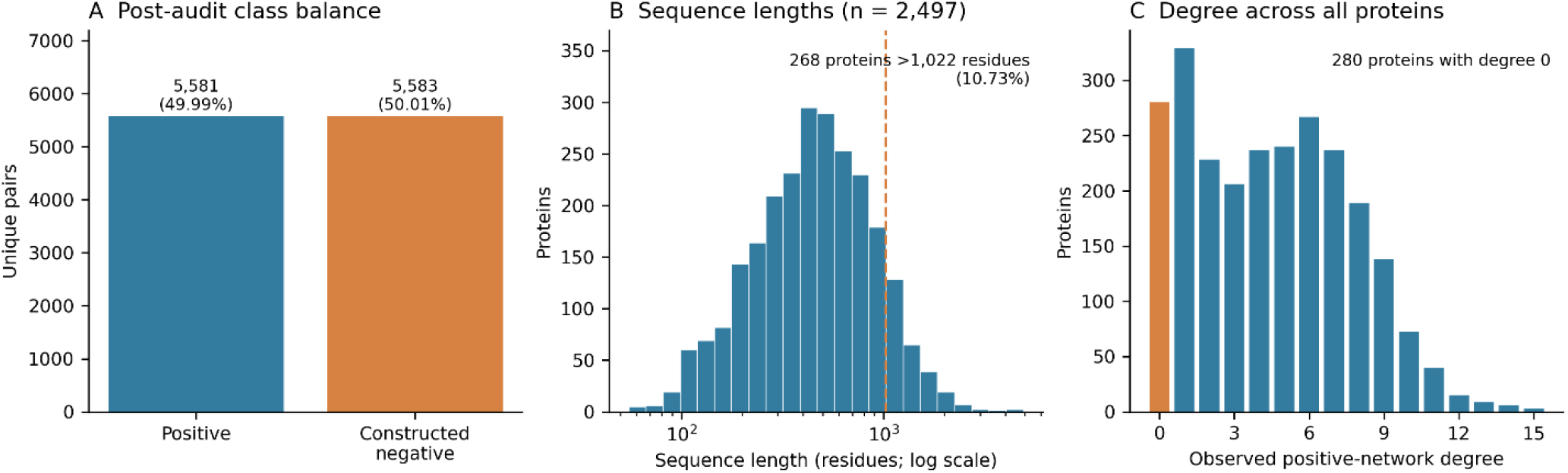
Yeast benchmark distributions after the integrity audit. (A) Class counts after removal of 24 duplicate pairs. (B) Sequence-length distribution across all 2,497 protein identifiers; the dashed line marks the 1,022-residue embedding limit. The horizontal axis is logarithmic, and bars show counts in logarithmically spaced bins. (C) Positive-network degree across the same protein universe, including 280 zero-degree proteins. Each protein contributes once to panels B–C; these distributions differ from the endpoint-occurrence-weighted summaries used in the negative-set audit.

For AAC+CTD–MLP, R0 ROC-AUC was 0.701. Within the matched R1→R2→R3 curve, ROC-AUC declined from 0.697 to 0.626 and 0.587 as the number of unseen endpoints increased from zero to two. MCC declined from 0.305 to 0.191 and 0.129, while PR-AUC declined from 0.684 to 0.587 and 0.549. Random forest showed a similar trajectory; XGBoost did not decrease monotonically, although its R3 value remained below R1. Because R1–R3 share training data but contain different test pairs and class compositions, the curve describes novelty-associated performance and is not a causal estimate of protein recurrence. Supplementary Table S5 reports full AAC+CTD baseline metrics and realized pair counts for these regimes.

The independently regenerated R3-50, R3-40, R3-30, and R3-20 variants passed all protein, exact-sequence, cluster, pair, and reversal overlap checks. Their train/test pair counts were 7,227/450, 7,111/423, 7,174/419, and 7,043/445. Across four AAC+CTD models, ROC-AUC ranges were 0.583–0.630, 0.586–0.641, 0.544–0.576, and 0.586–0.660, respectively. R3-20 was not uniformly the most difficult condition, and model rankings varied across thresholds. The absence of a monotonic pattern therefore reflects sensitivity across independently generated operational cluster definitions rather than evidence that sequence similarity is irrelevant. Representation effects were also regime-dependent: with the same MLP, ESM-2 CLS and mean pooling achieved ROC-AUC 0.755 and 0.730 at R0, but 0.562 and 0.617 at R3, versus 0.701 and 0.587 for AAC+CTD; mean and CLS pooling also exchanged relative rank across R3-*τ* conditions. Supplementary Table S6 and Figure S2 show complete representation comparisons. Figure 3 summarizes these results across ROC-AUC, PR-AUC, and MCC, placing the matched protein-novelty curve alongside the independently generated sequence-cluster-controlled R3 sensitivities.

**Fig. 3.**
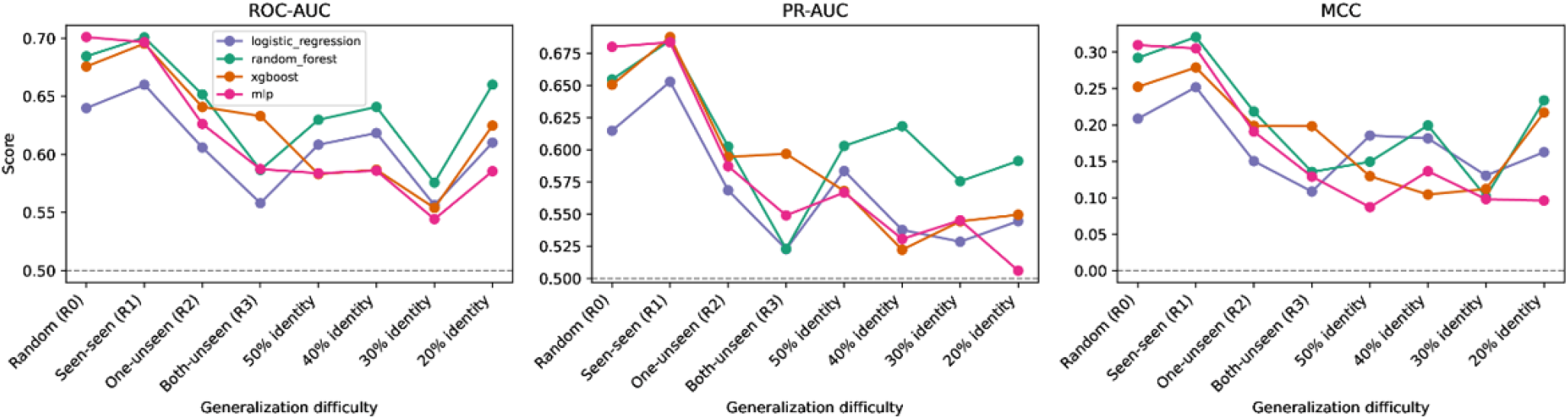
Generalization across protein-novelty and sequence-cluster regimes. ROC-AUC, PR-AUC, and MCC for AAC+CTD baseline models. R0 is a separate random-pair reference; R1*→*R2*→*R3 is the matched novelty curve; R3-50 through R3-20 are independently generated cluster-disjoint R3 sensitivities.

### Guo original negatives carry strong degree-associated label separation

Positive-pair endpoint occurrences had a mean positive-network degree of 6.72, compared with 3.34 for N0 endpoints, corresponding to an endpoint-occurrence standardized mean difference of 1.160. Random negatives (N1) and length-matched negatives (N3) reduced but did not eliminate this imbalance, whereas degree-matched N2 and joint degree–length-matched N5 closely aligned with the positive endpoint-degree distribution. Because positive-network degree was calculated from the observed positive-interaction graph, it is treated as a retrospective benchmark-construction diagnostic rather than as an independent biological feature or prospective predictor. Endpoint-occurrence degree summaries, distributional balance diagnostics, and degree-only classifier performance under decile, exact, and nearest-degree matching are reported in Supplementary Tables S7–S9. Figure 4 visualizes the same endpoint-degree distributions, showing the strong shift in N0 and the near alignment achieved by N2 and N5.

**Fig. 4.**
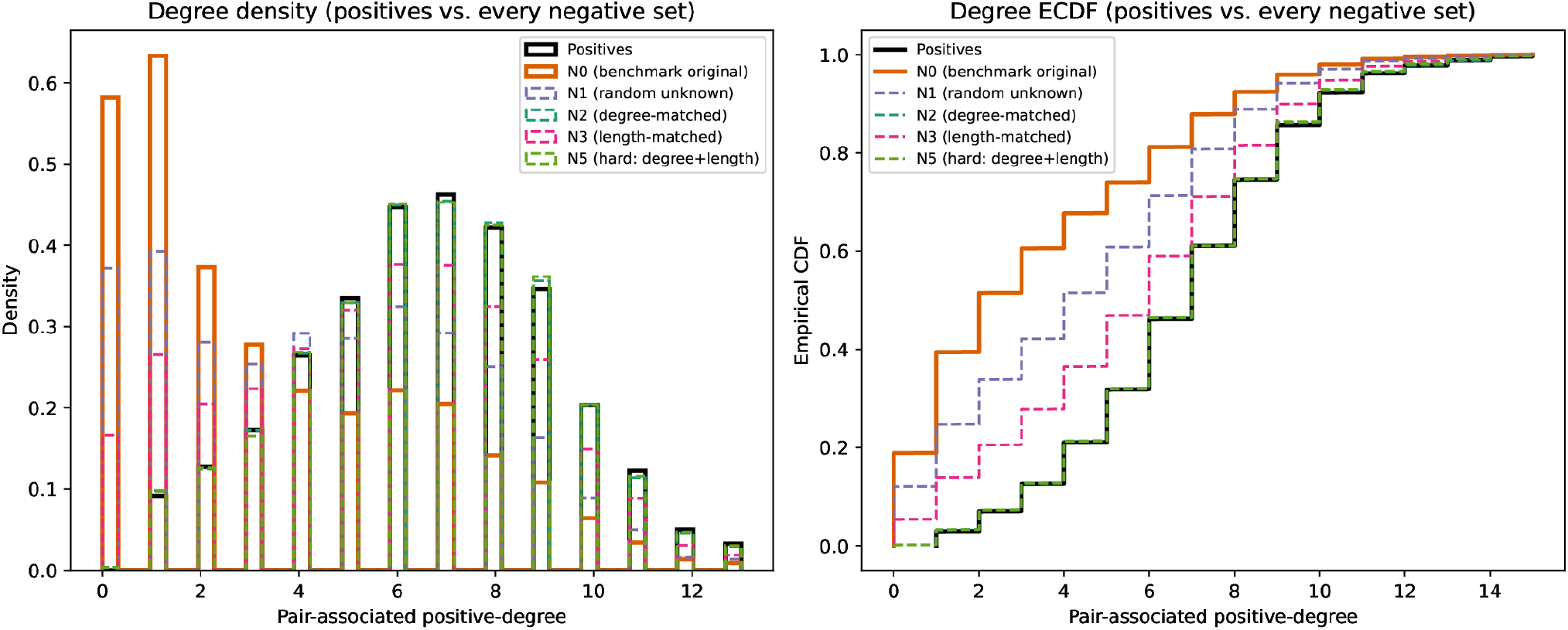
Endpoint positive-network degree distributions across positive and constructed-negative pairs. The left panel shows the normalized distribution of endpoint positive-network degree, and the right panel shows the corresponding empirical cumulative distribution functions (ECDFs). Original benchmark negatives (N0) are strongly shifted toward lower-degree endpoints relative to positive pairs. Random negatives (N1) and length-matched negatives (N3) reduce but do not eliminate this difference, whereas degree-matched N2 and joint degree–length-matched N5 closely align with the positive endpoint-degree distribution. Positive-network degree is derived retrospectively from the observed positive-interaction graph and is used here as a benchmark diagnostic rather than as a prospective predictor.

A logistic classifier using only four endpoint-degree summaries reached ROC-AUC 0.962 and MCC 0.779 against N0; a degree-only random forest reached ROC-AUC 0.966. Against N2, ROC-AUC fell to 0.502–0.504 and remained near chance with joint degree–length matching. Exact- and nearest-degree matching were comparable, whereas length-only controls were much weaker. These results show that label-derived degree is sufficient to reproduce much of N0 separability in this retrospective benchmark. This does not mean sequence models literally encode degree or that degree is uniquely causal. Supporting analyses are provided in Supplementary Tables S9–S13: Table S9 covers the degree-matching rule; Table S10, degree- and length-controlled analyses; Table S11, ESM-2, degree, and combined-feature analyses; Table S12, the N2 versus N6 comparison; and Table S13, N7 cluster-exclusion sensitivity.

Sequence-model performance also changed with the negative definition. In the distribution-shift experiment, an N0-trained ESM-2–MLP declined from ROC-AUC 0.754 on N0 to 0.647 on N1, 0.565 on N3, 0.467 on N2, and 0.494 on N5. The reverse experiment in which models were trained on alternative negatives and evaluated against the common N0 test set likewise showed strong sensitivity to negative construction. Because these experiments change the training or evaluation distribution, they diagnose benchmark dependence rather than provide a single deployable operating condition. Supplementary Table S14 reports complete bidirectional negative-set training/evaluation results. Figure 5 visualizes these complementary distribution-shift experiments, contrasting the degree-only control with the ESM-2–MLP sequence model as the negative-set definition changes.

**Fig. 5.**
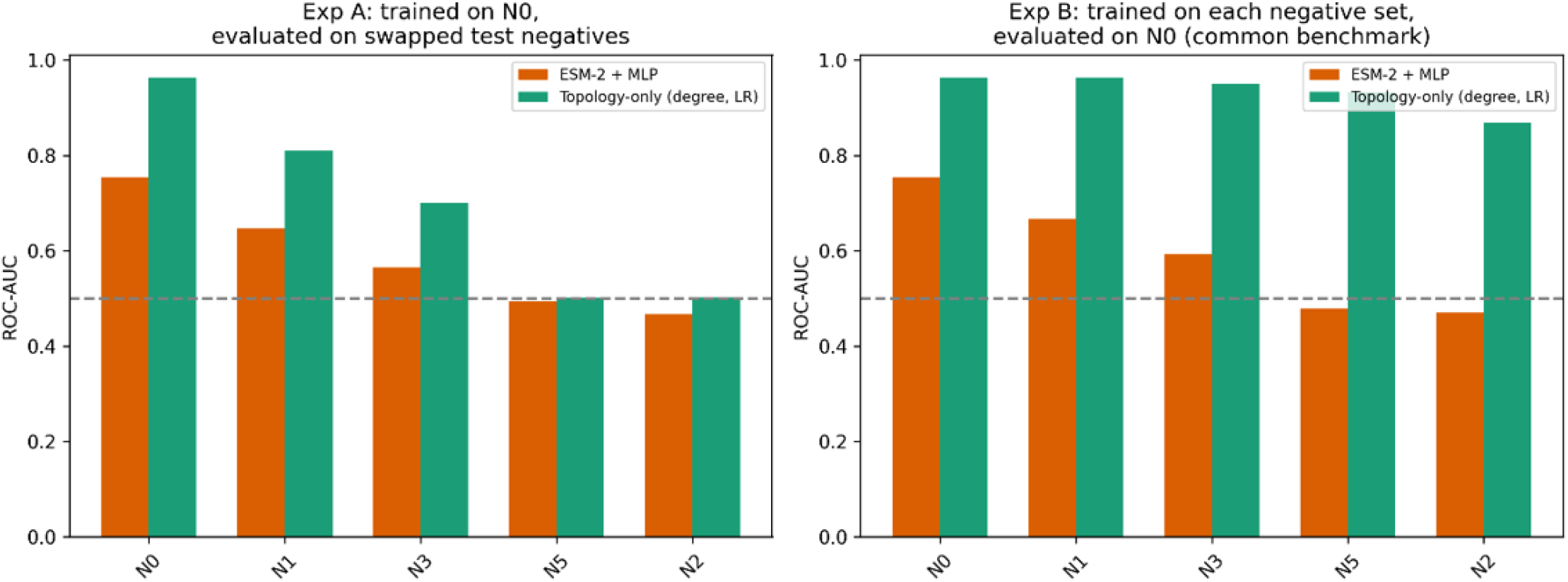
Degree-only and sequence-model sensitivity to the negative definition. Models trained on N0 are evaluated after test-negative replacement (left), and models trained on each negative set are evaluated on the common N0 test set (right). Sequence-model results use ESM-2–MLP; numerical values are in Supplementary Tables S11 and S14.

**Fig. 6.**
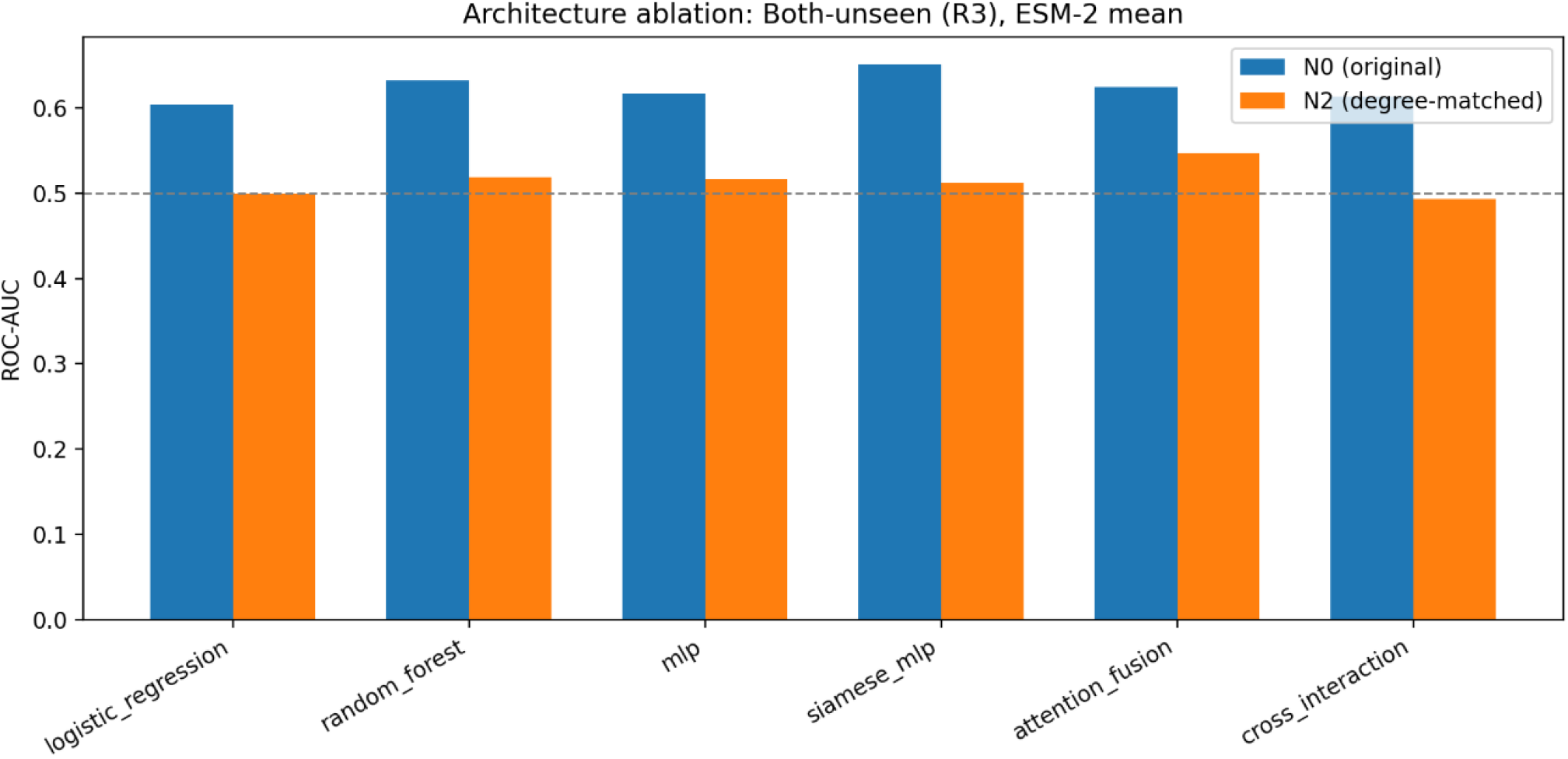
R3 model-configuration comparison under N0 and N2. Baseline and neural models retain modest discrimination under N0 but have ROC-AUC point estimates close to 0.5 under degree-matched N2. Full single- and multi-seed results are in Supplementary Tables S16 and S17

### Consistent degree matching removes the main Guo representation and model advantages in point estimates

When we used each negative definition consistently in both training and evaluation, Guo R0/N2 ROC-AUC was 0.455 for AAC+CTD and 0.463 for ESM-2 mean, while R3/N2 was 0.517 for both representations. Thus, the R0-versus-R3 gap and the ESM-2-versus-AAC+CTD advantage observed under N0 were not retained in these degree-matched point estimates. The full consistent-negative representation comparison is in Supplementary Table S15 and Supplementary Figure S3.

Across six R3/N2 model configurations, single-seed ROC-AUC values were 0.500 for logistic regression, 0.518 for random forest, 0.517 for MLP, 0.512 for the Siamese model, 0.547 for attention fusion, and 0.493 for cross-interaction. PR-AUC ranged from 0.498 to 0.544 against a positive prevalence of 0.500. Across five training seeds, mean ROC-AUC ranged from 0.500 to 0.525. Across 30 independently generated N2 sets, R3 ROC-AUC was 0.489 ± 0.023 for ESM-2 logistic regression, 0.505 ± 0.023 for ESM-2–MLP, and 0.504 ± 0.003 for the degree-only control. These repetitions show algorithmic stability on the same graph, not independent biological replication. Single-seed and five-training-seed model-configuration results are in Supplementary Tables S16 and S17.

Component-aware inference was constrained by graph structure: strict N0 and N2 test graphs supplied only 19 and five usable connected components, respectively, with most pairs in a giant component. After Benjamini–Hochberg correction, no tested N2 model configuration showed corrected evidence of above-chance discrimination with either resampling method. N0–N2 point-estimate differences were directionally consistent but not significant after correction. This limits inferential strength and does not establish equivalence to chance or the absence of biological sequence signal. Detailed inferential analyses are presented in Supplementary Tables S18–S23, covering the test family (Table S18), above-chance tests (Table S19), giant-component exclusion (Table S20), the paired Siamese–LR contrast (Table S21), N0–N2 contrasts (Table S22), and leave-one-component-out sensitivity analysis (Table S23). The corresponding jackknife intervals are shown in Supplementary Figure S4.

Additional sensitivity analyses did not change the main interpretation. Supporting analyses are provided in Supplementary Tables S24–S33: Table S24 reports the 30 N2-generation seeds; Table S25, the degree source; Table S26, score inversion; Table S27, raw calibration intervals; Table S28, threshold and precision–recall operating points; Table S29, pair fusion; Table S30, truncation; Table S31, raw versus calibrated predictions; Table S32, ECE binning; and Table S33, separate handcrafted baselines. Supplementary Figure S5 presents the corresponding inversion-diagnostic score distributions.

### PIPR reproduces degree-associated separation but reveals an additional taxonomic cue

In the pooled PIPR benchmark, degree-only logistic regression declined from ROC-AUC 0.849 under N0 to 0.535 under N2. Sequence models retained pooled N2 discrimination: AAC+CTD–MLP reached 0.646, ESM-2 logistic regression 0.682, and ESM-2–MLP 0.698. However, 14,284 of 31,678 reconstructed pooled N2 negatives (45.1%) joined proteins from different species, and a same-species indicator alone achieved ROC-AUC 0.729 on the pooled test pairs assigned to the three target species. Pooled residual discrimination is therefore compatible with a strong taxonomic-pairing cue and does not establish molecular compatibility independent of benchmark construction. Supplementary Tables S34–S38 provide the corresponding pooled analyses: Table S34 reports performance under decile, exact, and nearest-degree matching; Table S35 presents pooled N0 and degree-matched N2 model performance; Table S36 summarizes pooled endpoint-degree balance; Table S37 details the taxonomic composition of the reconstructed pooled N2 negatives; and Table S38 reports the same-species indicator diagnostic.

Within-species reconstruction removed cross-species pairing. Under degree-decile matching, ESM-2–MLP ROC-AUC was 0.596 in C. elegans, 0.567 in D. melanogaster, and 0.647 in E. coli. Degree-only values also moved toward chance relative to N0. Exact matching omitted some proposed negatives and changed class composition, whereas nearest-degree matching retained one proposed negative per positive. These remain descriptive single random-pair splits with protein recurrence and do not test unseen-protein generalization or statistically compare species. Supporting analyses are provided in Supplementary Tables S39–S45: Table S39 reports dataset sizes; Table S40, retained and excluded pairs; Table S41, C. elegans results; Table S42, D. melanogaster results; Table S43, E. coli results; Table S44, degree balance and omissions; and Table S45, train/test composition and protein overlap. Supplementary Figure S6 summarizes the central cross-analysis results, while Figure 7 provides a direct comparison of the within-species N0 and N2 results.

**Fig. 7.**
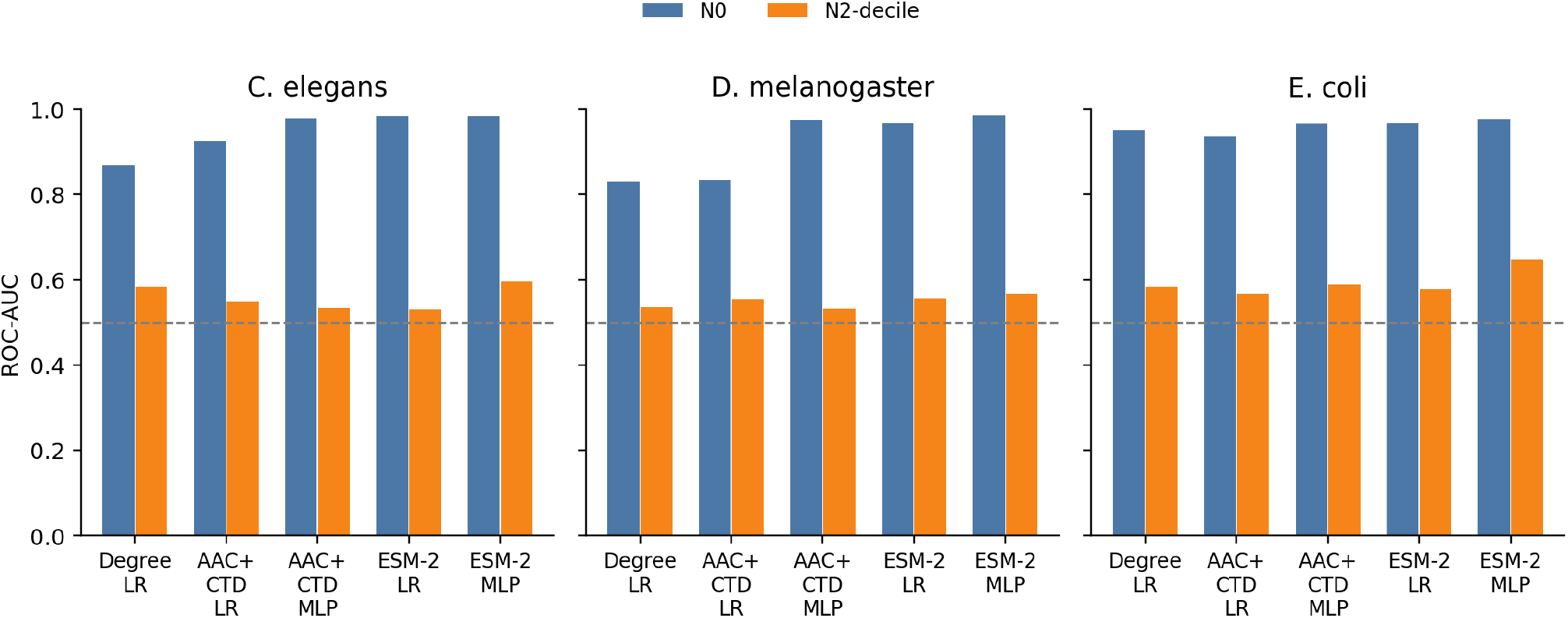
Species-stratified PIPR replication. ROC-AUC under original (N0) and within-species degree-decile-matched (N2) negatives for degree-only logistic regression and AAC+CTD/ESM-2 models. Each species uses an independently reconstructed random pair split; bars are descriptive point estimates.

## Discussion

The Guo analyses separate two evaluation problems that can coexist: protein recurrence across training and test pairs, and the construction of negative examples with network properties that differ from those of positive pairs. Performance decreased across the matched R1→R2→R3 protein-novelty strata, while label-derived degree alone strongly separated the original benchmark labels. Because R1–R3 shared one training set but contained different test pairs and class compositions, this gradient describes novelty-associated performance rather than an isolated causal effect of protein recurrence. When degree-matched negatives were used consistently, sequence-model point estimates under strict R3 remained close to 0.5 across the evaluated representations, model configurations, and seed analyses. These observations support auditing split design and negative construction before interpreting high benchmark discrimination as evidence of transferable molecular compatibility.

The sequence-cluster results should be interpreted as sensitivity analyses rather than as a dose-response experiment on sequence identity. R3-50 through R3-20 were independently clustered and partitioned, so changing the MMseqs2 threshold also changed the realized training and test compositions. Consequently, performance differences across these conditions reflect both the operational clustering criterion and the particular proteins and interaction pairs retained in each dataset. The observed non-monotonic pattern therefore cannot be attributed solely to progressively decreasing train–test sequence similarity. Moreover, MMseqs2 thresholds under the stated coverage and search parameters define operational cluster separation rather than exhaustive pairwise identity ceilings or evidence of evolutionary non-homology. Prior literature already established sequence-aware C3 evaluation and multi-threshold identity filtering; the contribution here is their integration with the same Guo protein-novelty, negative-set, and topology audit.

The degree result is also retrospective. Positive-network degree was calculated from the same incomplete observed positive graph that supplies the benchmark’s positive labels. High N0 discrimination by degree-only classifiers therefore establishes an exploitable label-associated property within this benchmark; it does not provide a deployable predictor for proteins without observed interactions and does not show that sequence models use degree internally or that degree is the unique causal source of their discrimination. Annotation density, experimental attention, detectability, biological centrality, or other correlated properties may contribute but were not independently measured. Likewise, degree matching changes the candidate negative-pair population and may alter correlated properties. It should therefore be interpreted as a benchmark-construction stress test rather than a causal intervention on biological interaction propensity.

The component-aware analysis makes the inferential limitation explicit. Large pair counts do not translate into many connected-component resampling units when most test pairs belong to one giant component. Bootstrap and jackknife conclusions diverged for some N0 tests, no tested N2 model configuration met the corrected above-chance criterion, and the N0–N2 contrast was not statistically established after correction. Moreover, the N0–N2 procedure is a working unpaired approximation because the two evaluations use different negative pairs but share positive pairs and originate from the same biological graph. Nonsignificant results should therefore not be interpreted as evidence of equivalence to chance or absence of a difference. Consistency across model-training and negative-generation seeds provides evidence of algorithmic robustness on the same graph but does not add independent biological replication.

The PIPR analysis further limits generalization from Guo. Degree-associated label separation was also present in PIPR, whereas sequence models retained pooled discrimination after degree matching. However, 45.1% of the reconstructed pooled N2 negatives connected proteins from different species, and a same-species indicator alone achieved substantial discrimination. The pooled benchmark therefore contains a strong taxonomic pairing cue that could contribute to residual sequence-model discrimination; these analyses do not establish that the sequence models specifically exploited that cue. Within-species reconstruction removed cross-species pairing and retained modest residual sequence discrimination, but the species-specific evaluations used single random pair splits in which proteins could recur across training and test pairs. They therefore do not establish generalization to unseen proteins. Taken together, the Guo and PIPR analyses do not support a universal conclusion about sequence-based PPI prediction or protein language models.

For PPI benchmark reporting, the practical implication is to treat protein overlap, operational sequence-cluster separation, negative-set provenance, endpoint-property balance, topology-only baselines, and graph-aware uncertainty as distinct evaluation dimensions. Representation and model comparisons are more interpretable when they use the same realized partitions and are examined under more than one defensible negative definition. Ranking performance, probability calibration, and fixed-threshold metrics should also be reported separately, particularly when constructed approximately balanced evaluation sets differ from prospective interactome prevalence.

## Conclusion

An integrated audit of the Guo benchmark shows that protein novelty and negative-set topology alter apparent sequence-based PPI discrimination in different ways. Discrimination declined across a matched seen–seen to both-unseen curve; independently generated cluster-disjoint R3 variants showed no monotonic threshold trend; and original negatives carried a strong degree-associated imbalance that degree-only models could exploit. Under consistently degree-matched R3 evaluation, model point estimates were near chance, and corrected component-aware tests did not show above-chance discrimination. PIPR reproduced degree-associated separation but also revealed a taxonomic-pairing cue in pooled negatives. These findings motivate benchmark designs that report protein and sequence-cluster overlap, negative provenance, topology controls, and graph-aware uncertainty before attributing performance to transferable molecular interaction signal.

## Data Availability

All datasets and source code supporting this study are publicly available at the GitHub repository: https://github.com/morufwork/guoAudit.git. Additional supplementary material is provided in the accompanying supplementary document.

## Competing interests

No competing interest is declared.

## Acknowledgments

This study was financed in part by the Coordenação de Aperfeiçoamento de Pessoal de Nível Superior - Brasil (CAPES); Conselho Nacional de Desenvolvimento Científico e Tecnológico (CNPq) under grant number 444426/2024-8; Fundação de Amparo à Pesquisa do Estado de Minas Gerais (FAPEMIG). The funding agencies had no role in study design, data collection, analysis and interpretation, decision to publish, or preparation of the manuscript.

**Moruf A. Adeagbo** is a PhD candidate in Computer Science at the Universidade Federal de Viçosa, Brazil, and a lecturer at Abiola Ajimobi Technical University, Nigeria. His research interests include bioinformatics, software engineering, machine learning, and artificial intelligence.

**Halimah A. Agbaje** is a PhD student in Animal Biology at the Universidade Federal de Vicosa, Brazil. Her research interest include Animal Breeding and Genetics, Biotechnology, Nutrition and Nutrigenomics, applied statistics, geographical information systems and present fish nutrition with special interest in nutrition and circular biology.

**Sabrina A. Silveira** is a professor of Computer Science at the Universidade Federal de Viçosa, Brazil. Her research interests include bioinformatics, machine learning, data mining, data visualization, and structural bioinformatics, with a focus on computational methods for biological pattern discovery.

## Notes

https://github.com/morufwork/guoAudit.git

### Competing Interest Statement

The authors have declared no competing interest.

