## Supplementary Material for "Protein-novelty, sequence-similarity, and negative-sampling controls reveal benchmark-specific discrimination in protein–protein interaction prediction"

**Title:**

**Guide to supplementary tables and figures**

Supplementary Methods M1–M8 describe the procedures. Supplementary Results S1–S19 present the evidence, with Tables S1–S45 in numerical order and Figures S1–S6 placed with the sections they support. Table and figure identifiers match the citations used in the manuscript. Continued table panels repeat the table number, title, and identifying columns. Threshold metrics use a probability cutoff of 0.5 unless stated otherwise.

Abbreviations. LR, logistic regression; RF, random forest; MLP, multilayer perceptron; AAC, amino-acid composition; CTD, composition/transition/distribution descriptors; ROC-AUC, area under the receiver operating characteristic curve; PR-AUC, average precision; MCC, Matthews correlation coefficient; ECE, expected calibration error; SD, standard deviation; IQR, interquartile range; CI, confidence interval; BH, Benjamini–Hochberg correction; SMD, standardized mean difference; KS, Kolmogorov–Smirnov statistic. “Boot.” and “Jack.” denote connected-component bootstrap and delete-one-component jackknife estimates. A dash denotes an unavailable value.

**Evaluation regimes.** R0 is a random pair split; R1, R2, and R3 contain zero, one, and two unseen test endpoints, respectively. R3-50, R3-40, R3-30, and R3-20 are sequence-cluster-controlled variants generated using MMseqs2 minimum-sequence-identity clustering thresholds of 50%, 40%, 30%, and 20%. N0 denotes original negatives; N1, random unobserved pairs; N2, degree-matched pairs; N3, length-matched pairs; N5, degree–length-matched pairs; N6, two-sided degree substitution; and N7, N5 with within-pair sequence-cluster exclusion. N2-decile, N2-exact, and N2-nearest distinguish degree-matching rules. These labels identify putative negatives, not experimentally established non-interactions.

**Training/evaluation designs.** Design A trains on N0 and replaces the test negatives. Design B trains on each negative set and evaluates on the common N0 test set. Consistent-negative revalidation uses the same negative definition during training and evaluation and is identified separately.

**Supplementary Information**

**M1.** Dataset audit

The primary analysis used the supplied Guo yeast protein sequences and labeled interaction pairs (Guo et al., 2008). Before modeling, we programmatically audited identifiers, sequences, labels, and pair structure for missing identifiers or sequences, residue-alphabet validity, self-pairs, duplicate rows, reversed duplicates, and conflicting labels. We canonicalized each undirected pair to lexical endpoint order before removing duplicates. Identifiers with identical amino-acid sequences were assigned to a common sequence-equivalence group for strict splitting. We characterized the observed positive graph and the originally supplied train/test partition separately.

**M2.** Split construction and sequence-cluster controls

Four protein-overlap regimes were used. R0 was a stratified random pair split and represents the conventional reference in which proteins may recur freely across training and test pairs. R1 contained test pairs for which both proteins occurred somewhere in training, excluding the same pair. R2 contained test pairs with exactly one protein absent from training. R3 contained pairs for which both proteins were absent from training. R1–R3 were created from one partition of exact-sequence-equivalence groups into protein pools and shared a single training set. They therefore constitute the matched protein-novelty curve. A degree-safe holdout constructed R1 without removing the final training occurrence of any protein that was meant to remain “seen.” Mixed-pool pairs formed R2, and pairs with two test-pool proteins formed R3. R0 used an independently sampled stratified 80/20 pair split and is not a matched fourth point in the R1→R2→R3 curve.

Automated gates tested exact-pair overlap, reversed-pair overlap, protein overlap where forbidden, exact-sequence-group overlap, and label integrity. We did not write a split if a required check failed.

We independently generated sequence-cluster-controlled R3 variants at minimum sequence identities of 50%, 40%, 30%, and 20%, denoted R3-50, R3-40, R3-30, and R3-20. All supplied sequences were clustered independently at each threshold before any train/test assignment. The versioned MMseqs2 easy-cluster command used greedy set-cover clustering (--cluster-mode 0), required both query and target to achieve at least 80% alignment coverage (--cov-mode 0 -c 0.8), fixed sensitivity at 4.0, and normalized identity by alignment length (--seq-id-mode 0). Stable representative accessions identified clusters, and we permuted the sorted cluster universe with seed 42. Clusters were assigned to train or test pools; only pairs whose two endpoints fell in the same pool were retained, and cross-pool pairs were discarded.

Clustering and pool assignment were rerun independently at each threshold. Consequently, R3-50, R3-40, R3-30, and R3-20 are non-nested sensitivity analyses rather than progressively filtered versions of a single fixed test set. Differences among these conditions therefore reflect changes in both the operational clustering criterion and the realized train/test composition and should not be interpreted as an isolated dose-response effect of decreasing sequence identity.

We assessed reproducibility by repeated generation and bytewise comparison, alongside exact-protein, exact-sequence, cluster, pair, and reverse-pair overlap checks. These analyses constitute operational sequence-cluster controls under the stated search and coverage parameters; sequence identity thresholds cannot establish the presence or absence of evolutionary homology.

The exact MMseqs2 command template was:

mmseqs easy-cluster INPUT OUTPUT TMP --min-seq-id TAU -c 0.8 --cov-mode 0 --cluster-mode 0 -s 4.0 --seq-id-mode 0 -v 1

MMseqs2 build: 8cc5ce367b5638c4306c2d7cfc652dd099a4643f.

**M3.** Representations, model fitting, and sensitivity procedures

We represented each protein independently of its interaction partner before pair construction. The principal handcrafted representation concatenated 20-dimensional amino-acid composition (AAC) and 147-dimensional composition/transition/distribution (CTD) descriptors. We evaluated eight Biopython ProtParam physicochemical summaries separately in the conventional five-fold baseline and did not include them in AAC+CTD. We computed features once per unique sequence and cached them. The principal pretrained representation was the frozen ESM-2 checkpoint facebook/esm2_t12_35M_UR50D, which produces 480-dimensional residue embeddings. We evaluated mean pooling and CLS-token pooling because they provide two common, reproducible reductions from residue-level to protein-level vectors. No PLM weights were fine-tuned.

We capped sequences at 1,022 residues for computational tractability. For sequences exceeding this length, the first 1,022 residues were retained, corresponding to deterministic C-terminal truncation. We logged truncation per protein and ran a dedicated sensitivity analysis to assess its influence. We included a frozen, mean-pooled Rostlab/prot_t5_base_mt_uniref50 representation only as a limited cross-check; we did not treat it as an exhaustive test of ProtT5 or PLMs.

Since PPIs are undirected, primary pair features were symmetric by construction. For protein embeddings hA and hB, evaluated operators included hA+hB, |hA−hB|, the Hadamard product hA⊙hB, and their concatenated combination. We retained plain concatenation only as an explicitly order-sensitive comparator. We also tested a learned symmetric operator. We assessed pair-order invariance with automated endpoint-swap tests.

Baseline models included logistic regression, random forest, XGBoost, and a multilayer perceptron (MLP). Logistic regression used L2 regularization with C=1 and a maximum of 2,000 iterations. The main sequence-baseline and architecture random forests used 300 trees; topology/length controls used 200 trees with a maximum depth of 6. XGBoost used 300 estimators and log-loss evaluation. The MLP used hidden layers of 128 and 64 units, ReLU activation, Adam optimization, a maximum of 500 iterations, and early stopping. We standardized classical pair features using the training pairs; we standardized neural architecture inputs using the unique proteins in the training pool. We selected internal validation subsets afterward, so these transformations did not use external test data and were not refit on the smaller internal fitting subset. No class weighting was used. We report class counts for each realized dataset; exact-matching omissions can introduce imbalance, particularly in the species-specific analyses.

Architecture experiments added a shared-encoder Siamese network, a single-head attention-fusion model, and a low-rank cross-interaction module. The Siamese encoder mapped each protein through a 256-unit hidden layer to a 128-dimensional latent vector, combined the two latent vectors symmetrically, and used a 64-unit classifier head. Attention fusion projected the two protein tokens to a 128-dimensional query/key/value space before symmetric aggregation. Cross-interaction combined the elementwise product of shared 128-dimensional projections with the sum and absolute difference of the original protein embeddings, followed by 128- and 64-unit classifier layers. Neural architectures used Adam with learning rate 10−3, batch size 128, up to 100 epochs, and early stopping with patience 10 on a 15% validation subset drawn from training only. We did not perform per-condition hyperparameter searches; the reported comparisons used fixed settings rather than condition-specific optimization.

AAC comprises the frequencies of the 20 standard amino acids; CTD comprises 147 composition, transition, and distribution descriptors across physicochemical residue groupings. The eight separately evaluated ProtParam summaries were molecular weight, aromaticity, instability index, isoelectric point, GRAVY, and predicted helix, turn, and sheet fractions. Truncation sensitivity compared N-terminal retention, C-terminal retention, full-sequence processing, and averaging of non-overlapping 1,022-residue segments, both across all test pairs and within affected/unaffected subsets. The conventional baseline used stratified pair-level five-fold cross-validation (seed 42). Five model-training seeds held the split and negative realization fixed. The sklearn MLP used its internal early-stopping validation defaults; the three neural pair architectures used the 15% pair-level validation procedure described above.

**M4.** Negative construction and topology diagnostics

N0 denotes the benchmark's original constructed negatives. We generated alternative sets to expose properties N0 might encode: N1 comprised sampled unobserved pairs randomly; N2 matched positive-network degree; N3 matched sequence length; and N5 matched both degree and length. N6 permitted two-sided degree-matched substitution, and N7 added within-pair sequence-cluster exclusion to N5's joint degree–length matching, using the saved 20%-identity cluster assignment. This within-pair cluster constraint is distinct from the versioned R3-20 train/test split control. We did not create a localization-aware set because Guo contains no localization, Gene Ontology, or complex-membership annotations. Throughout, we refer to these examples as constructed, putative, or unobserved negative pairs and do not interpret them as experimentally established biological non-interactions.

We created N2, N3, and N5 by one-sided matched endpoint substitution. For each positive pair, we selected one endpoint by a reproducible random draw, while the other remained fixed. Candidate replacements were drawn from the replaced protein's degree decile, length decile, or joint degree–length stratum. We rejected candidates if they produced a self-pair, recreated a known positive in either order, or duplicated a previously constructed negative pair. If no candidate met all requirements, we omitted the proposed negative instead of relaxing the matching rule; we retained the positive pair. Proteins could recur across constructed pairs, but resulting pairs were unique.

For R3, we constructed negatives after protein-pool assignment. We sampled train and test negatives independently from their respective pools, and rechecked protein disjointness after substitution. For R0, a full negative set was generated before the pair-level split because protein disjointness was not required. Within any comparison, every model used the same realized negative set.

We evaluated negative-set effects using complementary designs. In the distribution-shift experiment, we partitioned positive pairs once into 80% training and 20% test subsets (seed 42), and partitioned each negative set separately using the same proportions and seed. We evaluated an ESM-2–MLP trained on N0 after replacing test negatives with N1, N2, N3, or N5; in the reverse direction, we evaluated models trained on each negative set on the common N0 test set. We shared positive partitions across these comparisons. These designs assess sensitivity to the training or evaluation negative definition. Separate R0/R3 revalidation used the same negative definition in both training and evaluation.

Positive-network degree di was defined as the number of positive partners observed for protein i in the deduplicated Guo positive graph, including zero for proteins without an observed positive edge. The graph's mean degree, the mean across distinct proteins, and pair-associated endpoint degree are different quantities. Pair-associated degree pools endpoint occurrences and therefore upweights frequently occurring proteins. This explains why positive-pair endpoint degree can exceed the ordinary graph mean.

Topology-only classifiers used the sum, absolute difference, minimum, and maximum of endpoint positive-network degrees, without sequence information. Length-only controls used the same summaries of sequence length. These controls tested how well a label-derived graph property or a simple sequence attribute separated the benchmark labels. Full-graph degree is a retrospective audit feature, not a prospective input for a protein with no known interaction graph. It cannot isolate degree from other properties that may covary with the observed network.

Robustness analyses regenerated N2 under 30 data-generation seeds while fixing model-training seed and split, distinguishing sensitivity to decoy selection from sensitivity to optimization. Degree-source analyses compared the full-positive graph and training-only degree where defined. Exact-degree and nearest-degree matching tested whether decile coarseness explained residual results. One- versus two-sided substitution and sequence-cluster-excluded N7 provided further sensitivity checks.

Exact-degree substitution required equality of observed degree; nearest-degree substitution selected eligible candidates with the smallest absolute degree difference. Failed substitutions omitted the proposed negative while retaining the positive. Balance diagnostics included endpoint-occurrence SMD, KS distance, and Wasserstein distance. Endpoint occurrences are dependent; their distributional statistics are descriptive. The distinct-protein mean-difference diagnostic used one observation per protein and a separate 2,000-replicate bootstrap, rather than the model-level component bootstrap. N7 used the saved 20%-identity cluster assignment for within-pair rejection, separately from the regenerated R3-20 split audit.

**M5.** Metrics, score diagnostics, and calibration

Primary discrimination metrics were ROC-AUC and PR-AUC, with PR-AUC calculated as average precision; we also recorded Matthews correlation coefficient (MCC), balanced accuracy, precision, recall, specificity, F1, and accuracy at a fixed probability threshold of 0.5. Test thresholds were never optimized. The fixed cutoff of 0.5 provided a common reporting convention, while calibrated and precision–recall operating-point sensitivity analyses assessed whether conclusions depended on raw probability scaling. We saved every prediction at the sample level.

We calibrated probabilities using Platt and isotonic mappings fitted exclusively on validation data carved from training. We assessed raw and calibrated probabilities using Brier score, expected calibration error, and calibration slope/intercept. Calibration can change probability reliability and threshold performance. Isotonic mappings introduce ties, and an unconstrained fitted sigmoid can reverse score ordering; neither change establishes additional interaction information. Calibration models reserved 20% of the R3 training pairs for fitting the mappings and were therefore fitted separately from the architecture comparison. We compare raw and calibrated metrics within the same fitted model. Precision–recall operating points are descriptive summaries of the saved model test curves at fixed targets of 0.5; they are not prospectively validated thresholds.

Below-chance results were examined using the saved probability p and its complement 1−p. For each principal R0/R3 evaluation under N0, N2, and N5, we calculated AUC(p), AUC(1−p), positive- and negative-class means, medians, interquartile ranges, and Cohen's d. Because AUC(1−p)=1−AUC(p), inversion is an algebraic diagnostic rather than independent evidence. Material class-conditional reversal was distinguished from fluctuations close to 0.5.

We evaluated ECE with equal-width and equal-frequency bins at 5, 10, 15, and 20 bins; the primary calibration comparison used 10 equal-width bins. We fitted the calibration slope and intercept against the logit of the predicted probability. Precision at recall 0.5 and recall at precision 0.5 summarize the observed precision–recall curves without selecting a prospective decision rule.

**M6.** Graph-aware inference

PPI pairs sharing an endpoint may be dependent. To accommodate dependence extending through connected pairs, the analysis grouped test pairs by connected component. Confidence procedures therefore used connected components of the undirected test-pair graph as resampling units rather than treating pairs as independent or clustering on only one endpoint column. A percentile connected-component bootstrap resampled whole components with replacement for 2,000 replicates. A delete-one-component jackknife provided a small-component-count sensitivity analysis with a t reference distribution based on the effective number of components. We used paired differences when models shared an identical test set; for N0–N2 comparisons, we resampled the two component distributions separately because their test-pair sets differed. This unpaired procedure is a working approximation: the sets share positive pairs and originate from the same biological graph, so their statistical independence is not established.

The defined inferential family contained chance tests for ROC-AUC, MCC, and PR-AUC across six model configurations and two negative sets, paired architecture-versus-baseline comparisons, and unpaired N0–N2 comparisons. We adjusted two-sided p-values using Benjamini–Hochberg within each method's complete 44-test family. We treated bootstrap and jackknife as alternative inferential methods and corrected them separately. Giant-component exclusion and leave-one-component-out analyses assessed sensitivity to component composition.

We distinguish three evidentiary levels. A descriptive result reports performance on the observed fixed split. Algorithmic replication shows that a direction persists across model configurations or random seeds but does not add independent biological units. Formal inference refers only to the corrected component-aware tests. Terms such as “large,” “consistent,” “near chance,” and “not statistically distinguishable from chance” are used according to these separate meanings.

The null values were 0.5 for ROC-AUC, 0 for MCC, and the realized positive prevalence for average precision. The 44-test family comprised 36 metric-versus-null tests (three metrics × six model configurations × two negative sets), two paired Siamese-versus-logistic-regression ROC-AUC contrasts, and six N0–N2 ROC-AUC contrasts. Degree-only controls were outside this family. “Above chance” required a positive point difference from the relevant null and a two-sided BH-adjusted P value below 0.05.

**M7.** PIPR multispecies and within-species benchmark procedures

The secondary analysis used the PIPR multispecies benchmark comprising C. elegans, D. melanogaster, and E. coli (Chen et al., 2019). We used this benchmark to assess whether the degree-associated pattern and the response of sequence-based models to degree matching observed in Guo also occurred in a separate dataset. Protein accessions were first resolved taxonomically, after which pooled and within-species negative sets were reconstructed. For species-specific analyses, we recomputed positive-network degrees and matching strata independently within each species, and restricted replacement endpoints to proteins from the same species. Each species-specific replication used an independently reconstructed 80/20 random pair split (seed 42). In the pooled analysis, we used a same-species indicator to assess the contribution of taxonomic pairing to discrimination.

We resolved taxonomy from cached UniProtKB accession records and NCBI protein ESummary records for legacy GI identifiers. We checked retired UniProt accessions against UniSave version 1; we used archived organism names when those early records lacked a numerical taxon identifier. Unresolved labels were assigned by exact sequence identity only when all matching resolved benchmark records belonged to one species. E. coli strain records were consolidated. We excluded proteins from other organisms, unresolved endpoints, and cross-species pairs from the three species-specific datasets; we retained the original combined results.

Within each species, we partitioned the existing canonicalized, self-pair-filtered input without changing its sequences or original labels. Degrees were computed from that species’s full positive graph, including zero-degree proteins in the candidate universe. Negative sets used the original N0 labels or the same one-sided decile, exact-degree, and nearest-degree substitution rules as the pooled analysis. We rejected self-pairs, known positives, and duplicate negatives. For unmatchable substitutions, we omitted the proposed negative while retaining its positive pair. We split positive pairs once (80/20; random_state=42); we split each negative set independently with the same settings. We shared the positive split across variants, and all five models used identical realized pairs within each variant. We used N2 negatives consistently for both training and testing.

AAC+CTD features and frozen ESM-2 mean embeddings were reused from the combined analysis. Pair fusion concatenated the sum, absolute difference, and element-wise product. We fitted StandardScaler on training data only. Sequence logistic regression used max_iter=2000; the MLP used hidden layers (128,64), max_iter=500, and early_stopping=True, with the existing sklearn validation defaults. Degree-only logistic regression used the same settings as the pooled analysis. All estimators used random_state=42. ROC-AUC, average precision (reported as PR-AUC), MCC, F1, balanced accuracy, accuracy, precision, recall, and specificity were saved; threshold-based metrics used 0.5. No per-species hyperparameter tuning was performed.

Degree SMD, Kolmogorov–Smirnov (KS) distance, and Wasserstein distance describe pooled endpoint occurrences; they are not independent-endpoint significance tests. Distinct-protein balance intervals used 2,000 bootstrap replicates and were interpreted separately from model-level inference. This single-split replication included no cluster-aware model comparisons or multi-seed analyses. Full-graph degree is a retrospective benchmark diagnostic. Random pair splits permit protein and sequence recurrence, so residual discrimination cannot be attributed exclusively to molecular compatibility.

The same-species diagnostic assigned score 1 to endpoints from the same target species and 0 otherwise, without fitting a classifier. It used the reconstructed pooled N0/N2 test partitions and retained only pairs with both endpoints resolved to the three target species.

**M8.** Reproducibility and provenance

The repository contains numbered scripts, configurations, immutable input data, generated split files, sample-level predictions, tables, figures, environment reports, and automated checks.

**Repository:** <https://github.com/morufwork/guoAudit.git>

Principal regeneration order: environment setup; dataset audit/canonicalization; protein and MMseqs2 split generation; handcrafted and PLM features; baseline and strict-split models; negative-set analyses; architecture and statistical analyses; manuscript outputs. The split report records the exact MMseqs2 command and build.

Tables and figures derive from saved analysis outputs. The accompanying reproducibility materials contain the table-to-source mapping, analysis scripts, sample-level predictions, and environment records. Supplementary Figure S1 summarizes how M1–M8 connect dataset audit, split construction, representations/models, negative-set controls, metrics/calibration, graph-aware inference, and the secondary PIPR analysis.

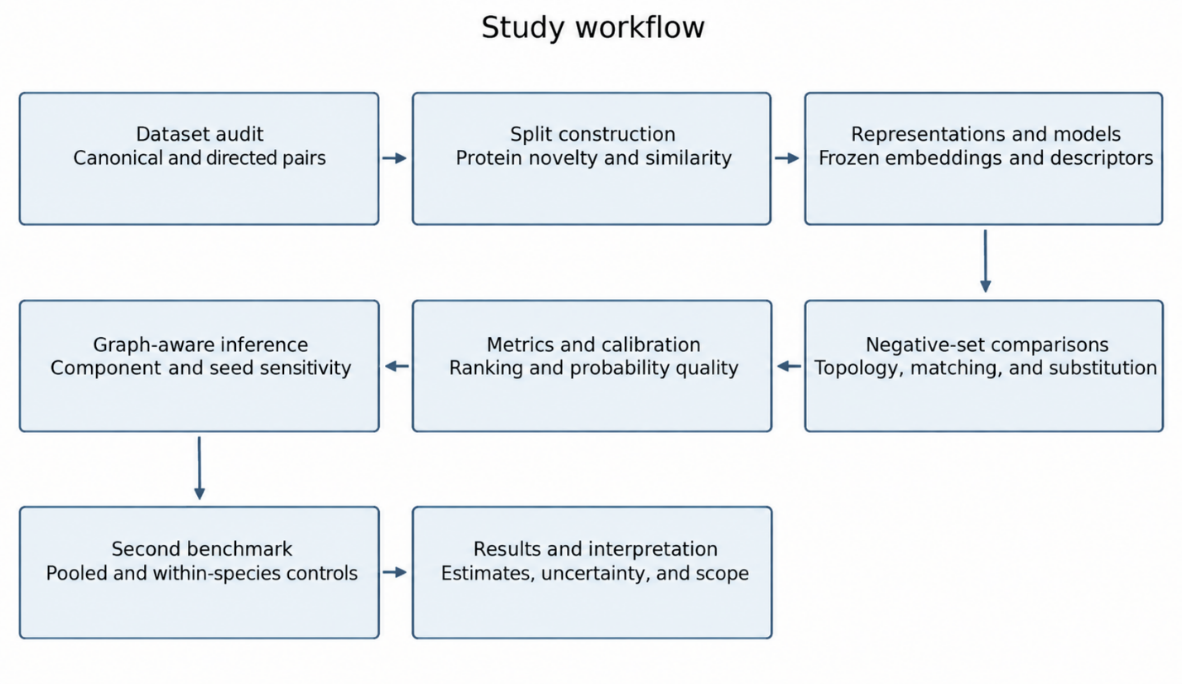

**Supplementary Figure S1. Study workflow.** Dataset audit and canonicalization precede split construction, representation/model evaluation, negative-set controls, graph-aware inference, and secondary-benchmark analysis. The boxes map directly to Supplementary Methods M1–M8 and the corresponding main-text Methods sections.

**Supplementary Results**

**S1 Dataset integrity**

Canonicalization removed 24 duplicate pairs, leaving 5,581 positives and 5,583 constructed negatives. Eight identifiers formed three identical-sequence groups. The positive graph contained 98 components, with 1,631 proteins in the largest component.

**Supplementary Table S1.** Guo yeast dataset integrity and positive-network characteristics

| **Characteristic** | **Value** |
| --- | --- |
| Unique proteins (raw) | 2,497 |
| PPI pairs (raw) | 11,188 |
| Positive pairs (raw) | 5,594 |
| Negative pairs (raw, N0) | 5,594 |
| Canonical (deduplicated) pairs | 11,164 |
| Exact duplicate pairs removed | 24 |
| Sequence-identity equivalence groups | 2,492 |
| Non-singleton identical-sequence groups | 3 |
| Positive-interaction graph: nodes | 2,217 |
| Positive-interaction graph: edges | 5,581 |
| Positive-interaction graph: density | 0.0023 |
| Positive-interaction graph: mean degree | 5.0300 |
| Positive-interaction graph: max degree | 15 |
| Positive-interaction graph: connected components | 98 |
| Positive-interaction graph: largest component size | 1,631 |
| Clustering coefficient | 0.0031 |
| Proteins belonging to a non-singleton R3-20 MMseqs2 cluster (fraction) | 0.2022 |

Counts refer to the supplied dataset unless marked canonical. Positive-network statistics use the deduplicated positive graph; sequence-identity groups include singleton sequences.

**S2 Split composition and integrity**

R1–R3 shared 5,703 training pairs and contained 1,426, 3,570, and 462 test pairs. All regenerated MMseqs2 variants passed the required overlap checks and were byte-identical on repeated generation. Table S2 summarizes the matched R0–R3 protein-novelty splits and their sequence-similarity characterization; Table S3 reports the regenerated sequence-cluster-controlled R3 variants used in the current comparisons.

**Supplementary Table S2.** Protein-novelty split composition and sequence-similarity characterization. Panel A (1 of 4)

| **Row** | **Regime** | **Train pairs** | **Test pairs** | **Train proteins** | **Test proteins** |
| --- | --- | --- | --- | --- | --- |
| 1 | R0 | 8,931 | 2,233 | 2,497 | 2,080 |
| 2 | R1 | 5,703 | 1,426 | 1,997 | 1,561 |
| 3 | R2 | 5,703 | 3,570 | 1,997 | 2,164 |
| 4 | R3 | 5,703 | 462 | 1,997 | 420 |

**Supplementary Table S2 (continued).** Protein-novelty split composition and sequence-similarity characterization. Panel B (2 of 4)

| **Row** | **Regime** | **Train positives** | **Train negatives** | **Test positives** | **Test negatives** |
| --- | --- | --- | --- | --- | --- |
| 1 | R0 | 4,465 | 4,466 | 1,116 | 1,117 |
| 2 | R1 | 2,869 | 2,834 | 740 | 686 |
| 3 | R2 | 2,869 | 2,834 | 1,746 | 1,824 |
| 4 | R3 | 2,869 | 2,834 | 223 | 239 |

**Supplementary Table S2 (continued).** Protein-novelty split composition and sequence-similarity characterization. Panel C (3 of 4)

| **Row** | **Regime** | **Test components** | **Largest test component (proteins)** | **Test mean degree** | **Test median degree** |
| --- | --- | --- | --- | --- | --- |
| 1 | R0 | 97 | 1,818 | 4.7000 | 5 |
| 2 | R1 | 178 | 1,034 | 4.7300 | 5 |
| 3 | R2 | 2 | 2,162 | 4.5800 | 5 |
| 4 | R3 | 19 | 377 | 4.6000 | 5 |

**Supplementary Table S2 (continued).** Protein-novelty split composition and sequence-similarity characterization. Panel D (4 of 4)

| **Row** | **Regime** | **Test proteins with training match at 50% identity (%)** | **Test proteins with training match at 40% identity (%)** | **Test proteins with training match at 30% identity (%)** | **Test proteins with training match at 20% identity (%)** |
| --- | --- | --- | --- | --- | --- |
| 1 | R0 | 100 | 100 | 100 | 100 |
| 2 | R1 | 100 | 100 | 100 | 100 |
| 3 | R2 | 78.9000 | 79.9000 | 80.6000 | 81.3000 |
| 4 | R3 | 8.6000 | 13.1000 | 16.2000 | 18.8000 |

R1–R3 share one training set. Training-match percentages describe sequence similarity at the stated identity threshold and do not establish evolutionary homology.

**Supplementary Table S3.** Integrity of the regenerated MMseqs2 sequence-cluster-controlled splits. Panel A (1 of 3)

| **Row** | **Regime** | **Train pairs** | **Test pairs** | **Train proteins** | **Test proteins** |
| --- | --- | --- | --- | --- | --- |
| 1 | R3-50 | 7,227 | 450 | 2,004 | 407 |
| 2 | R3-40 | 7,111 | 423 | 1,998 | 391 |
| 3 | R3-30 | 7,174 | 419 | 2,007 | 394 |
| 4 | R3-20 | 7,043 | 445 | 1,996 | 420 |

**Supplementary Table S3 (continued).** Integrity of the regenerated MMseqs2 sequence-cluster-controlled splits. Panel B (2 of 3)

| **Row** | **Regime** | **Exact protein overlap** | **Exact sequence overlap** | **Cluster overlap** | **Pair/reversal overlap** |
| --- | --- | --- | --- | --- | --- |
| 1 | R3-50 | 0.0000 | 0.0000 | 0.0000 | 0.0000 |
| 2 | R3-40 | 0.0000 | 0.0000 | 0.0000 | 0.0000 |
| 3 | R3-30 | 0.0000 | 0.0000 | 0.0000 | 0.0000 |
| 4 | R3-20 | 0.0000 | 0.0000 | 0.0000 | 0.0000 |

**Supplementary Table S3 (continued).** Integrity of the regenerated MMseqs2 sequence-cluster-controlled splits. Panel C (3 of 3)

| **Row** | **Regime** | **Identity threshold (proportion)** | **Query coverage (proportion)** | **Target coverage (proportion)** | **All checks passed** |
| --- | --- | --- | --- | --- | --- |
| 1 | R3-50 | 0.5000 | 0.8000 | 0.8000 | Yes |
| 2 | R3-40 | 0.4000 | 0.8000 | 0.8000 | Yes |
| 3 | R3-30 | 0.3000 | 0.8000 | 0.8000 | Yes |
| 4 | R3-20 | 0.2000 | 0.8000 | 0.8000 | Yes |

Identity and query/target coverage thresholds are proportions. Overlap counts compare training with test data. “Yes” indicates that every required integrity check passed.

**S3 Post-audit yeast distributions**

All 2,497 protein identifiers were retained after pair deduplication. Of these, 280 had no observed positive partner and 268 exceeded 1,022 residues.

**Supplementary Table S4.** Post-audit Guo class counts, protein counts, and sequence lengths

| **Quantity** | **Value** |
| --- | --- |
| Unique pairs | 11,164 |
| Positive pairs | 5,581 |
| Constructed negative pairs | 5,583 |
| Protein identifiers | 2,497 |
| Observed-positive-graph nodes | 2,217 |
| Zero-positive-degree proteins | 280 |
| Sequences >1,022 residues | 268 |
| Minimum sequence length | 55 |
| Median sequence length | 461 |
| Maximum sequence length | 4,910 |

Sequence lengths are in residues. Zero observed positive degree does not establish biological isolation.

**S4 Protein-novelty and representation comparisons**

AAC+CTD–MLP ROC-AUC decreased from 0.697 at R1 to 0.626 at R2 and 0.587 at R3. Independently generated sequence-cluster-controlled variants showed no monotonic trend. ESM-2 CLS and mean pooling exchanged relative advantage across regimes. Supplementary Table S5 provides the complete AAC+CTD metrics underlying main-text Figure 3, whereas Supplementary Table S6 provides the representation comparison visualized in Supplementary Figure S2.

**Supplementary Table S5.** AAC+CTD baseline-model performance across evaluation regimes (main-text Figure 3)

| **Regime** | **Model** | **Train pairs** | **Test pairs** | **ROC-AUC** | **PR-AUC** | **MCC** |
| --- | --- | --- | --- | --- | --- | --- |
| R0 | LR | 8,931 | 2,233 | 0.6399 | 0.6149 | 0.2086 |
| R0 | RF | 8,931 | 2,233 | 0.6845 | 0.6548 | 0.2920 |
| R0 | XGBoost | 8,931 | 2,233 | 0.6756 | 0.6506 | 0.2523 |
| R0 | MLP | 8,931 | 2,233 | 0.7010 | 0.6801 | 0.3096 |
| R1 | LR | 5,703 | 1,426 | 0.6600 | 0.6530 | 0.2518 |
| R1 | RF | 5,703 | 1,426 | 0.7007 | 0.6850 | 0.3206 |
| R1 | XGBoost | 5,703 | 1,426 | 0.6954 | 0.6877 | 0.2788 |
| R1 | MLP | 5,703 | 1,426 | 0.6966 | 0.6837 | 0.3050 |
| R2 | LR | 5,703 | 3,570 | 0.6060 | 0.5686 | 0.1503 |
| R2 | RF | 5,703 | 3,570 | 0.6516 | 0.6026 | 0.2183 |
| R2 | XGBoost | 5,703 | 3,570 | 0.6409 | 0.5945 | 0.1988 |
| R2 | MLP | 5,703 | 3,570 | 0.6263 | 0.5873 | 0.1908 |
| R3 | LR | 5,703 | 462 | 0.5581 | 0.5228 | 0.1086 |
| R3 | RF | 5,703 | 462 | 0.5867 | 0.5230 | 0.1356 |
| R3 | XGBoost | 5,703 | 462 | 0.6331 | 0.5970 | 0.1983 |
| R3 | MLP | 5,703 | 462 | 0.5874 | 0.5491 | 0.1291 |
| R3-50 | LR | 7,227 | 450 | 0.6084 | 0.5836 | 0.1855 |
| R3-50 | RF | 7,227 | 450 | 0.6298 | 0.6031 | 0.1495 |
| R3-50 | XGBoost | 7,227 | 450 | 0.5831 | 0.5681 | 0.1297 |
| R3-50 | MLP | 7,227 | 450 | 0.5838 | 0.5669 | 0.0872 |
| R3-40 | LR | 7,111 | 423 | 0.6184 | 0.5378 | 0.1816 |
| R3-40 | RF | 7,111 | 423 | 0.6409 | 0.6184 | 0.1994 |
| R3-40 | XGBoost | 7,111 | 423 | 0.5868 | 0.5224 | 0.1045 |
| R3-40 | MLP | 7,111 | 423 | 0.5860 | 0.5308 | 0.1365 |
| R3-30 | LR | 7,174 | 419 | 0.5564 | 0.5287 | 0.1306 |
| R3-30 | RF | 7,174 | 419 | 0.5758 | 0.5756 | 0.1014 |
| R3-30 | XGBoost | 7,174 | 419 | 0.5540 | 0.5445 | 0.1121 |
| R3-30 | MLP | 7,174 | 419 | 0.5443 | 0.5453 | 0.0982 |
| R3-20 | LR | 7,043 | 445 | 0.6102 | 0.5446 | 0.1627 |
| R3-20 | RF | 7,043 | 445 | 0.6600 | 0.5914 | 0.2337 |
| R3-20 | XGBoost | 7,043 | 445 | 0.6249 | 0.5496 | 0.2169 |
| R3-20 | MLP | 7,043 | 445 | 0.5856 | 0.5061 | 0.0962 |

R1–R3 share a training set; R0 is an independent reference. Sequence-cluster-controlled variants are generated independently at each clustering threshold.

**Supplementary Table S6.** Representation performance with a shared MLP classifier (Supplementary Figure S2)

| **Regime** | **Representation** | **Train pairs** | **Test pairs** | **ROC-AUC** | **PR-AUC** | **MCC** |
| --- | --- | --- | --- | --- | --- | --- |
| R0 | AAC+CTD | 8,931 | 2,233 | 0.7010 | 0.6801 | 0.3096 |
| R0 | ESM-2 mean | 8,931 | 2,233 | 0.7305 | 0.7162 | 0.3561 |
| R0 | ESM-2 CLS | 8,931 | 2,233 | 0.7555 | 0.7493 | 0.3839 |
| R1 | AAC+CTD | 5,703 | 1,426 | 0.6966 | 0.6837 | 0.3050 |
| R1 | ESM-2 mean | 5,703 | 1,426 | 0.7258 | 0.7169 | 0.3414 |
| R1 | ESM-2 CLS | 5,703 | 1,426 | 0.7324 | 0.7342 | 0.3443 |
| R2 | AAC+CTD | 5,703 | 3,570 | 0.6263 | 0.5873 | 0.1908 |
| R2 | ESM-2 mean | 5,703 | 3,570 | 0.6700 | 0.6397 | 0.2395 |
| R2 | ESM-2 CLS | 5,703 | 3,570 | 0.6668 | 0.6359 | 0.2366 |
| R3 | AAC+CTD | 5,703 | 462 | 0.5874 | 0.5491 | 0.1291 |
| R3 | ESM-2 mean | 5,703 | 462 | 0.6168 | 0.5726 | 0.1488 |
| R3 | ESM-2 CLS | 5,703 | 462 | 0.5624 | 0.5467 | 0.1122 |
| R3-50 | AAC+CTD | 7,227 | 450 | 0.5838 | 0.5669 | 0.0872 |
| R3-50 | ESM-2 mean | 7,227 | 450 | 0.5730 | 0.5904 | 0.1005 |
| R3-50 | ESM-2 CLS | 7,227 | 450 | 0.6134 | 0.6114 | 0.1731 |
| R3-40 | AAC+CTD | 7,111 | 423 | 0.5860 | 0.5308 | 0.1365 |
| R3-40 | ESM-2 mean | 7,111 | 423 | 0.6497 | 0.6217 | 0.1868 |
| R3-40 | ESM-2 CLS | 7,111 | 423 | 0.6049 | 0.5517 | 0.1399 |
| R3-30 | AAC+CTD | 7,174 | 419 | 0.5443 | 0.5453 | 0.0982 |
| R3-30 | ESM-2 mean | 7,174 | 419 | 0.6292 | 0.6229 | 0.1835 |
| R3-30 | ESM-2 CLS | 7,174 | 419 | 0.6462 | 0.6759 | 0.2140 |
| R3-20 | AAC+CTD | 7,043 | 445 | 0.5856 | 0.5061 | 0.0962 |
| R3-20 | ESM-2 mean | 7,043 | 445 | 0.6595 | 0.6037 | 0.1730 |
| R3-20 | ESM-2 CLS | 7,043 | 445 | 0.6644 | 0.5868 | 0.2187 |

AAC+CTD, ESM-2 mean, and ESM-2 CLS use the same downstream MLP and saved evaluation splits. Supplementary Figure S2 visualizes the regime-dependent representation comparison in Table S6; it does not imply an overall ranking of handcrafted descriptors versus protein language models.

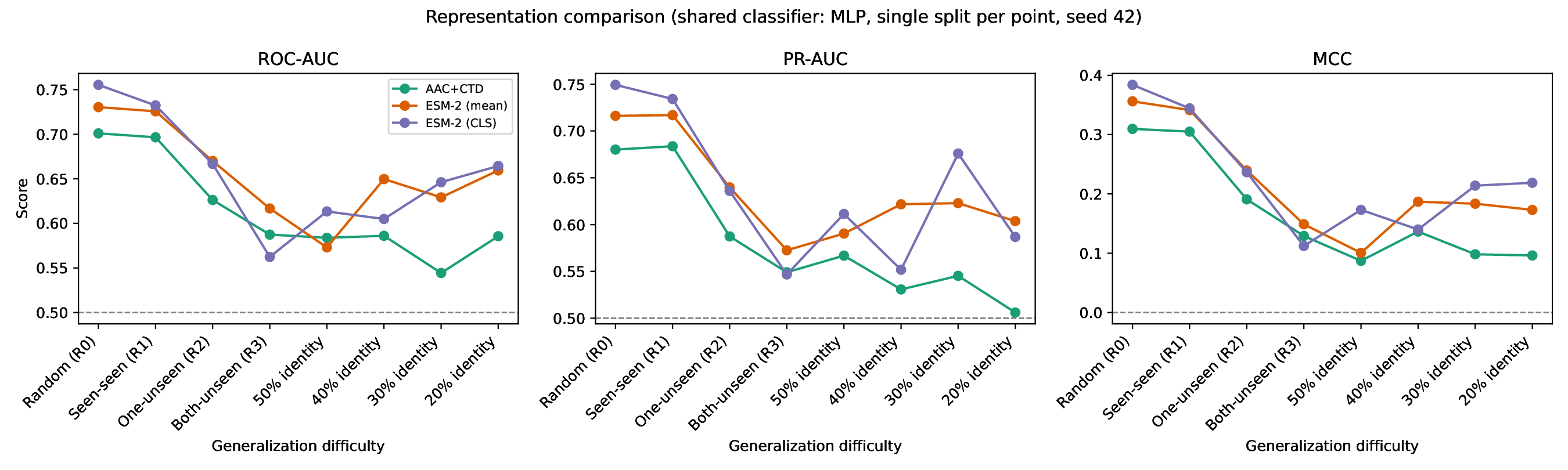

**Supplementary Figure S2.** Representation performance across evaluation regimes. AAC+CTD, ESM-2 mean, and ESM-2 CLS use the same downstream MLP and saved splits. Relative advantages depend on the evaluation regime; the complete values are reported in Supplementary Table S6

**S5 Guo degree imbalance and matching controls**

Mean positive-network degree was 6.72 across positive endpoint occurrences and 3.34 across N0 endpoint occurrences (SMD 1.160). The N0-trained degree-only logistic model yielded ROC-AUC 0.502 under decile matching and 0.499 under exact and nearest matching. The distributional pattern is shown in the main Figure 4.

**Supplementary Table S7.** Endpoint-occurrence-weighted degree distributions in Guo positive and negative sets

| **Pair set** | **Endpoint occurrences** | **Mean** | **Median** | **SD** | **Q1** | **Q3** | **IQR** |
| --- | --- | --- | --- | --- | --- | --- | --- |
| Positive | 11,162 | 6.7244 | 7 | 2.7273 | 5 | 9 | 4 |
| N0 | 11,166 | 3.3418 | 2 | 3.0915 | 1 | 6 | 5 |
| N1 | 11,162 | 4.4517 | 4 | 3.1924 | 2 | 7 | 5 |
| N2 | 11,162 | 6.6952 | 7 | 2.7277 | 5 | 9 | 4 |
| N3 | 11,162 | 5.5727 | 6 | 3.1653 | 3 | 8 | 5 |
| N5 | 11,162 | 6.6833 | 7 | 2.6978 | 5 | 9 | 4 |

Each pair contributes two endpoint occurrences. Degree is the number of observed positive partners. Q1 and Q3 are the 25th and 75th percentiles; IQR = Q3 − Q1.

**Supplementary Table S8.** Degree-distribution differences between Guo positive and negative sets. Panel A (1 of 4).

| **Row** | **Comparison** | **Mean difference** | **Median difference** | **Cohen’s d** | **SMD** |
| --- | --- | --- | --- | --- | --- |
| 1 | Positive − N0 | 3.3827 | 5 | 1.1604 | 1.1604 |
| 2 | Positive − N1 | 2.2727 | 3 | 0.7655 | 0.7655 |
| 3 | Positive − N2 | 0.0292 | 0.0000 | 0.0107 | 0.0107 |
| 4 | Positive − N3 | 1.1517 | 1 | 0.3898 | 0.3898 |
| 5 | Positive − N5 | 0.0411 | 0.0000 | 0.0152 | 0.0152 |

**Supplementary Table S8 (continued).** Degree-distribution differences between Guo positive and negative sets. Panel B (2 of 4)

| **Row** | **Comparison** | **\|SMD\| < 0.10** | **\|SMD\| < 0.25** | **KS statistic** | **KS P value** |
| --- | --- | --- | --- | --- | --- |
| 1 | Positive − N0 | No | No | 0.4798 | 0.0000 |
| 2 | Positive − N1 | No | No | 0.3049 | 0.0000 |
| 3 | Positive − N2 | Yes | Yes | 0.0049 | 0.9992 |
| 4 | Positive − N3 | No | No | 0.1549 | 2.08e-117 |
| 5 | Positive − N5 | Yes | Yes | 0.0070 | 0.9463 |

**Supplementary Table S8 (continued).** Degree-distribution differences between Guo positive and negative sets. Panel C (3 of 4)

| **Row** | **Comparison** | **Wasserstein distance** | **Distinct-protein mean difference** | **Distinct-protein 95% CI lower** | **Distinct-protein 95% CI upper** |
| --- | --- | --- | --- | --- | --- |
| 1 | Positive − N0 | 3.3827 | 0.6118 | 0.4328 | 0.7858 |
| 2 | Positive − N1 | 2.2727 | 0.5711 | 0.3920 | 0.7526 |
| 3 | Positive − N2 | 0.0306 | -0.1945 | -0.3663 | -0.0274 |
| 4 | Positive − N3 | 1.1517 | 0.4910 | 0.3269 | 0.6612 |
| 5 | Positive − N5 | 0.0411 | -0.1802 | -0.3508 | -0.0014 |

**Supplementary Table S8 (continued).** Degree-distribution differences between Guo positive and negative sets. Panel D (4 of 4)

| **Row** | **Comparison** | **Distinct-protein CI excludes 0** | **Positive-set proteins** | **Negative-set proteins** |
| --- | --- | --- | --- | --- |
| 1 | Positive − N0 | Yes | 2,217 | 2,421 |
| 2 | Positive − N1 | Yes | 2,217 | 2,459 |
| 3 | Positive − N2 | Yes | 2,217 | 2,103 |
| 4 | Positive − N3 | Yes | 2,217 | 2,444 |
| 5 | Positive − N5 | Yes | 2,217 | 2,108 |

Differences are positive minus negative. Endpoint-occurrence statistics are descriptive; the distinct-protein mean difference and its 95% bootstrap interval use one observation per protein. KS P values do not account for dependence between endpoint occurrences.

**Supplementary Table S9.** Guo degree-only model performance under decile, exact, and nearest-degree matching. Panel A (1 of 3)

| **Row** | **Model** | **Training set** | **Test set** | **Generated negatives** | **ROC-AUC** | **PR-AUC** | **F1** |
| --- | --- | --- | --- | --- | --- | --- | --- |
| 1 | LR | N0 | N0 | 5,583 | 0.9624 | 0.9650 | 0.8881 |
| 2 | LR | N0 | N2-decile | 5,581 | 0.5025 | 0.5055 | 0.6399 |
| 3 | LR | N0 | N2-exact | 5,581 | 0.4994 | 0.5000 | 0.6412 |
| 4 | LR | N0 | N2-nearest | 5,581 | 0.4994 | 0.5000 | 0.6412 |
| 5 | RF | N0 | N0 | 5,583 | 0.9663 | 0.9691 | 0.8923 |
| 6 | RF | N0 | N2-decile | 5,581 | 0.5038 | 0.5025 | 0.6282 |
| 7 | RF | N0 | N2-exact | 5,581 | 0.4980 | 0.4987 | 0.6257 |
| 8 | RF | N0 | N2-nearest | 5,581 | 0.4980 | 0.4987 | 0.6257 |

**Supplementary Table S9 (continued).** Guo degree-only model performance under decile, exact, and nearest-degree matching. Panel B (2 of 3)

| **Row** | **Model** | **Training set** | **Test set** | **MCC** | **Balanced accuracy** | **Accuracy** | **Precision** |
| --- | --- | --- | --- | --- | --- | --- | --- |
| 1 | LR | N0 | N0 | 0.7791 | 0.8894 | 0.8894 | 0.8991 |
| 2 | LR | N0 | N2-decile | 0.0187 | 0.5063 | 0.5063 | 0.5036 |
| 3 | LR | N0 | N2-exact | 0.0265 | 0.5090 | 0.5090 | 0.5052 |
| 4 | LR | N0 | N2-nearest | 0.0265 | 0.5090 | 0.5090 | 0.5052 |
| 5 | RF | N0 | N0 | 0.8037 | 0.8988 | 0.8988 | 0.9541 |
| 6 | RF | N0 | N2-decile | 0.0108 | 0.5040 | 0.5040 | 0.5024 |
| 7 | RF | N0 | N2-exact | -0.0037 | 0.4987 | 0.4987 | 0.4992 |
| 8 | RF | N0 | N2-nearest | -0.0037 | 0.4987 | 0.4987 | 0.4992 |

**Supplementary Table S9 (continued).** Guo degree-only model performance under decile, exact, and nearest-degree matching. Panel C (3 of 3)

| **Row** | **Model** | **Training set** | **Test set** | **Recall** | **Specificity** |
| --- | --- | --- | --- | --- | --- |
| 1 | LR | N0 | N0 | 0.8774 | 0.9015 |
| 2 | LR | N0 | N2-decile | 0.8774 | 0.1352 |
| 3 | LR | N0 | N2-exact | 0.8774 | 0.1406 |
| 4 | LR | N0 | N2-nearest | 0.8774 | 0.1406 |
| 5 | RF | N0 | N0 | 0.8380 | 0.9597 |
| 6 | RF | N0 | N2-decile | 0.8380 | 0.1701 |
| 7 | RF | N0 | N2-exact | 0.8380 | 0.1594 |
| 8 | RF | N0 | N2-nearest | 0.8380 | 0.1594 |

Degree-only models are trained on N0; the evaluation negative set changes. Negative counts refer to the full generated set, before its training/test partition. Main Figure 4 visualizes the endpoint-degree distributions summarized in Tables S7–S9; as with those tables, the distributions are descriptive because endpoint occurrences are dependent and degree is derived from the observed positive graph.

**S6 Topology, substitution, and complete sensitivity results**

Degree-only N0 ROC-AUC was 0.962–0.966, compared with 0.540–0.551 for length-only controls. Adding degree to ESM-2 features yielded logistic-regression ROC-AUC 0.928 on N0 and 0.488 on N2. N6 and N7 each contained 5,581 negatives; the saved N7 assignment comprised 2,191 clusters, with zero within-pair cluster matches and no additional omissions from cluster-exclusion exhaustion. N0-trained N6/N7 sequence-model ROC-AUC was 0.482/0.494. R3/N0 MLP Brier score decreased from 0.344 to 0.253 with Platt scaling, whereas the random-forest Brier score increased from 0.234 to 0.239. The R3/N2 isotonic MLP had Brier score 0.250 and ROC-AUC 0.502. Random-forest N2 ECE ranged from 0.002 to 0.044 with equal-width bins and 0.051 to 0.093 with equal-frequency bins. Corrected N0–N2 contrast P values exceeded 0.05. The separate physicochemical-only five-fold baselines yielded mean ROC-AUC 0.588–0.617.

**Supplementary Table S10.** Degree-only and length-only controls across negative-set training/evaluation directions

| **Feature** | **Model** | **Design** | **Training set** | **Test set** | **ROC-AUC** | **PR-AUC** | **MCC** |
| --- | --- | --- | --- | --- | --- | --- | --- |
| Degree | LR | A | N0 | N0 | 0.9624 | 0.9650 | 0.7791 |
| Degree | LR | A | N0 | N1 | 0.8089 | 0.7677 | 0.4863 |
| Degree | LR | A | N0 | N2 | 0.5025 | 0.5055 | 0.0187 |
| Degree | LR | A | N0 | N3 | 0.6996 | 0.6496 | 0.3326 |
| Degree | LR | A | N0 | N5 | 0.5012 | 0.5016 | 0.0290 |
| Degree | LR | B | N0 | N0 | 0.9624 | 0.9650 | 0.7791 |
| Degree | LR | B | N1 | N0 | 0.9627 | 0.9652 | 0.7474 |
| Degree | LR | B | N2 | N0 | 0.8689 | 0.8839 | 0.5414 |
| Degree | LR | B | N3 | N0 | 0.9503 | 0.9547 | 0.7093 |
| Degree | LR | B | N5 | N0 | 0.9327 | 0.9498 | 0.5819 |
| Degree | RF | A | N0 | N0 | 0.9663 | 0.9691 | 0.8037 |
| Degree | RF | A | N0 | N1 | 0.8161 | 0.7725 | 0.4959 |
| Degree | RF | A | N0 | N2 | 0.5038 | 0.5025 | 0.0108 |
| Degree | RF | A | N0 | N3 | 0.7044 | 0.6487 | 0.3211 |
| Degree | RF | A | N0 | N5 | 0.5038 | 0.5015 | 0.0024 |
| Degree | RF | B | N0 | N0 | 0.9663 | 0.9691 | 0.8037 |
| Degree | RF | B | N1 | N0 | 0.9627 | 0.9650 | 0.7848 |
| Degree | RF | B | N2 | N0 | 0.7974 | 0.6885 | 0.3011 |
| Degree | RF | B | N3 | N0 | 0.8998 | 0.8682 | 0.7037 |
| Degree | RF | B | N5 | N0 | 0.7735 | 0.6650 | 0.2671 |
| Length | LR | A | N0 | N0 | 0.5403 | 0.5320 | 0.0639 |
| Length | LR | A | N0 | N1 | 0.5109 | 0.5137 | 0.0281 |
| Length | LR | A | N0 | N2 | 0.5061 | 0.5091 | 0.0190 |
| Length | LR | A | N0 | N3 | 0.5093 | 0.5134 | 0.0199 |
| Length | LR | A | N0 | N5 | 0.5086 | 0.5115 | 0.0118 |
| Length | LR | B | N0 | N0 | 0.5403 | 0.5320 | 0.0639 |
| Length | LR | B | N1 | N0 | 0.5348 | 0.5291 | 0.0607 |
| Length | LR | B | N2 | N0 | 0.4904 | 0.4946 | -0.0055 |
| Length | LR | B | N3 | N0 | 0.4636 | 0.4703 | -0.0542 |
| Length | LR | B | N5 | N0 | 0.4725 | 0.4740 | -0.0137 |
| Length | RF | A | N0 | N0 | 0.5513 | 0.5454 | 0.0555 |
| Length | RF | A | N0 | N1 | 0.5243 | 0.5232 | 0.0304 |
| Length | RF | A | N0 | N2 | 0.5005 | 0.4988 | -0.0009 |
| Length | RF | A | N0 | N3 | 0.5139 | 0.5084 | 0.0054 |
| Length | RF | A | N0 | N5 | 0.4935 | 0.4923 | -0.0251 |
| Length | RF | B | N0 | N0 | 0.5513 | 0.5454 | 0.0555 |
| Length | RF | B | N1 | N0 | 0.5289 | 0.5228 | 0.0331 |
| Length | RF | B | N2 | N0 | 0.5060 | 0.4928 | 0.0108 |
| Length | RF | B | N3 | N0 | 0.5045 | 0.4959 | 0.0216 |
| Length | RF | B | N5 | N0 | 0.4778 | 0.4812 | -0.0253 |

Each control uses the sum, absolute difference, minimum, and maximum of endpoint degree or length. Designs A and B change the evaluation and training negative definition, respectively.

**Supplementary Table S11.** ESM-2, degree, and combined-feature logistic-regression performance

| **Features** | **Training set** | **Test set** | **ROC-AUC** | **PR-AUC** | **MCC** |
| --- | --- | --- | --- | --- | --- |
| ESM-2 | N0 | N0 | 0.7440 | 0.7325 | 0.3591 |
| ESM-2 | N0 | N2 | 0.4672 | 0.4804 | -0.0582 |
| ESM-2 | N0 | N5 | 0.4771 | 0.4931 | -0.0503 |
| Degree | N0 | N0 | 0.9624 | 0.9650 | 0.7791 |
| Degree | N0 | N2 | 0.5025 | 0.5055 | 0.0187 |
| Degree | N0 | N5 | 0.5012 | 0.5016 | 0.0290 |
| ESM-2 + degree | N0 | N0 | 0.9281 | 0.9378 | 0.7037 |
| ESM-2 + degree | N0 | N2 | 0.4879 | 0.4941 | -0.0152 |
| ESM-2 + degree | N0 | N5 | 0.4889 | 0.4977 | -0.0113 |

All models are trained on N0 and evaluated after test-negative replacement. Sequence features are derived from frozen ESM-2 mean embeddings.

**Supplementary Table S12.** One-sided N2 versus two-sided N6 degree-matched negative controls

| **Representation** | **Training set** | **Test set** | **ROC-AUC** | **PR-AUC** | **MCC** |
| --- | --- | --- | --- | --- | --- |
| ESM-2–MLP | N0 | N0 | 0.7542 | 0.7437 | 0.3808 |
| ESM-2–MLP | N0 | N2 | 0.4675 | 0.4822 | -0.0475 |
| ESM-2–MLP | N0 | N6 | 0.4820 | 0.4958 | -0.0575 |
| Degree-only LR | N0 | N0 | 0.9624 | 0.9650 | 0.7791 |
| Degree-only LR | N0 | N2 | 0.5025 | 0.5055 | 0.0187 |
| Degree-only LR | N0 | N6 | 0.5026 | 0.5060 | 0.0290 |

N2 uses one-sided substitution and N6 permits two-sided substitution. Both contain 5,581 negatives. Models are trained on N0; only test negatives are replaced.

**Supplementary Table S13.** ESM-2–MLP sensitivity to within-pair cluster exclusion in N7

| **Design** | **Training set** | **Test set** | **ROC-AUC** | **PR-AUC** | **MCC** |
| --- | --- | --- | --- | --- | --- |
| A | N0 | N0 | 0.7542 | 0.7437 | 0.3808 |
| A | N0 | N2 | 0.4675 | 0.4822 | -0.0475 |
| A | N0 | N5 | 0.4942 | 0.4985 | -0.0038 |
| A | N0 | N7 | 0.4939 | 0.4982 | -0.0038 |
| B | N0 | N0 | 0.7542 | 0.7437 | 0.3808 |
| B | N2 | N0 | 0.4702 | 0.4733 | -0.0463 |
| B | N5 | N0 | 0.4783 | 0.4755 | -0.0338 |
| B | N7 | N0 | 0.4783 | 0.4755 | -0.0338 |

N7 adds within-pair exclusion using the saved 20%-identity clusters to N5 degree–length matching. This is a negative-pair constraint, distinct from train/test sequence-cluster exclusion.

**S7 Sequence-model negative-definition sensitivity**

Random resampling and length matching reduced sequence-model discrimination relative to the original benchmark negatives. With N0 training, ESM-2–MLP ROC-AUC was 0.647 on N1 and 0.565 on N3, compared with 0.754 on N0, 0.467 on N2, and 0.494 on N5. Training on N1 or N3 and evaluating on the common N0 test set yielded 0.666 and 0.593, respectively. Both directions show dependence on the negative definition; neither isolates a causal effect of degree or length. The consistent-negative revalidation is reported separately in Table S15.

**Supplementary Table S14.** ESM-2–MLP performance across negative-set training/evaluation directions. Panel A (1 of 3)

| **Row** | **Design** | **Training set** | **Test set** | **ROC-AUC** | **PR-AUC** | **MCC** | **F1** |
| --- | --- | --- | --- | --- | --- | --- | --- |
| 1 | A | N0 | N0 | 0.7542 | 0.7437 | 0.3808 | 0.6840 |
| 2 | A | N0 | N1 | 0.6465 | 0.6121 | 0.2122 | 0.6294 |
| 3 | A | N0 | N2 | 0.4675 | 0.4822 | -0.0475 | 0.5623 |
| 4 | A | N0 | N3 | 0.5648 | 0.5515 | 0.0844 | 0.5935 |
| 5 | A | N0 | N5 | 0.4942 | 0.4985 | -0.0038 | 0.5720 |
| 6 | B | N0 | N0 | 0.7542 | 0.7437 | 0.3808 | 0.6840 |
| 7 | B | N1 | N0 | 0.6663 | 0.6346 | 0.2438 | 0.6017 |
| 8 | B | N2 | N0 | 0.4702 | 0.4733 | -0.0463 | 0.4302 |
| 9 | B | N3 | N0 | 0.5926 | 0.5667 | 0.1451 | 0.5343 |
| 10 | B | N5 | N0 | 0.4783 | 0.4755 | -0.0338 | 0.5294 |

**Supplementary Table S14 (continued).** ESM-2–MLP performance across negative-set training/evaluation directions. Panel B (2 of 3)

| **Row** | **Design** | **Training set** | **Test set** | **Balanced accuracy** | **Accuracy** | **Precision** | **Recall** |
| --- | --- | --- | --- | --- | --- | --- | --- |
| 1 | A | N0 | N0 | 0.6902 | 0.6902 | 0.6980 | 0.6705 |
| 2 | A | N0 | N1 | 0.6052 | 0.6052 | 0.5930 | 0.6705 |
| 3 | A | N0 | N2 | 0.4781 | 0.4781 | 0.4842 | 0.6705 |
| 4 | A | N0 | N3 | 0.5407 | 0.5407 | 0.5323 | 0.6705 |
| 5 | A | N0 | N5 | 0.4982 | 0.4982 | 0.4987 | 0.6705 |
| 6 | B | N0 | N0 | 0.6902 | 0.6902 | 0.6980 | 0.6705 |
| 7 | B | N1 | N0 | 0.6213 | 0.6213 | 0.6346 | 0.5721 |
| 8 | B | N2 | N0 | 0.4772 | 0.4772 | 0.4727 | 0.3948 |
| 9 | B | N3 | N0 | 0.5716 | 0.5716 | 0.5853 | 0.4915 |
| 10 | B | N5 | N0 | 0.4834 | 0.4834 | 0.4861 | 0.5810 |

**Supplementary Table S14 (continued).** ESM-2–MLP performance across negative-set training/evaluation directions. Panel C (3 of 3)

| **Row** | **Design** | **Training set** | **Test set** | **Specificity** | **Brier score** | **ECE** |
| --- | --- | --- | --- | --- | --- | --- |
| 1 | A | N0 | N0 | 0.7099 | 0.2528 | 0.2140 |
| 2 | A | N0 | N1 | 0.5398 | 0.3309 | 0.3019 |
| 3 | A | N0 | N2 | 0.2856 | 0.4509 | 0.4248 |
| 4 | A | N0 | N3 | 0.4109 | 0.3884 | 0.3608 |
| 5 | A | N0 | N5 | 0.3259 | 0.4326 | 0.4047 |
| 6 | B | N0 | N0 | 0.7099 | 0.2528 | 0.2140 |
| 7 | B | N1 | N0 | 0.6705 | 0.3010 | 0.2534 |
| 8 | B | N2 | N0 | 0.5595 | 0.2756 | 0.1313 |
| 9 | B | N3 | N0 | 0.6517 | 0.2489 | 0.0733 |
| 10 | B | N5 | N0 | 0.3859 | 0.2648 | 0.0981 |

Frozen ESM-2 mean embeddings and combined symmetric fusion are used throughout. Positive partitions are shared (80/20; seed 42). The N0/N0 baseline is repeated for designs A and B. Probability metrics use raw scores; these are distribution-shift comparisons, not consistent-negative N1/N3 evaluations.

**S8 Protein Representation Comparison Under Consistent Negative-Set Evaluation**

AAC+CTD, ESM-2, and the limited ProtT5 representation were evaluated under matched negative-set definitions across R0 and R3. Under N0, ESM-2 showed higher discrimination than AAC+CTD in both regimes, whereas all three representations showed substantially reduced discrimination under the harder N2 and N5 negative sets. In R3, ROC-AUC for ESM-2 was 0.617 under N0 but approximately 0.52 under N2 and N5, while the limited ProtT5 checkpoint ranged from 0.454 to 0.501. These results support the representation comparison while indicating that the limited ProtT5 experiment should be interpreted only as a sensitivity analysis.

**Supplementary Table S15.** AAC+CTD, ESM-2, and limited ProtT5 comparison with consistent negative-set training and evaluation. Panel A (1 of 4)

| **Row** | **Regime** | **Negative set** | **Representation** | **Train pairs** | **Test pairs** | **ROC-AUC** | **PR-AUC** |
| --- | --- | --- | --- | --- | --- | --- | --- |
| 1 | R0 | N0 | AAC+CTD | 8,930 | 2,234 | 0.7197 | 0.6906 |
| 2 | R0 | N0 | ESM-2 mean | 8,930 | 2,234 | 0.7542 | 0.7437 |
| 3 | R0 | N0 | ProtT5 mean | 8,930 | 2,234 | 0.5279 | 0.5119 |
| 4 | R0 | N2 | AAC+CTD | 8,928 | 2,234 | 0.4551 | 0.4642 |
| 5 | R0 | N2 | ESM-2 mean | 8,928 | 2,234 | 0.4627 | 0.4664 |
| 6 | R0 | N2 | ProtT5 mean | 8,928 | 2,234 | 0.5090 | 0.5114 |
| 7 | R0 | N5 | AAC+CTD | 8,928 | 2,234 | 0.4462 | 0.4643 |
| 8 | R0 | N5 | ESM-2 mean | 8,928 | 2,234 | 0.4628 | 0.4734 |
| 9 | R0 | N5 | ProtT5 mean | 8,928 | 2,234 | 0.4693 | 0.4765 |
| 10 | R3 | N0 | AAC+CTD | 5,703 | 462 | 0.5874 | 0.5491 |
| 11 | R3 | N0 | ESM-2 mean | 5,703 | 462 | 0.6168 | 0.5726 |
| 12 | R3 | N0 | ProtT5 mean | 5,703 | 462 | 0.4535 | 0.4661 |
| 13 | R3 | N2 | AAC+CTD | 5,738 | 446 | 0.5173 | 0.5285 |
| 14 | R3 | N2 | ESM-2 mean | 5,738 | 446 | 0.5166 | 0.5049 |
| 15 | R3 | N2 | ProtT5 mean | 5,738 | 446 | 0.4943 | 0.4949 |
| 16 | R3 | N5 | AAC+CTD | 5,738 | 445 | 0.4618 | 0.4773 |
| 17 | R3 | N5 | ESM-2 mean | 5,738 | 445 | 0.5074 | 0.5065 |
| 18 | R3 | N5 | ProtT5 mean | 5,738 | 445 | 0.5008 | 0.4995 |

**Supplementary Table S15 (continued).** AAC+CTD, ESM-2, and limited ProtT5 comparison with consistent negative-set training and evaluation. Panel B (2 of 4)

| **Row** | **Regime** | **Negative set** | **Representation** | **F1** | **MCC** | **Balanced accuracy** | **Accuracy** |
| --- | --- | --- | --- | --- | --- | --- | --- |
| 1 | R0 | N0 | AAC+CTD | 0.6786 | 0.3291 | 0.6638 | 0.6638 |
| 2 | R0 | N0 | ESM-2 mean | 0.6840 | 0.3808 | 0.6902 | 0.6902 |
| 3 | R0 | N0 | ProtT5 mean | 0.6067 | 0.0652 | 0.5300 | 0.5300 |
| 4 | R0 | N2 | AAC+CTD | 0.4156 | -0.0610 | 0.4700 | 0.4700 |
| 5 | R0 | N2 | ESM-2 mean | 0.4242 | -0.0723 | 0.4642 | 0.4642 |
| 6 | R0 | N2 | ProtT5 mean | 0.5454 | 0.0181 | 0.5090 | 0.5090 |
| 7 | R0 | N5 | AAC+CTD | 0.4309 | -0.0739 | 0.4633 | 0.4633 |
| 8 | R0 | N5 | ESM-2 mean | 0.5211 | -0.0699 | 0.4660 | 0.4660 |
| 9 | R0 | N5 | ProtT5 mean | 0.4199 | -0.0448 | 0.4781 | 0.4781 |
| 10 | R3 | N0 | AAC+CTD | 0.5738 | 0.1291 | 0.5643 | 0.5628 |
| 11 | R3 | N0 | ESM-2 mean | 0.5442 | 0.1488 | 0.5740 | 0.5758 |
| 12 | R3 | N0 | ProtT5 mean | 0.4306 | -0.0362 | 0.4821 | 0.4848 |
| 13 | R3 | N2 | AAC+CTD | 0.4365 | -0.0543 | 0.4731 | 0.4731 |
| 14 | R3 | N2 | ESM-2 mean | 0.4433 | 0.0092 | 0.5045 | 0.5045 |
| 15 | R3 | N2 | ProtT5 mean | 0.5447 | 0.0406 | 0.5202 | 0.5202 |
| 16 | R3 | N5 | AAC+CTD | 0.6004 | -0.0146 | 0.4938 | 0.4944 |
| 17 | R3 | N5 | ESM-2 mean | 0.5056 | 0.0067 | 0.5034 | 0.5034 |
| 18 | R3 | N5 | ProtT5 mean | 0.2956 | -0.0067 | 0.4973 | 0.4966 |

**Supplementary Table S15 (continued).** AAC+CTD, ESM-2, and limited ProtT5 comparison with consistent negative-set training and evaluation. Panel C (3 of 4)

| **Row** | **Regime** | **Negative set** | **Representation** | **Precision** | **Recall** | **Specificity** | **Brier score** |
| --- | --- | --- | --- | --- | --- | --- | --- |
| 1 | R0 | N0 | AAC+CTD | 0.6500 | 0.7099 | 0.6177 | 0.2533 |
| 2 | R0 | N0 | ESM-2 mean | 0.6980 | 0.6705 | 0.7099 | 0.2528 |
| 3 | R0 | N0 | ProtT5 mean | 0.5216 | 0.7252 | 0.3348 | 0.2544 |
| 4 | R0 | N2 | AAC+CTD | 0.4631 | 0.3769 | 0.5631 | 0.2820 |
| 5 | R0 | N2 | ESM-2 mean | 0.4584 | 0.3948 | 0.5336 | 0.2779 |
| 6 | R0 | N2 | ProtT5 mean | 0.5077 | 0.5891 | 0.4288 | 0.2629 |
| 7 | R0 | N5 | AAC+CTD | 0.4586 | 0.4064 | 0.5201 | 0.2823 |
| 8 | R0 | N5 | ESM-2 mean | 0.4723 | 0.5810 | 0.3509 | 0.2664 |
| 9 | R0 | N5 | ProtT5 mean | 0.4726 | 0.3778 | 0.5783 | 0.2765 |
| 10 | R3 | N0 | AAC+CTD | 0.5418 | 0.6099 | 0.5188 | 0.2727 |
| 11 | R3 | N0 | ESM-2 mean | 0.5652 | 0.5247 | 0.6234 | 0.3123 |
| 12 | R3 | N0 | ProtT5 mean | 0.4615 | 0.4036 | 0.5607 | 0.2900 |
| 13 | R3 | N2 | AAC+CTD | 0.4691 | 0.4081 | 0.5381 | 0.2553 |
| 14 | R3 | N2 | ESM-2 mean | 0.5057 | 0.3946 | 0.6143 | 0.2639 |
| 15 | R3 | N2 | ProtT5 mean | 0.5182 | 0.5740 | 0.4664 | 0.2649 |
| 16 | R3 | N5 | AAC+CTD | 0.4971 | 0.7578 | 0.2297 | 0.2686 |
| 17 | R3 | N5 | ESM-2 mean | 0.5045 | 0.5067 | 0.5000 | 0.4111 |
| 18 | R3 | N5 | ProtT5 mean | 0.4947 | 0.2108 | 0.7838 | 0.2772 |

**Supplementary Table S15 (continued).** AAC+CTD, ESM-2, and limited ProtT5 comparison with consistent negative-set training and evaluation. Panel D (4 of 4)

| **Row** | **Regime** | **Negative set** | **Representation** | **ECE** |
| --- | --- | --- | --- | --- |
| 1 | R0 | N0 | AAC+CTD | 0.1863 |
| 2 | R0 | N0 | ESM-2 mean | 0.2140 |
| 3 | R0 | N0 | ProtT5 mean | 0.0487 |
| 4 | R0 | N2 | AAC+CTD | 0.1450 |
| 5 | R0 | N2 | ESM-2 mean | 0.1449 |
| 6 | R0 | N2 | ProtT5 mean | 0.0899 |
| 7 | R0 | N5 | AAC+CTD | 0.1480 |
| 8 | R0 | N5 | ESM-2 mean | 0.1137 |
| 9 | R0 | N5 | ProtT5 mean | 0.1318 |
| 10 | R3 | N0 | AAC+CTD | 0.1467 |
| 11 | R3 | N0 | ESM-2 mean | 0.2531 |
| 12 | R3 | N0 | ProtT5 mean | 0.1762 |
| 13 | R3 | N2 | AAC+CTD | 0.1113 |
| 14 | R3 | N2 | ESM-2 mean | 0.1052 |
| 15 | R3 | N2 | ProtT5 mean | 0.0908 |
| 16 | R3 | N5 | AAC+CTD | 0.1030 |
| 17 | R3 | N5 | ESM-2 mean | 0.3801 |
| 18 | R3 | N5 | ProtT5 mean | 0.1404 |

All representations use the MLP with the same negative definition in training and evaluation. The limited ProtT5 checkpoint is a sensitivity check, not a general comparison of protein-language-model families. Supplementary Figure S3 visualizes the consistent-negative revalidation summarized in Table S15 in which the same negative definition is used for model fitting and evaluation rather than only swapping the test negatives.

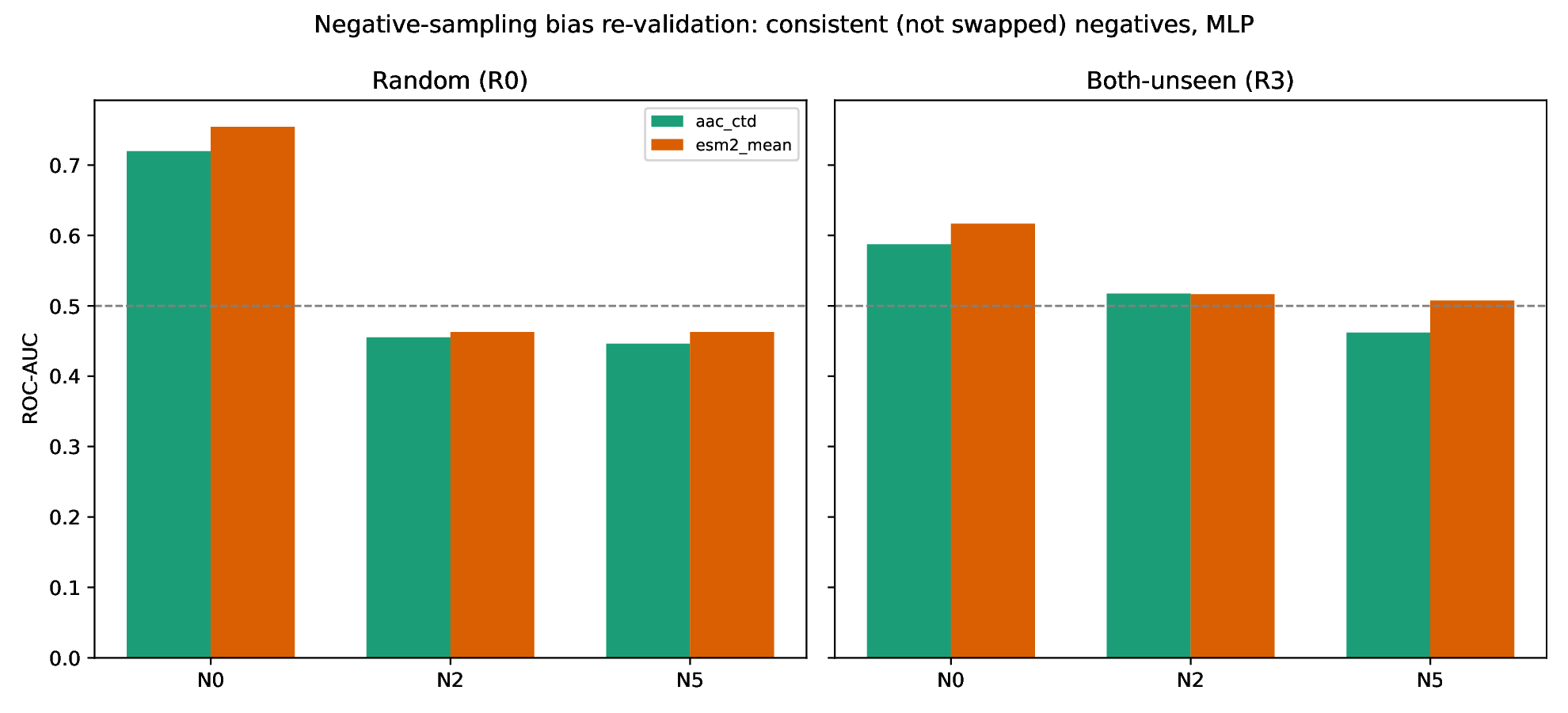

**Supplementary Figure S3**. Consistent negative-set revalidation. AAC+CTD and ESM-2 models are trained and evaluated under the same negative-set definition for R0 and R3. The figure visualizes the point estimates reported in Supplementary Table S15.

**S9 Single-seed model-configuration comparison**

R3/N2 ROC-AUC ranged from 0.493 to 0.547 and PR-AUC from 0.498 to 0.544. These point estimates do not establish equivalence to chance.

**Supplementary Table S16.** Single-seed model-configuration performance under R3/N0 and R3/N2. Panel A (1 of 4)

| **Row** | **Negative set** | **Model** | **Train pairs** | **Test pairs** | **ROC-AUC** | **PR-AUC** |
| --- | --- | --- | --- | --- | --- | --- |
| 1 | N0 | LR | 5,703 | 462 | 0.6041 | 0.5500 |
| 2 | N0 | RF | 5,703 | 462 | 0.6322 | 0.5883 |
| 3 | N0 | MLP | 5,703 | 462 | 0.6168 | 0.5726 |
| 4 | N0 | Siamese | 5,703 | 462 | 0.6512 | 0.6306 |
| 5 | N0 | Attention | 5,703 | 462 | 0.6245 | 0.6172 |
| 6 | N0 | Cross-interaction | 5,703 | 462 | 0.6132 | 0.5961 |
| 7 | N2 | LR | 5,738 | 446 | 0.4998 | 0.5047 |
| 8 | N2 | RF | 5,738 | 446 | 0.5184 | 0.4976 |
| 9 | N2 | MLP | 5,738 | 446 | 0.5166 | 0.5049 |
| 10 | N2 | Siamese | 5,738 | 446 | 0.5119 | 0.5094 |
| 11 | N2 | Attention | 5,738 | 446 | 0.5473 | 0.5441 |
| 12 | N2 | Cross-interaction | 5,738 | 446 | 0.4933 | 0.5001 |

**Supplementary Table S16 (continued).** Single-seed model-configuration performance under R3/N0 and R3/N2. Panel B (2 of 4)

| **Row** | **Negative set** | **Model** | **F1** | **MCC** | **Balanced accuracy** | **Accuracy** |
| --- | --- | --- | --- | --- | --- | --- |
| 1 | N0 | LR | 0.5463 | 0.1701 | 0.5842 | 0.5866 |
| 2 | N0 | RF | 0.5983 | 0.2042 | 0.6022 | 0.6017 |
| 3 | N0 | MLP | 0.5442 | 0.1488 | 0.5740 | 0.5758 |
| 4 | N0 | Siamese | 0.5728 | 0.2226 | 0.6101 | 0.6126 |
| 5 | N0 | Attention | 0.5600 | 0.1424 | 0.5712 | 0.5714 |
| 6 | N0 | Cross-interaction | 0.5738 | 0.1059 | 0.5524 | 0.5498 |
| 7 | N2 | LR | 0.4933 | -0.0135 | 0.4933 | 0.4933 |
| 8 | N2 | RF | 0.5470 | 0.0718 | 0.5359 | 0.5359 |
| 9 | N2 | MLP | 0.4433 | 0.0092 | 0.5045 | 0.5045 |
| 10 | N2 | Siamese | 0.1374 | -0.0238 | 0.4933 | 0.4933 |
| 11 | N2 | Attention | 0.5121 | 0.0952 | 0.5471 | 0.5471 |
| 12 | N2 | Cross-interaction | 0.1450 | -0.0079 | 0.4978 | 0.4978 |

**Supplementary Table S16 (continued).** Single-seed model-configuration performance under R3/N0 and R3/N2. Panel C (3 of 4)

| **Row** | **Negative set** | **Model** | **Precision** | **Recall** | **Specificity** | **Brier score** |
| --- | --- | --- | --- | --- | --- | --- |
| 1 | N0 | LR | 0.5808 | 0.5157 | 0.6527 | 0.3326 |
| 2 | N0 | RF | 0.5830 | 0.6143 | 0.5900 | 0.2365 |
| 3 | N0 | MLP | 0.5652 | 0.5247 | 0.6234 | 0.3123 |
| 4 | N0 | Siamese | 0.6122 | 0.5381 | 0.6820 | 0.2828 |
| 5 | N0 | Attention | 0.5551 | 0.5650 | 0.5774 | 0.2597 |
| 6 | N0 | Cross-interaction | 0.5283 | 0.6278 | 0.4770 | 0.2565 |
| 7 | N2 | LR | 0.4933 | 0.4933 | 0.4933 | 0.3320 |
| 8 | N2 | RF | 0.5342 | 0.5605 | 0.5112 | 0.2507 |
| 9 | N2 | MLP | 0.5057 | 0.3946 | 0.6143 | 0.2639 |
| 10 | N2 | Siamese | 0.4615 | 0.0807 | 0.9058 | 0.2506 |
| 11 | N2 | Attention | 0.5550 | 0.4753 | 0.6188 | 0.2499 |
| 12 | N2 | Cross-interaction | 0.4872 | 0.0852 | 0.9103 | 0.2524 |

**Supplementary Table S16 (continued).** Single-seed model-configuration performance under R3/N0 and R3/N2. Panel D (4 of 4)

| **Row** | **Negative set** | **Model** | **ECE** |
| --- | --- | --- | --- |
| 1 | N0 | LR | 0.2910 |
| 2 | N0 | RF | 0.0312 |
| 3 | N0 | MLP | 0.2531 |
| 4 | N0 | Siamese | 0.2154 |
| 5 | N0 | Attention | 0.1472 |
| 6 | N0 | Cross-interaction | 0.1405 |
| 7 | N2 | LR | 0.2555 |
| 8 | N2 | RF | 0.0224 |
| 9 | N2 | MLP | 0.1052 |
| 10 | N2 | Siamese | 0.0299 |
| 11 | N2 | Attention | 0.0325 |
| 12 | N2 | Cross-interaction | 0.0391 |

All model configurations use frozen ESM-2 representations. Each negative definition is used consistently during training and evaluation; metrics are single-seed estimates.

**S10 Model Configuration and Training-Seed Variability**

Table S17 evaluates six model configurations across five training seeds under R3. For N0, the models showed relatively similar mean ROC-AUC values, ranging from approximately 0.60 to 0.64, with modest between-seed variability. Under N2, discrimination was substantially reduced, with mean ROC-AUC values concentrated close to 0.50–0.53 across configurations. The corresponding PR-AUC values followed the same pattern, indicating that the choice of model configuration had limited ability to overcome the more stringent negative-set construction. The reported standard deviations quantify training-seed variability rather than independent biological replication.

**Supplementary Table S17.** R3 model-configuration performance across five model-training seeds. Panel A (1 of 2).

| **Row** | **Negative set** | **Model** | **ROC-AUC mean** | **ROC-AUC SD** | **PR-AUC mean** | **PR-AUC SD** |
| --- | --- | --- | --- | --- | --- | --- |
| 1 | N0 | Attention | 0.6384 | 0.0103 | 0.6245 | 0.0067 |
| 2 | N0 | Cross-interaction | 0.6375 | 0.0232 | 0.6143 | 0.0236 |
| 3 | N0 | LR | 0.6041 | 0.0000 | 0.5500 | 0.0000 |
| 4 | N0 | MLP | 0.6134 | 0.0118 | 0.5824 | 0.0197 |
| 5 | N0 | RF | 0.6268 | 0.0049 | 0.5901 | 0.0054 |
| 6 | N0 | Siamese | 0.6296 | 0.0180 | 0.6129 | 0.0172 |
| 7 | N2 | Attention | 0.5249 | 0.0182 | 0.5171 | 0.0213 |
| 8 | N2 | Cross-interaction | 0.5075 | 0.0096 | 0.5050 | 0.0049 |
| 9 | N2 | LR | 0.4998 | 0.0000 | 0.5047 | 0.0000 |
| 10 | N2 | MLP | 0.5149 | 0.0159 | 0.5157 | 0.0137 |
| 11 | N2 | RF | 0.5136 | 0.0269 | 0.5121 | 0.0130 |
| 12 | N2 | Siamese | 0.5127 | 0.0090 | 0.5204 | 0.0127 |

**Supplementary Table S17 (continued).** R3 model-configuration performance across five model-training seeds. Panel B (2 of 2)

| **Row** | **Negative set** | **Model** | **MCC mean** | **MCC SD** | **Seeds** |
| --- | --- | --- | --- | --- | --- |
| 1 | N0 | Attention | 0.1780 | 0.0408 | 5 |
| 2 | N0 | Cross-interaction | 0.1656 | 0.0386 | 5 |
| 3 | N0 | LR | 0.1701 | 0.0000 | 5 |
| 4 | N0 | MLP | 0.1510 | 0.0138 | 5 |
| 5 | N0 | RF | 0.1833 | 0.0171 | 5 |
| 6 | N0 | Siamese | 0.2036 | 0.0363 | 5 |
| 7 | N2 | Attention | 0.0367 | 0.0431 | 5 |
| 8 | N2 | Cross-interaction | 0.0055 | 0.0199 | 5 |
| 9 | N2 | LR | -0.0135 | 0.0000 | 5 |
| 10 | N2 | MLP | 0.0159 | 0.0349 | 5 |
| 11 | N2 | RF | 0.0386 | 0.0480 | 5 |
| 12 | N2 | Siamese | 0.0041 | 0.0281 | 5 |

The split and negative realization are fixed. SD describes variability across five training seeds, not independent biological replication.

**S11 Component-aware inference and component dominance**

Strict N0 and N2 test graphs supplied 19 and five usable components. The N0 Siamese ROC-AUC test had adjusted P = 0.022 under bootstrap inference and 0.999 under jackknife inference. No N2 model configuration met the corrected above-chance criterion. Neither paired architecture nor N0–N2 contrasts reached corrected significance (Tables S40–S41). Giant-component exclusion left only 25 N0 and eight N2 pairs. Supplementary Figure S4 visualizes the corresponding jackknife intervals.

**Supplementary Table S18.** Hypotheses and multiple-testing correction for the 44-test inferential family

| **Question** | **Null hypothesis** | **Metric** | **Comparison** | **N tests** | **Test** |
| --- | --- | --- | --- | --- | --- |
| Q1a: Is ROC-AUC distinguishable from chance? | ROC-AUC = 0.5 | ROC-AUC | 6 models x 2 negative sets (N0, N2) | 12 | Percentile connected-component bootstrap / delete-one-component jackknife |
| Q1b: Is MCC distinguishable from chance? | MCC = 0 | MCC | 6 models x 2 negative sets (N0, N2) | 12 | Percentile connected-component bootstrap / delete-one-component jackknife |
| Q1c: Is PR-AUC distinguishable from chance? | PR-AUC = test-set prevalence | PR-AUC | 6 models x 2 negative sets (N0, N2) | 12 | Percentile connected-component bootstrap / delete-one-component jackknife |
| Q2: Does the Siamese model differ from the logistic-regression baseline? | ΔROC-AUC (Siamese MLP - LR) = 0 | ΔROC-AUC | Siamese MLP vs. LR, same test set, 2 negative sets | 2 | Paired percentile bootstrap / paired jackknife |
| Q3: Does the N0-vs-N2 gap differ from zero? | ROC-AUC(N0) - ROC-AUC(N2) = 0 | ΔROC-AUC | 6 models, different (unpaired) N0 vs. N2 test-pair sets | 6 | Unpaired percentile bootstrap / unpaired jackknife (Welch-Satterthwaite) |

BH correction is applied separately to each method’s complete 44-test family. The PR-AUC null is the realized positive prevalence; the degree-only controls are outside this family.

**Supplementary Table S19.** R3 above-chance tests using component bootstrap and jackknife inference. Panel A (1 of 7). ROC-AUC — bootstrap

| **Negative set** | **Model** | **ROC-AUC** | **ROC-AUC Boot. 95% CI lower** | **ROC-AUC Boot. 95% CI upper** | **ROC-AUC Boot. P** | **ROC-AUC Boot. BH-adjusted P** | **ROC-AUC Boot. above chance (BH)** |
| --- | --- | --- | --- | --- | --- | --- | --- |
| N0 | LR | 0.6041 | 0.4791 | 0.7804 | 0.0610 | 0.1299 | No |
| N0 | RF | 0.6322 | 0.5166 | 0.8500 | 0.0390 | 0.1144 | No |
| N0 | MLP | 0.6168 | 0.5524 | 0.9112 | 0.0240 | 0.0880 | No |
| N0 | Siamese | 0.6512 | 0.6303 | 0.9725 | 0.0030 | 0.0220 | Yes |
| N0 | Attention | 0.6245 | 0.5000 | 0.9091 | 0.0510 | 0.1296 | No |
| N0 | Cross-interaction | 0.6132 | 0.4916 | 0.8741 | 0.0620 | 0.1299 | No |
| N2 | LR | 0.4998 | 0.0800 | 0.6000 | 0.9660 | 0.9660 | No |
| N2 | RF | 0.5184 | 0.2800 | 0.7600 | 0.2680 | 0.3685 | No |
| N2 | MLP | 0.5166 | 0.3600 | 0.9600 | 0.1480 | 0.2606 | No |
| N2 | Siamese | 0.5119 | 0.1600 | 0.7200 | 0.4810 | 0.6047 | No |
| N2 | Attention | 0.5473 | 0.0400 | 0.5504 | 0.6630 | 0.6946 | No |
| N2 | Cross-interaction | 0.4933 | 0.1600 | 0.6400 | 0.2200 | 0.3227 | No |

**Supplementary Table S19 (continued).** R3 above-chance tests using component bootstrap and jackknife inference. Panel B (2 of 7). ROC-AUC — jackknife

| **Negative set** | **Model** | **ROC-AUC** | **ROC-AUC Jack. 95% CI lower** | **ROC-AUC Jack. 95% CI upper** | **ROC-AUC Jack. P** | **ROC-AUC Jack. BH-adjusted P** | **ROC-AUC Jack. above chance (BH)** |
| --- | --- | --- | --- | --- | --- | --- | --- |
| N0 | LR | 0.6041 | 0.5297 | 0.6786 | 0.0088 | 0.1750 | No |
| N0 | RF | 0.6322 | 0.5177 | 0.7467 | 0.0260 | 0.2862 | No |
| N0 | MLP | 0.6168 | 0.3756 | 0.8579 | 0.3225 | 0.9993 | No |
| N0 | Siamese | 0.6512 | 0.3256 | 0.9767 | 0.3422 | 0.9993 | No |
| N0 | Attention | 0.6245 | 0.4373 | 0.8116 | 0.1793 | 0.8765 | No |
| N0 | Cross-interaction | 0.6132 | 0.4929 | 0.7336 | 0.0636 | 0.5598 | No |
| N2 | LR | 0.4998 | 0.0814 | 0.9182 | 0.9991 | 0.9993 | No |
| N2 | RF | 0.5184 | 0.4067 | 0.6301 | 0.6707 | 0.9993 | No |
| N2 | MLP | 0.5166 | 0.1341 | 0.8990 | 0.9101 | 0.9993 | No |
| N2 | Siamese | 0.5119 | 0.3470 | 0.6768 | 0.8515 | 0.9993 | No |
| N2 | Attention | 0.5473 | -0.1149 | 1.2096 | 0.8523 | 0.9993 | No |
| N2 | Cross-interaction | 0.4933 | 0.3678 | 0.6188 | 0.8892 | 0.9993 | No |

**Supplementary Table S19 (continued).** R3 above-chance tests using component bootstrap and jackknife inference. Panel C (3 of 7). PR-AUC — bootstrap

| **Negative set** | **Model** | **PR-AUC** | **PR-AUC Boot. 95% CI lower** | **PR-AUC Boot. 95% CI upper** | **PR-AUC Boot. P** | **PR-AUC Boot. BH-adjusted P** | **PR-AUC Boot. above chance (BH)** |
| --- | --- | --- | --- | --- | --- | --- | --- |
| N0 | LR | 0.5500 | 0.5222 | 0.8590 | 0.0280 | 0.0880 | No |
| N0 | RF | 0.5883 | 0.5767 | 0.8952 | 0.0040 | 0.0251 | Yes |
| N0 | MLP | 0.5726 | 0.5618 | 0.9456 | 0.0010 | 0.0110 | Yes |
| N0 | Siamese | 0.6306 | 0.6206 | 0.9720 | 0.0000 | 0.0000 | Yes |
| N0 | Attention | 0.6172 | 0.6053 | 0.9211 | 0.0030 | 0.0220 | Yes |
| N0 | Cross-interaction | 0.5961 | 0.5829 | 0.9143 | 0.0080 | 0.0440 | Yes |
| N2 | LR | 0.5047 | 0.4141 | 0.7889 | 0.1540 | 0.2606 | No |
| N2 | RF | 0.4976 | 0.4500 | 0.8254 | 0.7660 | 0.7838 | No |
| N2 | MLP | 0.5049 | 0.5017 | 0.9667 | 0.0180 | 0.0880 | No |
| N2 | Siamese | 0.5094 | 0.4500 | 0.7229 | 0.2300 | 0.3265 | No |
| N2 | Attention | 0.5441 | 0.4111 | 0.5456 | 0.6630 | 0.6946 | No |
| N2 | Cross-interaction | 0.5001 | 0.4500 | 0.8000 | 0.6180 | 0.6946 | No |

**Supplementary Table S19 (continued).** R3 above-chance tests using component bootstrap and jackknife inference. Panel D (4 of 7). PR-AUC — jackknife

| **Negative set** | **Model** | **PR-AUC** | **PR-AUC Jack. 95% CI lower** | **PR-AUC Jack. 95% CI upper** | **PR-AUC Jack. P** | **PR-AUC Jack. BH-adjusted P** | **PR-AUC Jack. above chance (BH)** |
| --- | --- | --- | --- | --- | --- | --- | --- |
| N0 | LR | 0.5500 | 0.2673 | 0.8326 | 0.6230 | 0.9993 | No |
| N0 | RF | 0.5883 | 0.2575 | 0.9192 | 0.5109 | 0.9993 | No |
| N0 | MLP | 0.5726 | 0.0774 | 1.0677 | 0.7074 | 0.9993 | No |
| N0 | Siamese | 0.6306 | 0.1817 | 1.0795 | 0.4976 | 0.9993 | No |
| N0 | Attention | 0.6172 | 0.2729 | 0.9615 | 0.4226 | 0.9993 | No |
| N0 | Cross-interaction | 0.5961 | 0.2513 | 0.9409 | 0.4983 | 0.9993 | No |
| N2 | LR | 0.5047 | 0.3331 | 0.6763 | 0.9432 | 0.9993 | No |
| N2 | RF | 0.4976 | 0.4527 | 0.5425 | 0.8902 | 0.9993 | No |
| N2 | MLP | 0.5049 | -0.1798 | 1.1896 | 0.9851 | 0.9993 | No |
| N2 | Siamese | 0.5094 | 0.4906 | 0.5282 | 0.2365 | 0.9580 | No |
| N2 | Attention | 0.5441 | 0.3500 | 0.7383 | 0.5622 | 0.9993 | No |
| N2 | Cross-interaction | 0.5001 | 0.2231 | 0.7771 | 0.9993 | 0.9993 | No |

**Supplementary Table S19 (continued).** R3 above-chance tests using component bootstrap and jackknife inference. Panel E (5 of 7). MCC — bootstrap

| **Negative set** | **Model** | **MCC** | **MCC Boot. 95% CI lower** | **MCC Boot. 95% CI upper** | **MCC Boot. P** | **MCC Boot. BH-adjusted P** | **MCC Boot. above chance (BH)** |
| --- | --- | --- | --- | --- | --- | --- | --- |
| N0 | LR | 0.1701 | 0.0422 | 0.4907 | 0.0280 | 0.0880 | No |
| N0 | RF | 0.2042 | -0.0156 | 0.5658 | 0.0610 | 0.1299 | No |
| N0 | MLP | 0.1488 | 0.1274 | 0.8461 | 0.0000 | 0.0000 | Yes |
| N0 | Siamese | 0.2226 | 0.2053 | 0.7628 | 0.0000 | 0.0000 | Yes |
| N0 | Attention | 0.1424 | 0.1111 | 0.7366 | 0.0210 | 0.0880 | No |
| N0 | Cross-interaction | 0.1059 | -0.2141 | 0.4693 | 0.2100 | 0.3227 | No |
| N2 | LR | -0.0135 | -0.6000 | 0.0000 | 0.1750 | 0.2852 | No |
| N2 | RF | 0.0718 | -0.6000 | 0.0775 | 0.6630 | 0.6946 | No |
| N2 | MLP | 0.0092 | -0.2182 | 0.8165 | 0.3800 | 0.4918 | No |
| N2 | Siamese | -0.0238 | -0.0246 | 0.0000 | 0.6630 | 0.6946 | No |
| N2 | Attention | 0.0952 | 0.0000 | 0.0965 | 0.6630 | 0.6946 | No |
| N2 | Cross-interaction | -0.0079 | -0.0080 | 0.0000 | 0.6630 | 0.6946 | No |

**Supplementary Table S19 (continued).** R3 above-chance tests using component bootstrap and jackknife inference. Panel F (6 of 7). MCC — jackknife

| **Negative set** | **Model** | **MCC** | **MCC Jack. 95% CI lower** | **MCC Jack. 95% CI upper** | **MCC Jack. P** | **MCC Jack. BH-adjusted P** | **MCC Jack. above chance (BH)** |
| --- | --- | --- | --- | --- | --- | --- | --- |
| N0 | LR | 0.1701 | -0.0504 | 0.3905 | 0.1225 | 0.6736 | No |
| N0 | RF | 0.2042 | 0.0508 | 0.3576 | 0.0119 | 0.1750 | No |
| N0 | MLP | 0.1488 | -0.7796 | 1.0772 | 0.7402 | 0.9993 | No |
| N0 | Siamese | 0.2226 | -0.4160 | 0.8611 | 0.4734 | 0.9993 | No |
| N0 | Attention | 0.1424 | -0.4498 | 0.7345 | 0.6196 | 0.9993 | No |
| N0 | Cross-interaction | 0.1059 | 0.0720 | 0.1397 | 3.55e-06 | 0.0002 | Yes |
| N2 | LR | -0.0135 | -0.5595 | 0.5326 | 0.9487 | 0.9993 | No |
| N2 | RF | 0.0718 | -0.6645 | 0.8082 | 0.7999 | 0.9993 | No |
| N2 | MLP | 0.0092 | -0.5467 | 0.5651 | 0.9656 | 0.9993 | No |
| N2 | Siamese | -0.0238 | -0.0772 | 0.0295 | 0.2830 | 0.9580 | No |
| N2 | Attention | 0.0952 | -0.1171 | 0.3074 | 0.2812 | 0.9580 | No |
| N2 | Cross-interaction | -0.0079 | -0.0256 | 0.0097 | 0.2805 | 0.9580 | No |

**Supplementary Table S19 (continued).** R3 above-chance tests using component bootstrap and jackknife inference. Panel G (7 of 7). Test-set composition and resampling units

| **Negative set** | **Model** | **Positive prevalence** | **Test components** | **Usable components** |
| --- | --- | --- | --- | --- |
| N0 | LR | 0.4827 | 19 | 19 |
| N0 | RF | 0.4827 | 19 | 19 |
| N0 | MLP | 0.4827 | 19 | 19 |
| N0 | Siamese | 0.4827 | 19 | 19 |
| N0 | Attention | 0.4827 | 19 | 19 |
| N0 | Cross-interaction | 0.4827 | 19 | 19 |
| N2 | LR | 0.5000 | 5 | 5 |
| N2 | RF | 0.5000 | 5 | 5 |
| N2 | MLP | 0.5000 | 5 | 5 |
| N2 | Siamese | 0.5000 | 5 | 5 |
| N2 | Attention | 0.5000 | 5 | 5 |
| N2 | Cross-interaction | 0.5000 | 5 | 5 |

P values are two-sided. “Above chance” requires a positive difference from the metric-specific null and BH-adjusted P < 0.05. Correction uses all 44 tests separately for bootstrap and jackknife; small component counts limit inference.

**Supplementary Table S20.** R3 model-configuration performance before and after giant-component exclusion. Panel A (1 of 3)

| **Row** | **Model** | **Full N0 ROC-AUC** | **Full N2 ROC-AUC** | **Full N0 − N2 gap** | **N0 ROC-AUC after exclusion** |
| --- | --- | --- | --- | --- | --- |
| 1 | LR | 0.6041 | 0.4998 | 0.1043 | 0.6410 |
| 2 | RF | 0.6322 | 0.5184 | 0.1138 | 0.6891 |
| 3 | MLP | 0.6168 | 0.5166 | 0.1002 | 0.7372 |
| 4 | Siamese | 0.6512 | 0.5119 | 0.1393 | 0.8141 |
| 5 | Attention | 0.6245 | 0.5473 | 0.0771 | 0.7179 |
| 6 | Cross-interaction | 0.6132 | 0.4933 | 0.1199 | 0.6731 |

**Supplementary Table S20 (continued).** R3 model-configuration performance before and after giant-component exclusion. Panel B (2 of 3)

| **Row** | **Model** | **N2 ROC-AUC after exclusion** | **N0 − N2 gap after exclusion** | **N0 remaining pairs** | **N2 remaining pairs** |
| --- | --- | --- | --- | --- | --- |
| 1 | LR | 0.3125 | 0.3285 | 25 | 8 |
| 2 | RF | 0.4688 | 0.2204 | 25 | 8 |
| 3 | MLP | 0.6875 | 0.0497 | 25 | 8 |
| 4 | Siamese | 0.4375 | 0.3766 | 25 | 8 |
| 5 | Attention | 0.2500 | 0.4679 | 25 | 8 |
| 6 | Cross-interaction | 0.4375 | 0.2356 | 25 | 8 |

**Supplementary Table S20 (continued).** R3 model-configuration performance before and after giant-component exclusion. Panel C (3 of 3)

| **Row** | **Model** | **Gap direction retained** |
| --- | --- | --- |
| 1 | LR | Yes |
| 2 | RF | Yes |
| 3 | MLP | Yes |
| 4 | Siamese | Yes |
| 5 | Attention | Yes |
| 6 | Cross-interaction | Yes |

The gap is ROC-AUC(N0) − ROC-AUC(N2). Only 25 N0 pairs and eight N2 pairs remain after giant-component removal; the resulting estimates are descriptive. Supplementary Figure S4 shows the delete-one-component jackknife intervals for the same R3 N0/N2 model configurations. The wide intervals, particularly under N2, should be read together with the small number of usable components in Tables S18–S20.

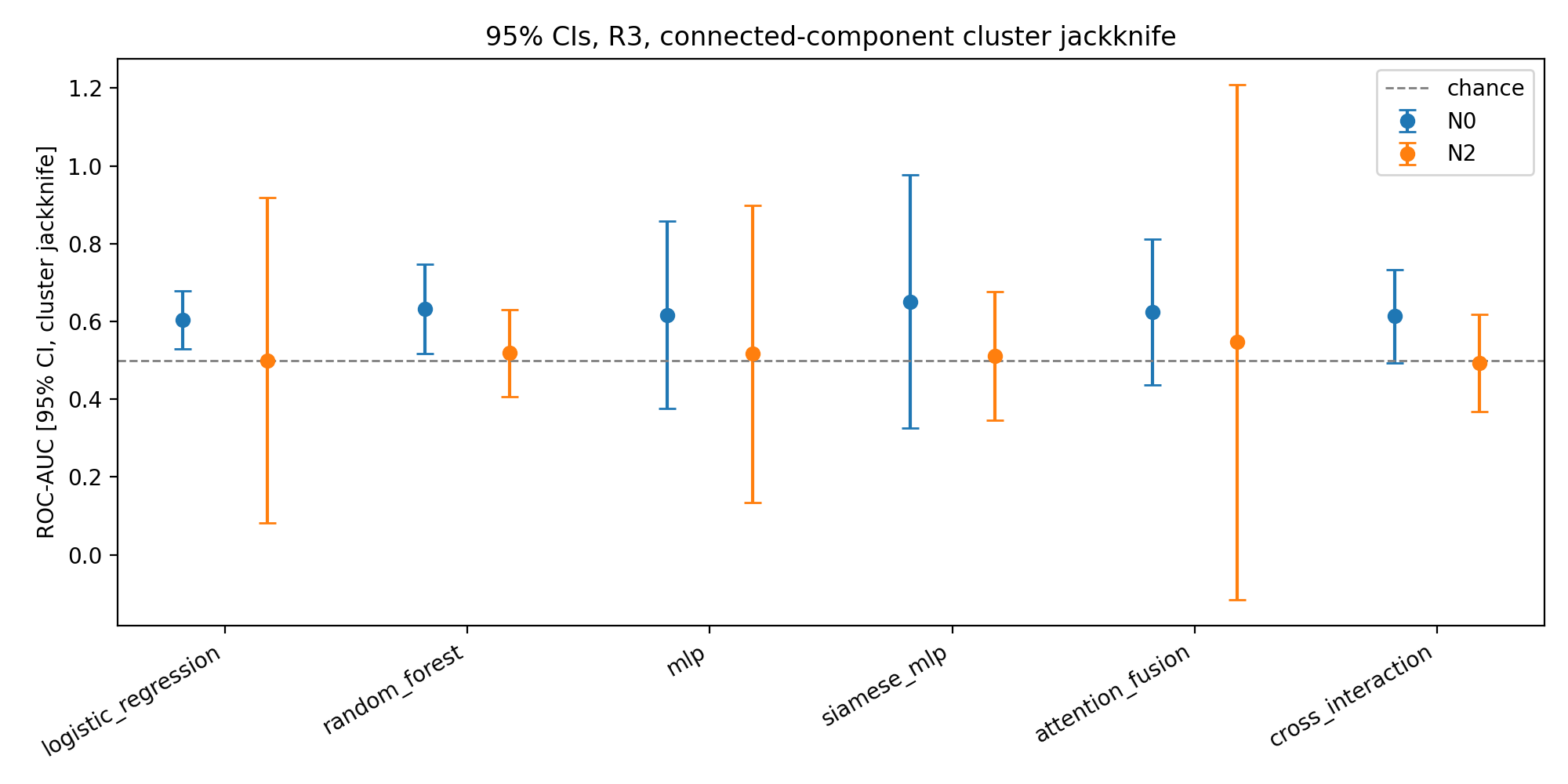

**Supplementary Figure S4.** Component-jackknife uncertainty for R3 model performance under N0 and N2. Points are ROC-AUC estimates and error bars are 95% delete-one-component jackknife intervals. These intervals describe uncertainty under the realized component structure; corrected hypothesis tests are reported in Supplementary Table S19.

**Supplementary Table S21.** Paired R3 ROC-AUC differences: Siamese model minus logistic regression

| **Negative set** | **Comparison** | **Method** | **ΔROC-AUC** | **95% CI lower** | **95% CI upper** | **Raw P value** | **BH-adjusted P value** |
| --- | --- | --- | --- | --- | --- | --- | --- |
| N0 | Siamese − LR | Bootstrap | 0.0470 | 0.0235 | 0.3393 | 0.0240 | 0.0880 |
| N0 | Siamese − LR | Jackknife | 0.0470 | -0.2047 | 0.2988 | 0.6992 | 0.9993 |
| N2 | Siamese − LR | Bootstrap | 0.0120 | -0.0800 | 0.2400 | 0.1520 | 0.2606 |
| N2 | Siamese − LR | Jackknife | 0.0120 | -0.2417 | 0.2657 | 0.9017 | 0.9993 |

The difference is Siamese minus LR on identical test pairs. Intervals are component-aware 95% intervals. Two-sided P values retain BH correction across the complete 44-test family for each method.

**Supplementary Table S22.** Unpaired R3 ROC-AUC differences: N0 minus N2

| **Negative set** | **Model** | **Method** | **ΔROC-AUC** | **95% CI lower** | **95% CI upper** | **Raw P value** | **BH-adjusted P value** |
| --- | --- | --- | --- | --- | --- | --- | --- |
| N0 vs. N2 | LR | Bootstrap | 0.1043 | -0.0392 | 0.5793 | 0.0790 | 0.1580 |
| N0 vs. N2 | LR | Jackknife | 0.1043 | -0.3089 | 0.5174 | 0.5339 | 0.9993 |
| N0 vs. N2 | RF | Bootstrap | 0.1138 | -0.1299 | 0.4136 | 0.2160 | 0.3227 |
| N0 vs. N2 | RF | Jackknife | 0.1138 | -0.0283 | 0.2559 | 0.1099 | 0.6736 |
| N0 vs. N2 | MLP | Bootstrap | 0.1002 | -0.3443 | 0.4198 | 0.3730 | 0.4918 |
| N0 vs. N2 | MLP | Jackknife | 0.1002 | -0.2974 | 0.4978 | 0.5882 | 0.9993 |
| N0 vs. N2 | Siamese | Bootstrap | 0.1393 | -0.0110 | 0.6368 | 0.0530 | 0.1296 |
| N0 vs. N2 | Siamese | Jackknife | 0.1393 | -0.2052 | 0.4838 | 0.4104 | 0.9993 |
| N0 vs. N2 | Attention | Bootstrap | 0.0771 | -0.0009 | 0.6424 | 0.0510 | 0.1296 |
| N0 vs. N2 | Attention | Jackknife | 0.0771 | -0.5709 | 0.7252 | 0.7737 | 0.9993 |
| N0 vs. N2 | Cross-interaction | Bootstrap | 0.1199 | -0.1045 | 0.5050 | 0.1260 | 0.2410 |
| N0 vs. N2 | Cross-interaction | Jackknife | 0.1199 | -0.0338 | 0.2737 | 0.1183 | 0.6736 |

The difference is ROC-AUC(N0) − ROC-AUC(N2) on different test-pair sets. Intervals are component-aware 95% intervals. Two-sided P values retain each method’s complete 44-test BH correction.

**Supplementary Table S23.** R3 leave-one-component-out ROC-AUC ranges and remaining sample sizes

| **Negative set** | **Model** | **Components deleted in turn** | **Minimum ROC-AUC** | **Maximum ROC-AUC** | **Minimum remaining pairs** | **Maximum remaining pairs** |
| --- | --- | --- | --- | --- | --- | --- |
| N0 | Attention | 19 | 0.6217 | 0.7179 | 25 | 461 |
| N0 | Cross-interaction | 19 | 0.6110 | 0.6731 | 25 | 461 |
| N0 | LR | 19 | 0.6025 | 0.6410 | 25 | 461 |
| N0 | MLP | 19 | 0.6148 | 0.7372 | 25 | 461 |
| N0 | RF | 19 | 0.6301 | 0.6891 | 25 | 461 |
| N0 | Siamese | 19 | 0.6486 | 0.8141 | 25 | 461 |
| N2 | Attention | 5 | 0.2500 | 0.5494 | 8 | 444 |
| N2 | Cross-interaction | 5 | 0.4375 | 0.4961 | 8 | 444 |
| N2 | LR | 5 | 0.3125 | 0.5030 | 8 | 444 |
| N2 | MLP | 5 | 0.5136 | 0.6875 | 8 | 444 |
| N2 | RF | 5 | 0.4688 | 0.5224 | 8 | 444 |
| N2 | Siamese | 5 | 0.4375 | 0.5133 | 8 | 444 |

Each deletion removes one whole test-graph component. Minima and maxima are sensitivity ranges, not confidence intervals. Remaining sample sizes are numbers of pairs.

**S12 Negative-generation and degree-source sensitivity**

Across 30 N2 draws, R3 ROC-AUC was 0.489 ± 0.023 for ESM-2 logistic regression, 0.505 ± 0.023 for ESM-2–MLP, and 0.504 ± 0.003 for degree-only logistic regression (mean ± SD). Using training-only degrees changed R0/N2 sequence-model estimates from 0.413 to 0.440 for logistic regression and from 0.463 to 0.482 for MLP. These sensitivities reuse the same biological graph.

**Supplementary Table S24.** Guo N2 performance across 30 negative-generation seeds. Panel A (1 of 3)

| **Row** | **Regime** | **Model** | **ROC-AUC mean** | **ROC-AUC SD** | **ROC-AUC min** | **ROC-AUC max** |
| --- | --- | --- | --- | --- | --- | --- |
| 1 | R3 | Degree-only LR | 0.5041 | 0.0031 | 0.4966 | 0.5099 |
| 2 | R3 | LR | 0.4886 | 0.0227 | 0.4464 | 0.5244 |
| 3 | R3 | MLP | 0.5050 | 0.0226 | 0.4564 | 0.5606 |
| 4 | R0 | Degree-only LR | 0.4990 | 0.0079 | 0.4792 | 0.5150 |
| 5 | R0 | LR | 0.4317 | 0.0102 | 0.4088 | 0.4493 |
| 6 | R0 | MLP | 0.4616 | 0.0103 | 0.4463 | 0.4876 |

**Supplementary Table S24 (continued).** Guo N2 performance across 30 negative-generation seeds. Panel B (2 of 3)

| **Row** | **Regime** | **Model** | **PR-AUC mean** | **PR-AUC SD** | **MCC mean** | **MCC SD** |
| --- | --- | --- | --- | --- | --- | --- |
| 1 | R3 | Degree-only LR | 0.5040 | 0.0053 | 0.0027 | 0.0132 |
| 2 | R3 | LR | 0.4917 | 0.0181 | -0.0199 | 0.0368 |
| 3 | R3 | MLP | 0.5107 | 0.0211 | 0.0048 | 0.0333 |
| 4 | R0 | Degree-only LR | 0.5055 | 0.0080 | -0.0079 | 0.0178 |
| 5 | R0 | LR | 0.4515 | 0.0071 | -0.0962 | 0.0200 |
| 6 | R0 | MLP | 0.4704 | 0.0105 | -0.0546 | 0.0183 |

**Supplementary Table S24 (continued).** Guo N2 performance across 30 negative-generation seeds. Panel C (3 of 3)

| **Row** | **Regime** | **Model** | **Seeds** |
| --- | --- | --- | --- |
| 1 | R3 | Degree-only LR | 30 |
| 2 | R3 | LR | 30 |
| 3 | R3 | MLP | 30 |
| 4 | R0 | Degree-only LR | 30 |
| 5 | R0 | LR | 30 |
| 6 | R0 | MLP | 30 |

The split and model-training seed are fixed. SD describes variation across negative-generation seeds on the same graph, not independent biological replication.

**Supplementary Table S25.** R0 model performance using full-graph or training-only matching degrees. Panel A (1 of 4)

| **Row** | **Degree source** | **Negative set** | **Model** | **Train pairs** | **Test pairs** | **ROC-AUC** | **PR-AUC** |
| --- | --- | --- | --- | --- | --- | --- | --- |
| 1 | Full graph | N0 | LR | 8,930 | 2,234 | 0.7440 | 0.7325 |
| 2 | Full graph | N0 | MLP | 8,930 | 2,234 | 0.7542 | 0.7437 |
| 3 | Full graph | N2 | LR | 8,928 | 2,234 | 0.4130 | 0.4411 |
| 4 | Full graph | N2 | MLP | 8,928 | 2,234 | 0.4627 | 0.4664 |
| 5 | Training only | N0 | LR | 8,930 | 2,234 | 0.7440 | 0.7325 |
| 6 | Training only | N0 | MLP | 8,930 | 2,234 | 0.7542 | 0.7437 |
| 7 | Training only | N2 | LR | 8,928 | 2,234 | 0.4396 | 0.4684 |
| 8 | Training only | N2 | MLP | 8,928 | 2,234 | 0.4822 | 0.4817 |

**Supplementary Table S25 (continued).** R0 model performance using full-graph or training-only matching degrees. Panel B (2 of 4)

| **Row** | **Degree source** | **Negative set** | **Model** | **F1** | **MCC** | **Balanced accuracy** | **Accuracy** |
| --- | --- | --- | --- | --- | --- | --- | --- |
| 1 | Full graph | N0 | LR | 0.6751 | 0.3591 | 0.6795 | 0.6795 |
| 2 | Full graph | N0 | MLP | 0.6840 | 0.3808 | 0.6902 | 0.6902 |
| 3 | Full graph | N2 | LR | 0.4344 | -0.1352 | 0.4324 | 0.4324 |
| 4 | Full graph | N2 | MLP | 0.4242 | -0.0723 | 0.4642 | 0.4642 |
| 5 | Training only | N0 | LR | 0.6751 | 0.3591 | 0.6795 | 0.6795 |
| 6 | Training only | N0 | MLP | 0.6840 | 0.3808 | 0.6902 | 0.6902 |
| 7 | Training only | N2 | LR | 0.4626 | -0.0923 | 0.4539 | 0.4539 |
| 8 | Training only | N2 | MLP | 0.4157 | -0.0530 | 0.4740 | 0.4740 |

**Supplementary Table S25 (continued).** R0 model performance using full-graph or training-only matching degrees. Panel C (3 of 4)

| **Row** | **Degree source** | **Negative set** | **Model** | **Precision** | **Recall** | **Specificity** | **Brier score** |
| --- | --- | --- | --- | --- | --- | --- | --- |
| 1 | Full graph | N0 | LR | 0.6845 | 0.6661 | 0.6929 | 0.2178 |
| 2 | Full graph | N0 | MLP | 0.6980 | 0.6705 | 0.7099 | 0.2528 |
| 3 | Full graph | N2 | LR | 0.4329 | 0.4360 | 0.4288 | 0.3193 |
| 4 | Full graph | N2 | MLP | 0.4584 | 0.3948 | 0.5336 | 0.2779 |
| 5 | Training only | N0 | LR | 0.6845 | 0.6661 | 0.6929 | 0.2178 |
| 6 | Training only | N0 | MLP | 0.6980 | 0.6705 | 0.7099 | 0.2528 |
| 7 | Training only | N2 | LR | 0.4553 | 0.4700 | 0.4378 | 0.3054 |
| 8 | Training only | N2 | MLP | 0.4676 | 0.3742 | 0.5739 | 0.3260 |

**Supplementary Table S25 (continued).** R0 model performance using full-graph or training-only matching degrees. Panel D (4 of 4)

| **Row** | **Degree source** | **Negative set** | **Model** | **ECE** |
| --- | --- | --- | --- | --- |
| 1 | Full graph | N0 | LR | 0.1054 |
| 2 | Full graph | N0 | MLP | 0.2140 |
| 3 | Full graph | N2 | LR | 0.2326 |
| 4 | Full graph | N2 | MLP | 0.1449 |
| 5 | Training only | N0 | LR | 0.1054 |
| 6 | Training only | N0 | MLP | 0.2140 |
| 7 | Training only | N2 | LR | 0.2031 |
| 8 | Training only | N2 | MLP | 0.2477 |

Matching degrees use either all positive pairs or training positives only. This sensitivity analysis concerns R0 and does not provide prospective degrees for unseen R3 proteins.

**S13 Score ordering and inversion**

R0/N2 and R0/N5 predictions showed modest reversed class ordering. Principal R3/N2 estimates were close to chance, including model-configuration logistic regression at ROC-AUC 0.4998. Probability complementation supplies an algebraic diagnostic, not independent validation. Supplementary Figure S5 displays the corresponding class-conditional probability distributions and is intended as a score-ordering diagnostic rather than an additional validation analysis.

**Supplementary Table S26.** Score inversion and class-conditional probability summaries under R0 and R3. Panel A (1 of 4). Ranking

| **Row** | **Regime** | **Negative set** | **Representation** | **Model** | **Test pairs** | **ROC-AUC(p)** | **ROC-AUC(1 − p)** |
| --- | --- | --- | --- | --- | --- | --- | --- |
| 1 | R0 | N0 | AAC+CTD | MLP | 2,234 | 0.7197 | 0.2803 |
| 2 | R0 | N0 | ESM-2 mean | MLP | 2,234 | 0.7542 | 0.2458 |
| 3 | R0 | N2 | AAC+CTD | MLP | 2,234 | 0.4551 | 0.5449 |
| 4 | R0 | N2 | ESM-2 mean | MLP | 2,234 | 0.4627 | 0.5373 |
| 5 | R0 | N5 | AAC+CTD | MLP | 2,234 | 0.4462 | 0.5538 |
| 6 | R0 | N5 | ESM-2 mean | MLP | 2,234 | 0.4628 | 0.5372 |
| 7 | R3 | N0 | AAC+CTD | MLP | 462 | 0.5874 | 0.4126 |
| 8 | R3 | N0 | ESM-2 mean | MLP | 462 | 0.6168 | 0.3832 |
| 9 | R3 | N2 | AAC+CTD | MLP | 446 | 0.5173 | 0.4827 |
| 10 | R3 | N2 | ESM-2 mean | MLP | 446 | 0.5166 | 0.4834 |
| 11 | R3 | N5 | AAC+CTD | MLP | 445 | 0.4618 | 0.5382 |
| 12 | R3 | N5 | ESM-2 mean | MLP | 445 | 0.5074 | 0.4926 |
| 13 | R3 | N2 | ESM-2 mean | LR | 446 | 0.4998 | 0.5002 |
| 14 | R3 | N2 | ESM-2 mean | MLP | 446 | 0.5166 | 0.4834 |

**Supplementary Table S26 (continued).** Score inversion and class-conditional probability summaries under R0 and R3. Panel B (2 of 4). Mean scores and effect size

| **Row** | **Regime** | **Negative set** | **Representation** | **Model** | **Mean probability positive** | **Mean probability negative** | **Cohen’s d (positive − negative)** |
| --- | --- | --- | --- | --- | --- | --- | --- |
| 1 | R0 | N0 | AAC+CTD | MLP | 0.6721 | 0.3926 | 0.7983 |
| 2 | R0 | N0 | ESM-2 mean | MLP | 0.6611 | 0.3063 | 0.9205 |
| 3 | R0 | N2 | AAC+CTD | MLP | 0.4611 | 0.4848 | -0.1703 |
| 4 | R0 | N2 | ESM-2 mean | MLP | 0.4738 | 0.4926 | -0.1395 |
| 5 | R0 | N5 | AAC+CTD | MLP | 0.4723 | 0.4986 | -0.1916 |
| 6 | R0 | N5 | ESM-2 mean | MLP | 0.5215 | 0.5338 | -0.1263 |
| 7 | R3 | N0 | AAC+CTD | MLP | 0.5355 | 0.4579 | 0.3162 |
| 8 | R3 | N0 | ESM-2 mean | MLP | 0.5271 | 0.3922 | 0.3810 |
| 9 | R3 | N2 | AAC+CTD | MLP | 0.4878 | 0.4779 | 0.0991 |
| 10 | R3 | N2 | ESM-2 mean | MLP | 0.4674 | 0.4604 | 0.0546 |
| 11 | R3 | N5 | AAC+CTD | MLP | 0.5592 | 0.5688 | -0.0960 |
| 12 | R3 | N5 | ESM-2 mean | MLP | 0.5197 | 0.5104 | 0.0229 |
| 13 | R3 | N2 | ESM-2 mean | LR | 0.4909 | 0.4912 | -0.0011 |
| 14 | R3 | N2 | ESM-2 mean | MLP | 0.4674 | 0.4604 | 0.0546 |

**Supplementary Table S26 (continued).** Score inversion and class-conditional probability summaries under R0 and R3. Panel C (3 of 4). Median scores and class ordering

| **Row** | **Regime** | **Negative set** | **Representation** | **Model** | **Median probability positive** | **Median probability negative** | **Class ordering** |
| --- | --- | --- | --- | --- | --- | --- | --- |
| 1 | R0 | N0 | AAC+CTD | MLP | 0.8096 | 0.2771 | Positive |
| 2 | R0 | N0 | ESM-2 mean | MLP | 0.9039 | 0.0784 | Positive |
| 3 | R0 | N2 | AAC+CTD | MLP | 0.4530 | 0.4760 | Reversed |
| 4 | R0 | N2 | ESM-2 mean | MLP | 0.4710 | 0.4861 | Reversed |
| 5 | R0 | N5 | AAC+CTD | MLP | 0.4697 | 0.4933 | Reversed |
| 6 | R0 | N5 | ESM-2 mean | MLP | 0.5186 | 0.5338 | Reversed |
| 7 | R3 | N0 | AAC+CTD | MLP | 0.5599 | 0.4926 | Positive |
| 8 | R3 | N0 | ESM-2 mean | MLP | 0.5414 | 0.2618 | Positive |
| 9 | R3 | N2 | AAC+CTD | MLP | 0.4824 | 0.4849 | Near chance |
| 10 | R3 | N2 | ESM-2 mean | MLP | 0.4673 | 0.4579 | Near chance |
| 11 | R3 | N5 | AAC+CTD | MLP | 0.5613 | 0.5838 | Reversed |
| 12 | R3 | N5 | ESM-2 mean | MLP | 0.5295 | 0.5138 | Near chance |
| 13 | R3 | N2 | ESM-2 mean | LR | 0.4855 | 0.5048 | Near chance |
| 14 | R3 | N2 | ESM-2 mean | MLP | 0.4673 | 0.4579 | Near chance |

**Supplementary Table S26 (continued).** Score inversion and class-conditional probability summaries under R0 and R3. Panel D (4 of 4). Score dispersion

| **Row** | **Regime** | **Negative set** | **Representation** | **Model** | **IQR probability positive** | **IQR probability negative** |
| --- | --- | --- | --- | --- | --- | --- |
| 1 | R0 | N0 | AAC+CTD | MLP | 0.5523 | 0.7328 |
| 2 | R0 | N0 | ESM-2 mean | MLP | 0.7498 | 0.6549 |
| 3 | R0 | N2 | AAC+CTD | MLP | 0.1845 | 0.1960 |
| 4 | R0 | N2 | ESM-2 mean | MLP | 0.1712 | 0.1854 |
| 5 | R0 | N5 | AAC+CTD | MLP | 0.1915 | 0.1861 |
| 6 | R0 | N5 | ESM-2 mean | MLP | 0.1256 | 0.1233 |
| 7 | R3 | N0 | AAC+CTD | MLP | 0.3508 | 0.4217 |
| 8 | R3 | N0 | ESM-2 mean | MLP | 0.7112 | 0.6739 |
| 9 | R3 | N2 | AAC+CTD | MLP | 0.1156 | 0.1436 |
| 10 | R3 | N2 | ESM-2 mean | MLP | 0.1627 | 0.1892 |
| 11 | R3 | N5 | AAC+CTD | MLP | 0.1132 | 0.1226 |
| 12 | R3 | N5 | ESM-2 mean | MLP | 0.8897 | 0.9090 |
| 13 | R3 | N2 | ESM-2 mean | LR | 0.4640 | 0.4896 |
| 14 | R3 | N2 | ESM-2 mean | MLP | 0.1627 | 0.1892 |

AUC(1 − p) = 1 − AUC(p). Class differences are positive minus negative. Rows 1–12 use consistent-negative revalidation predictions; rows 13–14 use model-configuration-comparison predictions. Inversion is a diagnostic, not an independently validated correction.

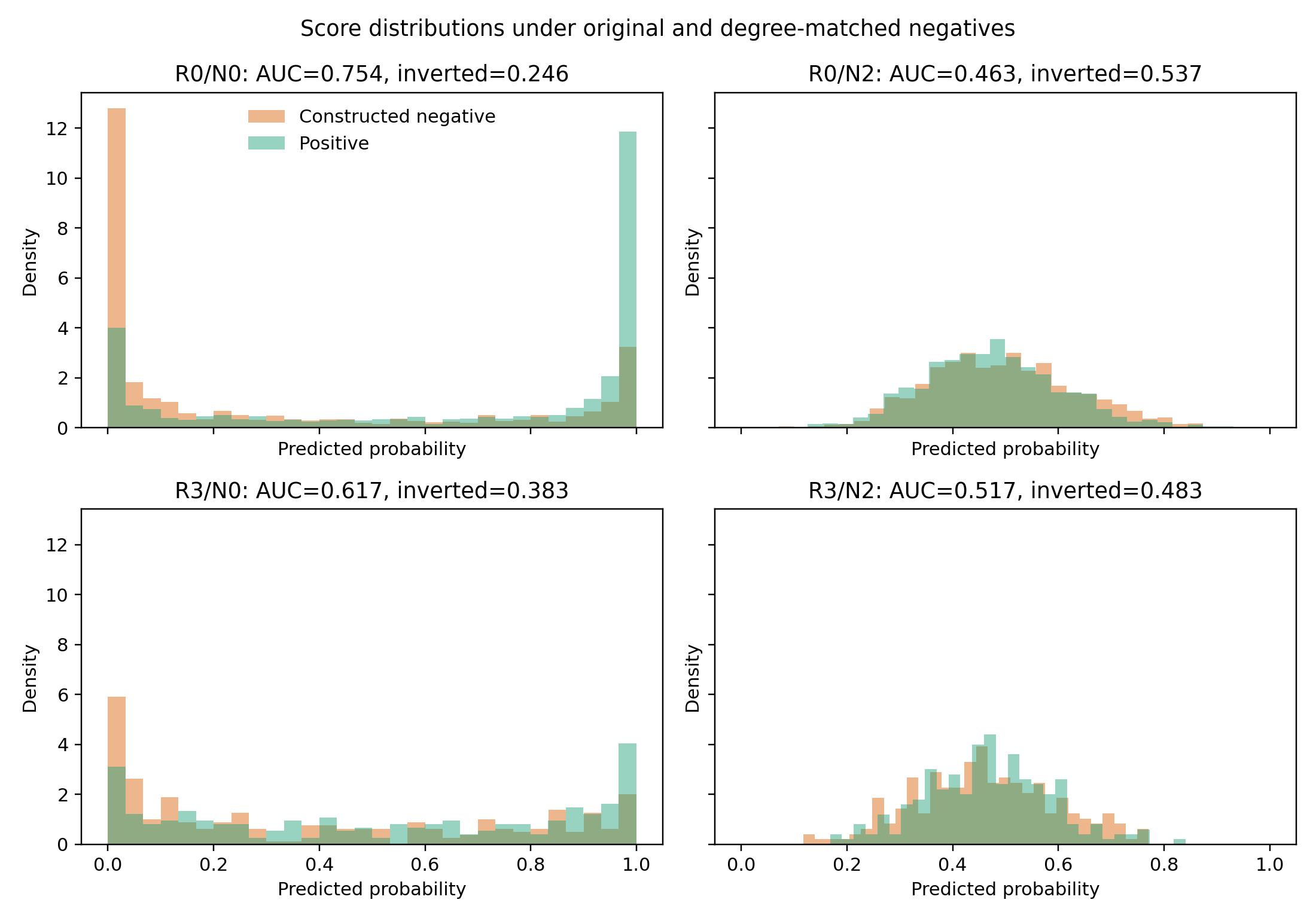

**Supplementary Figure S5. Score distributions under original and degree-matched negatives.** Positive and constructed-negative predicted-probability distributions for the ESM-2–MLP pipeline under R0 and R3, with each negative definition used consistently during training and evaluation.

**S14 Calibration uncertainty and operating points**

In the calibration experiment, R3/N2 MLP MCC was 0.0359 before calibration, −0.0404 after Platt scaling, and 0.0303 after isotonic regression. In the separate model-configuration experiment, precision at recall 0.5 ranged from 0.498 to 0.549 against a positive prevalence of 0.500. Full within-fit calibration and binning results appear in Tables S31–S32.

**Supplementary Table S27.** Raw-probability calibration estimates and component-aware 95% confidence intervals. Panel A (1 of 5). Expected calibration error

| **Row** | **Negative set** | **Model** | **ECE point** | **ECE Boot. 95% CI lower** | **ECE Boot. 95% CI upper** | **ECE Jack. 95% CI lower** | **ECE Jack. 95% CI upper** |
| --- | --- | --- | --- | --- | --- | --- | --- |
| 1 | N0 | LR | 0.3023 | 0.2201 | 0.4499 | 0.2446 | 0.3601 |
| 2 | N0 | RF | 0.0276 | 0.0206 | 0.2462 | -0.1295 | 0.1848 |
| 3 | N0 | MLP | 0.3225 | 0.1716 | 0.4133 | 0.2176 | 0.4274 |
| 4 | N0 | Siamese | 0.2719 | 0.1516 | 0.3811 | 0.2045 | 0.3392 |
| 5 | N0 | Attention | 0.1177 | 0.1104 | 0.4161 | -0.2595 | 0.4948 |
| 6 | N0 | Cross-interaction | 0.0563 | 0.0492 | 0.3378 | -0.1164 | 0.2290 |
| 7 | N2 | LR | 0.2835 | 0.2804 | 0.7103 | -0.2347 | 0.8016 |
| 8 | N2 | RF | 0.0410 | 0.0364 | 0.3803 | -0.3669 | 0.4489 |
| 9 | N2 | MLP | 0.0871 | 0.0828 | 0.3167 | -0.2439 | 0.4181 |
| 10 | N2 | Siamese | 0.0197 | 0.0168 | 0.2876 | 0.0141 | 0.0252 |
| 11 | N2 | Attention | 0.0153 | 0.0150 | 0.2814 | -0.1785 | 0.2091 |
| 12 | N2 | Cross-interaction | 0.0175 | 0.0122 | 0.0198 | 0.0144 | 0.0205 |

**Supplementary Table S27 (continued).** Raw-probability calibration estimates and component-aware 95% confidence intervals. Panel B (2 of 5). Brier score

| **Row** | **Negative set** | **Model** | **Brier point** | **Brier Boot. 95% CI lower** | **Brier Boot. 95% CI upper** | **Brier Jack. 95% CI lower** | **Brier Jack. 95% CI upper** |
| --- | --- | --- | --- | --- | --- | --- | --- |
| 1 | N0 | LR | 0.3474 | 0.1830 | 0.4230 | 0.2535 | 0.4412 |
| 2 | N0 | RF | 0.2344 | 0.1874 | 0.2534 | 0.2016 | 0.2673 |
| 3 | N0 | MLP | 0.3444 | 0.1589 | 0.3781 | 0.1903 | 0.4984 |
| 4 | N0 | Siamese | 0.3050 | 0.1397 | 0.3617 | 0.1891 | 0.4210 |
| 5 | N0 | Attention | 0.2566 | 0.1332 | 0.2881 | 0.1527 | 0.3605 |
| 6 | N0 | Cross-interaction | 0.2354 | 0.1627 | 0.2780 | 0.1902 | 0.2807 |
| 7 | N2 | LR | 0.3472 | 0.3134 | 0.5556 | 0.1293 | 0.5651 |
| 8 | N2 | RF | 0.2506 | 0.2322 | 0.2508 | 0.2158 | 0.2854 |
| 9 | N2 | MLP | 0.2590 | 0.2017 | 0.3265 | 0.2439 | 0.2741 |
| 10 | N2 | Siamese | 0.2521 | 0.2396 | 0.2527 | 0.2383 | 0.2660 |
| 11 | N2 | Attention | 0.2528 | 0.2374 | 0.2530 | 0.2319 | 0.2738 |
| 12 | N2 | Cross-interaction | 0.2503 | 0.2488 | 0.2524 | 0.2492 | 0.2514 |

**Supplementary Table S27 (continued).** Raw-probability calibration estimates and component-aware 95% confidence intervals. Panel C (3 of 5). Calibration slope

| **Row** | **Negative set** | **Model** | **Calibration slope point** | **Calibration slope Boot. 95% CI lower** | **Calibration slope Boot. 95% CI upper** | **Calibration slope Jack. 95% CI lower** | **Calibration slope Jack. 95% CI upper** |
| --- | --- | --- | --- | --- | --- | --- | --- |
| 1 | N0 | LR | 0.0647 | 0.0092 | 0.3069 | -0.0829 | 0.2122 |
| 2 | N0 | RF | 1.2153 | 0.4688 | 5.9013 | -0.1744 | 2.6049 |
| 3 | N0 | MLP | 0.0927 | 0.0288 | 0.4473 | -0.0337 | 0.2190 |
| 4 | N0 | Siamese | 0.1488 | 0.0394 | 0.8700 | -0.1176 | 0.4152 |
| 5 | N0 | Attention | 0.3319 | 0.1061 | 2.4966 | -0.4305 | 1.0943 |
| 6 | N0 | Cross-interaction | 0.7337 | 0.1390 | 5.6163 | -0.1672 | 1.6346 |
| 7 | N2 | LR | -0.0015 | -0.9552 | 0.1876 | -0.6939 | 0.6910 |
| 8 | N2 | RF | 0.2529 | 0.1540 | 33.1846 | -35.8771 | 36.3828 |
| 9 | N2 | MLP | 0.2088 | -0.3848 | 1.7497 | -0.0806 | 0.4981 |
| 10 | N2 | Siamese | -0.2237 | -4.4179 | 48.7159 | -16.1233 | 15.6760 |
| 11 | N2 | Attention | -0.2243 | -0.2706 | 28.4693 | -18.8361 | 18.3875 |
| 12 | N2 | Cross-interaction | 0.5824 | -54.4988 | 21.9038 | -13.9848 | 15.1496 |

**Supplementary Table S27 (continued).** Raw-probability calibration estimates and component-aware 95% confidence intervals. Panel D (4 of 5). Calibration intercept

| **Row** | **Negative set** | **Model** | **Calibration intercept point** | **Calibration intercept Boot. 95% CI lower** | **Calibration intercept Boot. 95% CI upper** | **Calibration intercept Jack. 95% CI lower** | **Calibration intercept Jack. 95% CI upper** |
| --- | --- | --- | --- | --- | --- | --- | --- |
| 1 | N0 | LR | -0.0245 | -0.5892 | 0.6199 | -0.2031 | 0.1541 |
| 2 | N0 | RF | -0.0416 | -0.3119 | 0.7537 | -0.5517 | 0.4686 |
| 3 | N0 | MLP | -0.0556 | -0.4837 | 0.6777 | -0.4265 | 0.3153 |
| 4 | N0 | Siamese | -0.0408 | -0.2880 | 0.9859 | -0.6556 | 0.5740 |
| 5 | N0 | Attention | -0.0249 | -0.4992 | 0.6587 | -0.3024 | 0.2525 |
| 6 | N0 | Cross-interaction | -0.0346 | -0.3463 | 0.7606 | -0.4939 | 0.4247 |
| 7 | N2 | LR | 1.02e-05 | -0.9904 | 0.5393 | -0.1406 | 0.1406 |
| 8 | N2 | RF | -5.8e-15 | -0.2267 | 2.7484 | -1.6899 | 1.6899 |
| 9 | N2 | MLP | -0.0019 | -0.3873 | 0.6799 | -0.4759 | 0.4721 |
| 10 | N2 | Siamese | 0.0177 | -2.8609 | 0.0770 | -0.8555 | 0.8910 |
| 11 | N2 | Attention | 0.0113 | -2.6568 | 0.0136 | -1.2938 | 1.3165 |
| 12 | N2 | Cross-interaction | -0.0407 | -1.0642 | 4.4664 | -0.9853 | 0.9039 |

**Supplementary Table S27 (continued).** Raw-probability calibration estimates and component-aware 95% confidence intervals. Panel E (5 of 5). Resampling units

| **Row** | **Negative set** | **Model** | **Test components** | **Usable components** |
| --- | --- | --- | --- | --- |
| 1 | N0 | LR | 19 | 19 |
| 2 | N0 | RF | 19 | 19 |
| 3 | N0 | MLP | 19 | 19 |
| 4 | N0 | Siamese | 19 | 19 |
| 5 | N0 | Attention | 19 | 19 |
| 6 | N0 | Cross-interaction | 19 | 19 |
| 7 | N2 | LR | 5 | 5 |
| 8 | N2 | RF | 5 | 5 |
| 9 | N2 | MLP | 5 | 5 |
| 10 | N2 | Siamese | 5 | 5 |
| 11 | N2 | Attention | 5 | 5 |
| 12 | N2 | Cross-interaction | 5 | 5 |

Intervals apply to raw probabilities from the calibration experiment. Resampling uses whole test-graph components; calibration slope and intercept refer to the logit of predicted probability. Calibrated point estimates are in Table S31.

**Supplementary Table S28.** Calibration effects on fixed-threshold MCC and separate precision–recall operating points

| **Negative set** | **Model** | **MCC, raw** | **MCC, Platt** | **MCC, isotonic** | **Maximum \|ΔMCC\| after calibration** | **Precision at recall 0.5** | **Recall at precision 0.5** |
| --- | --- | --- | --- | --- | --- | --- | --- |
| N0 | LR | 0.1527 | 0.1441 | 0.1527 | 0.0086 | 0.5803 | 0.9462 |
| N0 | RF | 0.2084 | 0.2238 | 0.1929 | 0.0155 | 0.6129 | 0.9686 |
| N0 | MLP | 0.1337 | 0.1337 | 0.1612 | 0.0275 | 0.5685 | 0.9955 |
| N0 | Siamese | 0.1789 | 0.1789 | 0.2050 | 0.0261 | 0.6400 | 0.9462 |
| N0 | Attention | 0.2054 | 0.2054 | 0.1920 | 0.0134 | 0.5864 | 0.9641 |
| N0 | Cross-interaction | 0.1887 | 0.2335 | 0.1693 | 0.0448 | 0.5895 | 0.9731 |
| N2 | LR | 0.0269 | -0.0225 | 0.0154 | 0.0494 | 0.4978 | 1 |
| N2 | RF | 0.0135 | 0.0269 | 0.0000 | 0.0135 | 0.5488 | 1 |
| N2 | MLP | 0.0359 | -0.0404 | 0.0303 | 0.0762 | 0.5234 | 1 |
| N2 | Siamese | 0.0255 | 0.0184 | 0.0000 | 0.0255 | 0.5283 | 1 |
| N2 | Attention | 0.0651 | 0.0137 | 0.0064 | 0.0587 | 0.5411 | 1 |
| N2 | Cross-interaction | 0.0650 | -0.0135 | 0.0257 | 0.0785 | 0.5045 | 1 |

MCC at threshold 0.5 compares raw and calibrated probabilities from the same fitted model. Precision–recall operating points use the separate model-configuration predictions and are descriptive test-curve summaries, not validated thresholds.

**S15 Pair-Fusion Performance Across Novelty Regimes**

Seven pair-fusion operators were evaluated under R0 and R3 using discrimination, threshold-based classification, and calibration measures. Performance varied substantially across fusion operators and between novelty regimes. Learned symmetric fusion produced ROC-AUC values of 0.9214 under R0 and 0.6512 under R3, whereas the remaining operators showed more modest discrimination. The table also reports F1, MCC, balanced accuracy, accuracy, precision, recall, specificity, Brier score, and ECE, allowing the effect of the fusion operator to be assessed beyond ROC-AUC alone. Learned fusion should be interpreted separately from fixed fusion operators because it also changes model capacity.

**Supplementary Table S29.** Pair-fusion performance under R0 and R3. Panel A (1 of 4)

| **Row** | **Regime** | **Pair fusion** | **Symmetric** | **ROC-AUC** | **PR-AUC** |
| --- | --- | --- | --- | --- | --- |
| 1 | R0 | Sum | Yes | 0.7628 | 0.7564 |
| 2 | R0 | Absolute difference | Yes | 0.6721 | 0.6454 |
| 3 | R0 | Product | Yes | 0.6575 | 0.6433 |
| 4 | R0 | Difference + product | Yes | 0.6961 | 0.6729 |
| 5 | R0 | Combined | Yes | 0.7305 | 0.7162 |
| 6 | R0 | Concatenation | No | 0.7347 | 0.7208 |
| 7 | R0 | Learned symmetric | Yes | 0.9214 | 0.9303 |
| 8 | R3 | Sum | Yes | 0.6089 | 0.5958 |
| 9 | R3 | Absolute difference | Yes | 0.6147 | 0.5721 |
| 10 | R3 | Product | Yes | 0.5879 | 0.5575 |
| 11 | R3 | Difference + product | Yes | 0.6207 | 0.5710 |
| 12 | R3 | Combined | Yes | 0.6168 | 0.5726 |
| 13 | R3 | Concatenation | No | 0.5981 | 0.5942 |
| 14 | R3 | Learned symmetric | Yes | 0.6512 | 0.6306 |

**Supplementary Table S29 (continued).** Pair-fusion performance under R0 and R3. Panel B (2 of 4)

| **Row** | **Regime** | **Pair fusion** | **F1** | **MCC** | **Balanced accuracy** | **Accuracy** |
| --- | --- | --- | --- | --- | --- | --- |
| 1 | R0 | Sum | 0.6958 | 0.3839 | 0.6919 | 0.6919 |
| 2 | R0 | Absolute difference | 0.6408 | 0.2471 | 0.6230 | 0.6229 |
| 3 | R0 | Product | 0.6009 | 0.2065 | 0.6032 | 0.6032 |
| 4 | R0 | Difference + product | 0.6323 | 0.2783 | 0.6390 | 0.6391 |
| 5 | R0 | Combined | 0.6742 | 0.3561 | 0.6780 | 0.6780 |
| 6 | R0 | Concatenation | 0.6730 | 0.3552 | 0.6776 | 0.6776 |
| 7 | R0 | Learned symmetric | 0.8652 | 0.7313 | 0.8657 | 0.8657 |
| 8 | R3 | Sum | 0.5400 | 0.1277 | 0.5637 | 0.5649 |
| 9 | R3 | Absolute difference | 0.5739 | 0.1526 | 0.5763 | 0.5758 |
| 10 | R3 | Product | 0.5455 | 0.1122 | 0.5561 | 0.5563 |
| 11 | R3 | Difference + product | 0.5688 | 0.1718 | 0.5859 | 0.5866 |
| 12 | R3 | Combined | 0.5442 | 0.1488 | 0.5740 | 0.5758 |
| 13 | R3 | Concatenation | 0.5135 | 0.0635 | 0.5318 | 0.5325 |
| 14 | R3 | Learned symmetric | 0.5728 | 0.2226 | 0.6101 | 0.6126 |

**Supplementary Table S29 (continued).** Pair-fusion performance under R0 and R3. Panel C (3 of 4)

| **Row** | **Regime** | **Pair fusion** | **Precision** | **Recall** | **Specificity** | **Brier score** |
| --- | --- | --- | --- | --- | --- | --- |
| 1 | R0 | Sum | 0.6867 | 0.7052 | 0.6786 | 0.2593 |
| 2 | R0 | Absolute difference | 0.6116 | 0.6729 | 0.5730 | 0.2789 |
| 3 | R0 | Product | 0.6042 | 0.5977 | 0.6088 | 0.2602 |
| 4 | R0 | Difference + product | 0.6441 | 0.6210 | 0.6571 | 0.2965 |
| 5 | R0 | Combined | 0.6819 | 0.6667 | 0.6893 | 0.2513 |
| 6 | R0 | Concatenation | 0.6823 | 0.6640 | 0.6911 | 0.2390 |
| 7 | R0 | Learned symmetric | 0.8676 | 0.8629 | 0.8684 | 0.1099 |
| 8 | R3 | Sum | 0.5514 | 0.5291 | 0.5983 | 0.3601 |
| 9 | R3 | Absolute difference | 0.5570 | 0.5919 | 0.5607 | 0.3414 |
| 10 | R3 | Product | 0.5395 | 0.5516 | 0.5607 | 0.3736 |
| 11 | R3 | Difference + product | 0.5727 | 0.5650 | 0.6067 | 0.3387 |
| 12 | R3 | Combined | 0.5652 | 0.5247 | 0.6234 | 0.3123 |
| 13 | R3 | Concatenation | 0.5158 | 0.5112 | 0.5523 | 0.2999 |
| 14 | R3 | Learned symmetric | 0.6122 | 0.5381 | 0.6820 | 0.2828 |

**Supplementary Table S29 (continued).** Pair-fusion performance under R0 and R3. Panel D (4 of 4)

| **Row** | **Regime** | **Pair fusion** | **ECE** |
| --- | --- | --- | --- |
| 1 | R0 | Sum | 0.2313 |
| 2 | R0 | Absolute difference | 0.2154 |
| 3 | R0 | Product | 0.1614 |
| 4 | R0 | Difference + product | 0.2629 |
| 5 | R0 | Combined | 0.1868 |
| 6 | R0 | Concatenation | 0.1650 |
| 7 | R0 | Learned symmetric | 0.0769 |
| 8 | R3 | Sum | 0.3330 |
| 9 | R3 | Absolute difference | 0.3158 |
| 10 | R3 | Product | 0.3410 |
| 11 | R3 | Difference + product | 0.3061 |
| 12 | R3 | Combined | 0.2531 |
| 13 | R3 | Concatenation | 0.2494 |
| 14 | R3 | Learned symmetric | 0.2154 |

Combined fusion concatenates sum, absolute difference, and product. Learned fusion changes model capacity as well as the pair operator.

**S16. ESM-2 Sequence-Truncation Sensitivity**

Table S30 evaluates ESM-2–MLP performance under N-terminal retention, C-terminal retention, full-sequence, and segment-averaging strategies in R0 and R3. Results are reported separately for all, affected, and unaffected test pairs, allowing truncation effects to be distinguished from effects associated with pairs affected by sequence-length handling. Across the full test sets, ROC-AUC ranged from approximately 0.73 to 0.74 in R0 and from approximately 0.61 to 0.62 in R3 across the truncation strategies, with broadly similar patterns for PR-AUC and F1. The affected-pair subsets generally showed lower discrimination than unaffected pairs, particularly under R3.

**Supplementary Table S30.** ESM-2–MLP truncation sensitivity in all, affected, and unaffected test pairs. Panel A (1 of 3)

| **Row** | **Regime** | **Truncation strategy** | **Test subset** | **Test pairs** | **ROC-AUC** | **PR-AUC** | **F1** |
| --- | --- | --- | --- | --- | --- | --- | --- |
| 1 | R0 | N-terminal retention | All pairs | 2,233 | 0.7305 | 0.7162 | 0.6742 |
| 2 | R0 | N-terminal retention | Affected pairs | 481 | 0.6915 | 0.6722 | 0.6465 |
| 3 | R0 | N-terminal retention | Unaffected pairs | 1,752 | 0.7413 | 0.7292 | 0.6822 |
| 4 | R0 | C-terminal retention | All pairs | 2,233 | 0.7205 | 0.7111 | 0.6492 |
| 5 | R0 | C-terminal retention | Affected pairs | 481 | 0.6969 | 0.6798 | 0.6393 |
| 6 | R0 | C-terminal retention | Unaffected pairs | 1,752 | 0.7267 | 0.7202 | 0.6519 |
| 7 | R0 | Full sequence | All pairs | 2,233 | 0.7338 | 0.7272 | 0.6750 |
| 8 | R0 | Full sequence | Affected pairs | 481 | 0.7094 | 0.6976 | 0.6763 |
| 9 | R0 | Full sequence | Unaffected pairs | 1,752 | 0.7404 | 0.7346 | 0.6746 |
| 10 | R0 | Segment averaging | All pairs | 2,233 | 0.7332 | 0.7214 | 0.6619 |
| 11 | R0 | Segment averaging | Affected pairs | 481 | 0.7077 | 0.6859 | 0.6191 |
| 12 | R0 | Segment averaging | Unaffected pairs | 1,752 | 0.7401 | 0.7309 | 0.6740 |
| 13 | R3 | N-terminal retention | All pairs | 462 | 0.6168 | 0.5726 | 0.5442 |
| 14 | R3 | N-terminal retention | Affected pairs | 119 | 0.6016 | 0.4097 | 0.4902 |
| 15 | R3 | N-terminal retention | Unaffected pairs | 343 | 0.6416 | 0.6468 | 0.5610 |
| 16 | R3 | C-terminal retention | All pairs | 462 | 0.6146 | 0.5695 | 0.5670 |
| 17 | R3 | C-terminal retention | Affected pairs | 119 | 0.6139 | 0.4209 | 0.5047 |
| 18 | R3 | C-terminal retention | Unaffected pairs | 343 | 0.6380 | 0.6417 | 0.5865 |
| 19 | R3 | Full sequence | All pairs | 462 | 0.6138 | 0.5744 | 0.5358 |
| 20 | R3 | Full sequence | Affected pairs | 119 | 0.6032 | 0.4170 | 0.4444 |
| 21 | R3 | Full sequence | Unaffected pairs | 343 | 0.6314 | 0.6442 | 0.5629 |
| 22 | R3 | Segment averaging | All pairs | 462 | 0.6150 | 0.5749 | 0.5497 |
| 23 | R3 | Segment averaging | Affected pairs | 119 | 0.6013 | 0.4269 | 0.4902 |
| 24 | R3 | Segment averaging | Unaffected pairs | 343 | 0.6409 | 0.6510 | 0.5680 |

**Supplementary Table S30 (continued).** ESM-2–MLP truncation sensitivity in all, affected, and unaffected test pairs. Panel B (2 of 3)

| **Row** | **Regime** | **Truncation strategy** | **Test subset** | **MCC** | **Balanced accuracy** | **Accuracy** | **Precision** |
| --- | --- | --- | --- | --- | --- | --- | --- |
| 1 | R0 | N-terminal retention | All pairs | 0.3561 | 0.6780 | 0.6780 | 0.6819 |
| 2 | R0 | N-terminal retention | Affected pairs | 0.2727 | 0.6361 | 0.6362 | 0.6299 |
| 3 | R0 | N-terminal retention | Unaffected pairs | 0.3793 | 0.6895 | 0.6895 | 0.6977 |
| 4 | R0 | C-terminal retention | All pairs | 0.3059 | 0.6529 | 0.6529 | 0.6560 |
| 5 | R0 | C-terminal retention | Affected pairs | 0.2682 | 0.6341 | 0.6341 | 0.6316 |
| 6 | R0 | C-terminal retention | Unaffected pairs | 0.3164 | 0.6581 | 0.6581 | 0.6631 |
| 7 | R0 | Full sequence | All pairs | 0.3480 | 0.6740 | 0.6740 | 0.6726 |
| 8 | R0 | Full sequence | Affected pairs | 0.3472 | 0.6736 | 0.6736 | 0.6721 |
| 9 | R0 | Full sequence | Unaffected pairs | 0.3482 | 0.6741 | 0.6741 | 0.6727 |
| 10 | R0 | Segment averaging | All pairs | 0.3256 | 0.6628 | 0.6628 | 0.6634 |
| 11 | R0 | Segment averaging | Affected pairs | 0.2225 | 0.6112 | 0.6112 | 0.6080 |
| 12 | R0 | Segment averaging | Unaffected pairs | 0.3539 | 0.6769 | 0.6769 | 0.6794 |
| 13 | R3 | N-terminal retention | All pairs | 0.1488 | 0.5740 | 0.5758 | 0.5652 |
| 14 | R3 | N-terminal retention | Affected pairs | 0.1481 | 0.5783 | 0.5630 | 0.4032 |
| 15 | R3 | N-terminal retention | Unaffected pairs | 0.1732 | 0.5857 | 0.5802 | 0.6345 |
| 16 | R3 | C-terminal retention | All pairs | 0.1594 | 0.5797 | 0.5801 | 0.5644 |
| 17 | R3 | C-terminal retention | Affected pairs | 0.1606 | 0.5843 | 0.5546 | 0.4030 |
| 18 | R3 | C-terminal retention | Unaffected pairs | 0.1841 | 0.5920 | 0.5889 | 0.6329 |
| 19 | R3 | Full sequence | All pairs | 0.1273 | 0.5634 | 0.5649 | 0.5524 |
| 20 | R3 | Full sequence | Affected pairs | 0.0771 | 0.5408 | 0.5378 | 0.3729 |
| 21 | R3 | Full sequence | Unaffected pairs | 0.1582 | 0.5787 | 0.5743 | 0.6225 |
| 22 | R3 | Segment averaging | All pairs | 0.1534 | 0.5764 | 0.5779 | 0.5667 |
| 23 | R3 | Segment averaging | Affected pairs | 0.1481 | 0.5783 | 0.5630 | 0.4032 |
| 24 | R3 | Segment averaging | Unaffected pairs | 0.1774 | 0.5881 | 0.5831 | 0.6351 |

**Supplementary Table S30 (continued).** ESM-2–MLP truncation sensitivity in all, affected, and unaffected test pairs. Panel C (3 of 3)

| **Row** | **Regime** | **Truncation strategy** | **Test subset** | **Recall** | **Specificity** |
| --- | --- | --- | --- | --- | --- |
| 1 | R0 | N-terminal retention | All pairs | 0.6667 | 0.6893 |
| 2 | R0 | N-terminal retention | Affected pairs | 0.6639 | 0.6083 |
| 3 | R0 | N-terminal retention | Unaffected pairs | 0.6674 | 0.7115 |
| 4 | R0 | C-terminal retention | All pairs | 0.6425 | 0.6634 |
| 5 | R0 | C-terminal retention | Affected pairs | 0.6473 | 0.6208 |
| 6 | R0 | C-terminal retention | Unaffected pairs | 0.6411 | 0.6750 |
| 7 | R0 | Full sequence | All pairs | 0.6774 | 0.6705 |
| 8 | R0 | Full sequence | Affected pairs | 0.6805 | 0.6667 |
| 9 | R0 | Full sequence | Unaffected pairs | 0.6766 | 0.6716 |
| 10 | R0 | Segment averaging | All pairs | 0.6604 | 0.6652 |
| 11 | R0 | Segment averaging | Affected pairs | 0.6307 | 0.5917 |
| 12 | R0 | Segment averaging | Unaffected pairs | 0.6686 | 0.6853 |
| 13 | R3 | N-terminal retention | All pairs | 0.5247 | 0.6234 |
| 14 | R3 | N-terminal retention | Affected pairs | 0.6250 | 0.5316 |
| 15 | R3 | N-terminal retention | Unaffected pairs | 0.5027 | 0.6687 |
| 16 | R3 | C-terminal retention | All pairs | 0.5695 | 0.5900 |
| 17 | R3 | C-terminal retention | Affected pairs | 0.6750 | 0.4937 |
| 18 | R3 | C-terminal retention | Unaffected pairs | 0.5464 | 0.6375 |
| 19 | R3 | Full sequence | All pairs | 0.5202 | 0.6067 |
| 20 | R3 | Full sequence | Affected pairs | 0.5500 | 0.5316 |
| 21 | R3 | Full sequence | Unaffected pairs | 0.5137 | 0.6438 |
| 22 | R3 | Segment averaging | All pairs | 0.5336 | 0.6192 |
| 23 | R3 | Segment averaging | Affected pairs | 0.6250 | 0.5316 |
| 24 | R3 | Segment averaging | Unaffected pairs | 0.5137 | 0.6625 |

Affected pairs contain at least one protein longer than 1,022 residues. N- and C-terminal retention specify which sequence end is kept. Full processing uses the whole sequence; segment averaging combines embeddings from non-overlapping 1,022-residue segments.

**Supplementary Table S31.** Raw, Platt-scaled, and isotonic-calibrated performance from the same R3 fitted models. Panel A (1 of 2)

| **Row** | **Negative set** | **Model** | **Calibration** | **ROC-AUC** | **MCC** | **Brier score** | **ECE** |
| --- | --- | --- | --- | --- | --- | --- | --- |
| 1 | N0 | LR | Raw | 0.5922 | 0.1527 | 0.3474 | 0.3023 |
| 2 | N0 | LR | Platt | 0.5922 | 0.1441 | 0.2640 | 0.1223 |
| 3 | N0 | LR | Isotonic | 0.5942 | 0.1527 | 0.2658 | 0.1318 |
| 4 | N0 | RF | Raw | 0.6414 | 0.2084 | 0.2344 | 0.0276 |
| 5 | N0 | RF | Platt | 0.6414 | 0.2238 | 0.2393 | 0.0744 |
| 6 | N0 | RF | Isotonic | 0.6439 | 0.1929 | 0.2436 | 0.0853 |
| 7 | N0 | MLP | Raw | 0.6144 | 0.1337 | 0.3444 | 0.3225 |
| 8 | N0 | MLP | Platt | 0.6144 | 0.1337 | 0.2526 | 0.1262 |
| 9 | N0 | MLP | Isotonic | 0.6155 | 0.1612 | 0.2569 | 0.1048 |
| 10 | N0 | Siamese | Raw | 0.6461 | 0.1789 | 0.3050 | 0.2719 |
| 11 | N0 | Siamese | Platt | 0.6461 | 0.1789 | 0.2571 | 0.1505 |
| 12 | N0 | Siamese | Isotonic | 0.6400 | 0.2050 | 0.2667 | 0.1664 |
| 13 | N0 | Attention | Raw | 0.6222 | 0.2054 | 0.2566 | 0.1177 |
| 14 | N0 | Attention | Platt | 0.6222 | 0.2054 | 0.2494 | 0.0996 |
| 15 | N0 | Attention | Isotonic | 0.6201 | 0.1920 | 0.2546 | 0.1012 |
| 16 | N0 | Cross-interaction | Raw | 0.6450 | 0.1887 | 0.2354 | 0.0563 |
| 17 | N0 | Cross-interaction | Platt | 0.6450 | 0.2335 | 0.2381 | 0.0647 |
| 18 | N0 | Cross-interaction | Isotonic | 0.6450 | 0.1693 | 0.2445 | 0.0970 |
| 19 | N2 | LR | Raw | 0.5181 | 0.0269 | 0.3472 | 0.2835 |
| 20 | N2 | LR | Platt | 0.4819 | -0.0225 | 0.2540 | 0.0607 |
| 21 | N2 | LR | Isotonic | 0.5045 | 0.0154 | 0.2500 | 0.0011 |
| 22 | N2 | RF | Raw | 0.5043 | 0.0135 | 0.2506 | 0.0410 |
| 23 | N2 | RF | Platt | 0.4957 | 0.0269 | 0.2525 | 0.0204 |
| 24 | N2 | RF | Isotonic | 0.5044 | 0.0000 | 0.2499 | 0.0010 |
| 25 | N2 | MLP | Raw | 0.5379 | 0.0359 | 0.2590 | 0.0871 |
| 26 | N2 | MLP | Platt | 0.4621 | -0.0404 | 0.2514 | 0.0344 |
| 27 | N2 | MLP | Isotonic | 0.5023 | 0.0303 | 0.2498 | 0.0046 |
| 28 | N2 | Siamese | Raw | 0.4940 | 0.0255 | 0.2521 | 0.0197 |
| 29 | N2 | Siamese | Platt | 0.5060 | 0.0184 | 0.2503 | 0.0129 |
| 30 | N2 | Siamese | Isotonic | 0.5089 | 0.0000 | 0.2495 | 0.0096 |
| 31 | N2 | Attention | Raw | 0.4959 | 0.0651 | 0.2528 | 0.0153 |
| 32 | N2 | Attention | Platt | 0.5041 | 0.0137 | 0.2501 | 0.0110 |
| 33 | N2 | Attention | Isotonic | 0.5019 | 0.0064 | 0.2507 | 0.0051 |
| 34 | N2 | Cross-interaction | Raw | 0.5018 | 0.0650 | 0.2503 | 0.0175 |
| 35 | N2 | Cross-interaction | Platt | 0.4982 | -0.0135 | 0.2511 | 0.0301 |
| 36 | N2 | Cross-interaction | Isotonic | 0.5068 | 0.0257 | 0.2514 | 0.0114 |

**Supplementary Table S31 (continued).** Raw, Platt-scaled, and isotonic-calibrated performance from the same R3 fitted models. Panel B (2 of 2)

| **Row** | **Negative set** | **Model** | **Calibration** | **Calibration slope** | **Calibration intercept** |
| --- | --- | --- | --- | --- | --- |
| 1 | N0 | LR | Raw | 0.0647 | -0.0245 |
| 2 | N0 | LR | Platt | 0.2600 | -0.0335 |
| 3 | N0 | LR | Isotonic | 0.0536 | -0.0613 |
| 4 | N0 | RF | Raw | 1.2153 | -0.0416 |
| 5 | N0 | RF | Platt | 0.5855 | -0.0109 |
| 6 | N0 | RF | Isotonic | 0.3817 | -0.0224 |
| 7 | N0 | MLP | Raw | 0.0927 | -0.0556 |
| 8 | N0 | MLP | Platt | 0.4201 | -0.0575 |
| 9 | N0 | MLP | Isotonic | 0.3447 | -0.0387 |
| 10 | N0 | Siamese | Raw | 0.1488 | -0.0408 |
| 11 | N0 | Siamese | Platt | 0.3307 | -0.0379 |
| 12 | N0 | Siamese | Isotonic | 0.1020 | -0.0314 |
| 13 | N0 | Attention | Raw | 0.3319 | -0.0249 |
| 14 | N0 | Attention | Platt | 0.4074 | -0.0249 |
| 15 | N0 | Attention | Isotonic | 0.1426 | -0.0419 |
| 16 | N0 | Cross-interaction | Raw | 0.7337 | -0.0346 |
| 17 | N0 | Cross-interaction | Platt | 0.6005 | 0.0142 |
| 18 | N0 | Cross-interaction | Isotonic | 0.1246 | 0.0086 |
| 19 | N2 | LR | Raw | -0.0015 | 1.02e-05 |
| 20 | N2 | LR | Platt | 0.0140 | -1.66e-15 |
| 21 | N2 | LR | Isotonic | 1.0151 | 0.0035 |
| 22 | N2 | RF | Raw | 0.2529 | -5.8e-15 |
| 23 | N2 | RF | Platt | -0.2053 | 0.0007 |
| 24 | N2 | RF | Isotonic | 3.5725 | 0.0055 |
| 25 | N2 | MLP | Raw | 0.2088 | -0.0019 |
| 26 | N2 | MLP | Platt | -1.6207 | -0.0001 |
| 27 | N2 | MLP | Isotonic | 0.8729 | -0.0064 |
| 28 | N2 | Siamese | Raw | -0.2237 | 0.0177 |
| 29 | N2 | Siamese | Platt | 0.2834 | -0.0049 |
| 30 | N2 | Siamese | Isotonic | 0.0255 | 0.0021 |
| 31 | N2 | Attention | Raw | -0.2243 | 0.0113 |
| 32 | N2 | Attention | Platt | 0.4415 | -0.0083 |
| 33 | N2 | Attention | Isotonic | -0.9223 | -0.0055 |
| 34 | N2 | Cross-interaction | Raw | 0.5824 | -0.0407 |
| 35 | N2 | Cross-interaction | Platt | -0.2013 | 0.0024 |
| 36 | N2 | Cross-interaction | Isotonic | -0.2235 | -0.0036 |

Fit/calibration/test pair counts are 4,562/1,141/462 for N0 and 4,590/1,148/446 for N2. Mappings are fitted on training-derived calibration data. ECE uses 10 equal-width bins; estimates are descriptive.

**Supplementary Table S32.** R3 expected calibration error by bin number and binning strategy. Panel A (1 of 2)

| **Row** | **Negative set** | **Model** | **Bins** | **ECE equal width** | **ECE equal frequency** |
| --- | --- | --- | --- | --- | --- |
| 1 | N0 | LR | 5 | 0.3023 | 0.2978 |
| 2 | N0 | LR | 10 | 0.3023 | 0.2978 |
| 3 | N0 | LR | 15 | 0.3189 | 0.2978 |
| 4 | N0 | LR | 20 | 0.3146 | 0.2984 |
| 5 | N0 | RF | 5 | 0.0107 | 0.0313 |
| 6 | N0 | RF | 10 | 0.0276 | 0.0466 |
| 7 | N0 | RF | 15 | 0.0566 | 0.0635 |
| 8 | N0 | RF | 20 | 0.0349 | 0.0595 |
| 9 | N0 | MLP | 5 | 0.3089 | 0.2798 |
| 10 | N0 | MLP | 10 | 0.3225 | 0.3179 |
| 11 | N0 | MLP | 15 | 0.3203 | 0.3182 |
| 12 | N0 | MLP | 20 | 0.3278 | 0.3179 |
| 13 | N0 | Siamese | 5 | 0.2582 | 0.2633 |
| 14 | N0 | Siamese | 10 | 0.2719 | 0.2633 |
| 15 | N0 | Siamese | 15 | 0.2651 | 0.2633 |
| 16 | N0 | Siamese | 20 | 0.2719 | 0.2633 |
| 17 | N0 | Attention | 5 | 0.1177 | 0.1183 |
| 18 | N0 | Attention | 10 | 0.1177 | 0.1183 |
| 19 | N0 | Attention | 15 | 0.1206 | 0.1251 |
| 20 | N0 | Attention | 20 | 0.1321 | 0.1341 |
| 21 | N0 | Cross-interaction | 5 | 0.0563 | 0.0419 |
| 22 | N0 | Cross-interaction | 10 | 0.0563 | 0.0419 |
| 23 | N0 | Cross-interaction | 15 | 0.0705 | 0.0837 |
| 24 | N0 | Cross-interaction | 20 | 0.0663 | 0.0651 |

**Supplementary Table S32 (continued).** R3 expected calibration error by bin number and binning strategy. Panel B (2 of 2)

| **Row** | **Negative set** | **Model** | **Bins** | **ECE equal width** | **ECE equal frequency** |
| --- | --- | --- | --- | --- | --- |
| 25 | N2 | LR | 5 | 0.2772 | 0.2646 |
| 26 | N2 | LR | 10 | 0.2835 | 0.2753 |
| 27 | N2 | LR | 15 | 0.2857 | 0.2874 |
| 28 | N2 | LR | 20 | 0.2869 | 0.2763 |
| 29 | N2 | RF | 5 | 0.0024 | 0.0506 |
| 30 | N2 | RF | 10 | 0.0410 | 0.0620 |
| 31 | N2 | RF | 15 | 0.0232 | 0.0670 |
| 32 | N2 | RF | 20 | 0.0444 | 0.0928 |
| 33 | N2 | MLP | 5 | 0.0513 | 0.0870 |
| 34 | N2 | MLP | 10 | 0.0871 | 0.0908 |
| 35 | N2 | MLP | 15 | 0.0929 | 0.1083 |
| 36 | N2 | MLP | 20 | 0.1058 | 0.1163 |
| 37 | N2 | Siamese | 5 | 0.0197 | 0.0405 |
| 38 | N2 | Siamese | 10 | 0.0197 | 0.0758 |
| 39 | N2 | Siamese | 15 | 0.0226 | 0.0804 |
| 40 | N2 | Siamese | 20 | 0.0304 | 0.1022 |
| 41 | N2 | Attention | 5 | 0.0153 | 0.0538 |
| 42 | N2 | Attention | 10 | 0.0153 | 0.0622 |
| 43 | N2 | Attention | 15 | 0.0522 | 0.0735 |
| 44 | N2 | Attention | 20 | 0.0510 | 0.0830 |
| 45 | N2 | Cross-interaction | 5 | 0.0175 | 0.0375 |
| 46 | N2 | Cross-interaction | 10 | 0.0175 | 0.0443 |
| 47 | N2 | Cross-interaction | 15 | 0.0258 | 0.0636 |
| 48 | N2 | Cross-interaction | 20 | 0.0175 | 0.0779 |

ECE is computed from raw calibration-experiment predictions using equal-width or equal-frequency bins.

**Supplementary Table S33.** Five-fold cross-validation of separate handcrafted representations

| **Representation** | **Model** | **ROC-AUC mean** | **ROC-AUC SD** | **PR-AUC mean** | **PR-AUC SD** | **MCC mean** | **MCC SD** |
| --- | --- | --- | --- | --- | --- | --- | --- |
| AAC | LR | 0.6092 | 0.0072 | 0.5802 | 0.0091 | 0.1580 | 0.0106 |
| AAC | MLP | 0.6306 | 0.0224 | 0.6064 | 0.0212 | 0.1933 | 0.0307 |
| AAC | RF | 0.6559 | 0.0036 | 0.6231 | 0.0097 | 0.2274 | 0.0136 |
| AAC | XGBoost | 0.6321 | 0.0074 | 0.6104 | 0.0071 | 0.1789 | 0.0216 |
| AAC+CTD | LR | 0.6479 | 0.0201 | 0.6245 | 0.0150 | 0.2066 | 0.0292 |
| AAC+CTD | MLP | 0.7047 | 0.0206 | 0.6805 | 0.0202 | 0.2976 | 0.0390 |
| AAC+CTD | RF | 0.7073 | 0.0141 | 0.6707 | 0.0129 | 0.3105 | 0.0276 |
| AAC+CTD | XGBoost | 0.7065 | 0.0151 | 0.6782 | 0.0137 | 0.3068 | 0.0254 |
| CTD | LR | 0.6438 | 0.0183 | 0.6182 | 0.0150 | 0.2076 | 0.0302 |
| CTD | MLP | 0.7062 | 0.0092 | 0.6790 | 0.0062 | 0.3061 | 0.0145 |
| CTD | RF | 0.7038 | 0.0113 | 0.6681 | 0.0122 | 0.3027 | 0.0135 |
| CTD | XGBoost | 0.6988 | 0.0176 | 0.6699 | 0.0204 | 0.2876 | 0.0274 |
| Physicochemical | LR | 0.5883 | 0.0039 | 0.5583 | 0.0027 | 0.1222 | 0.0259 |
| Physicochemical | MLP | 0.6106 | 0.0139 | 0.5885 | 0.0140 | 0.1555 | 0.0341 |
| Physicochemical | RF | 0.6171 | 0.0053 | 0.5877 | 0.0057 | 0.1689 | 0.0139 |
| Physicochemical | XGBoost | 0.5898 | 0.0093 | 0.5731 | 0.0158 | 0.1180 | 0.0141 |

Stratified pair-level five-fold cross-validation uses seed 42. SD is across folds. Physicochemical descriptors are evaluated separately from AAC+CTD.

**S17. Pooled benchmark matching sensitivity**

Residual pooled sequence-model discrimination remained under exact and nearest matching. Cross-species pairing remains an alternative explanation for pooled discrimination; taxonomy and within-species results are reported in Tables S38-39.

**Supplementary Table S34.** Pooled multispecies model performance under alternative degree-matching rules. Panel A (1 of 3)

| **Row** | **Matching rule** | **Representation** | **Model** | **Train pairs** | **Test pairs** | **ROC-AUC** | **PR-AUC** |
| --- | --- | --- | --- | --- | --- | --- | --- |
| 1 | N2-decile | Degree | LR | 50,684 | 12,672 | 0.5350 | 0.5469 |
| 2 | N2-decile | AAC+CTD | LR | 50,684 | 12,672 | 0.6042 | 0.5970 |
| 3 | N2-decile | AAC+CTD | MLP | 50,684 | 12,672 | 0.6459 | 0.6398 |
| 4 | N2-decile | ESM-2 mean | LR | 50,684 | 12,672 | 0.6815 | 0.6627 |
| 5 | N2-decile | ESM-2 mean | MLP | 50,684 | 12,672 | 0.6976 | 0.6833 |
| 6 | N2-exact | Degree | LR | 49,665 | 12,417 | 0.5056 | 0.5243 |
| 7 | N2-exact | AAC+CTD | LR | 49,665 | 12,417 | 0.6015 | 0.6028 |
| 8 | N2-exact | AAC+CTD | MLP | 49,665 | 12,417 | 0.6114 | 0.6216 |
| 9 | N2-exact | ESM-2 mean | LR | 49,665 | 12,417 | 0.6711 | 0.6606 |
| 10 | N2-exact | ESM-2 mean | MLP | 49,665 | 12,417 | 0.7033 | 0.7033 |
| 11 | N2-nearest | Degree | LR | 50,684 | 12,672 | 0.4904 | 0.4929 |
| 12 | N2-nearest | AAC+CTD | LR | 50,684 | 12,672 | 0.6068 | 0.5937 |
| 13 | N2-nearest | AAC+CTD | MLP | 50,684 | 12,672 | 0.6275 | 0.6191 |
| 14 | N2-nearest | ESM-2 mean | LR | 50,684 | 12,672 | 0.6660 | 0.6455 |
| 15 | N2-nearest | ESM-2 mean | MLP | 50,684 | 12,672 | 0.6908 | 0.6693 |

**Supplementary Table S34 (continued).** Pooled multispecies model performance under alternative degree-matching rules. Panel B (2 of 3)

| **Row** | **Matching rule** | **Representation** | **Model** | **F1** | **MCC** | **Balanced accuracy** | **Accuracy** |
| --- | --- | --- | --- | --- | --- | --- | --- |
| 1 | N2-decile | Degree | LR | 0.4526 | 0.0714 | 0.5341 | 0.5341 |
| 2 | N2-decile | AAC+CTD | LR | 0.5721 | 0.1370 | 0.5685 | 0.5685 |
| 3 | N2-decile | AAC+CTD | MLP | 0.5770 | 0.1950 | 0.5971 | 0.5971 |
| 4 | N2-decile | ESM-2 mean | LR | 0.6356 | 0.2486 | 0.6241 | 0.6241 |
| 5 | N2-decile | ESM-2 mean | MLP | 0.6278 | 0.2588 | 0.6294 | 0.6294 |
| 6 | N2-exact | Degree | LR | 0.5211 | 0.0093 | 0.5046 | 0.5051 |
| 7 | N2-exact | AAC+CTD | LR | 0.5966 | 0.1332 | 0.5661 | 0.5674 |
| 8 | N2-exact | AAC+CTD | MLP | 0.6033 | 0.1459 | 0.5724 | 0.5736 |
| 9 | N2-exact | ESM-2 mean | LR | 0.6366 | 0.2273 | 0.6131 | 0.6141 |
| 10 | N2-exact | ESM-2 mean | MLP | 0.6653 | 0.2667 | 0.6314 | 0.6331 |
| 11 | N2-nearest | Degree | LR | 0.4067 | -0.0171 | 0.4918 | 0.4918 |
| 12 | N2-nearest | AAC+CTD | LR | 0.5771 | 0.1448 | 0.5724 | 0.5724 |
| 13 | N2-nearest | AAC+CTD | MLP | 0.5965 | 0.1712 | 0.5855 | 0.5855 |
| 14 | N2-nearest | ESM-2 mean | LR | 0.6204 | 0.2109 | 0.6051 | 0.6051 |
| 15 | N2-nearest | ESM-2 mean | MLP | 0.6307 | 0.2524 | 0.6262 | 0.6262 |

**Supplementary Table S34 (continued).** Pooled multispecies model performance under alternative degree-matching rules. Panel C (3 of 3)

| **Row** | **Matching rule** | **Representation** | **Model** | **Precision** | **Recall** | **Specificity** |
| --- | --- | --- | --- | --- | --- | --- |
| 1 | N2-decile | Degree | LR | 0.5485 | 0.3853 | 0.6829 |
| 2 | N2-decile | AAC+CTD | LR | 0.5674 | 0.5769 | 0.5601 |
| 3 | N2-decile | AAC+CTD | MLP | 0.6073 | 0.5496 | 0.6446 |
| 4 | N2-decile | ESM-2 mean | LR | 0.6167 | 0.6558 | 0.5923 |
| 5 | N2-decile | ESM-2 mean | MLP | 0.6306 | 0.6250 | 0.6338 |
| 6 | N2-exact | Degree | LR | 0.5147 | 0.5276 | 0.4817 |
| 7 | N2-exact | AAC+CTD | LR | 0.5691 | 0.6269 | 0.5053 |
| 8 | N2-exact | AAC+CTD | MLP | 0.5743 | 0.6354 | 0.5093 |
| 9 | N2-exact | ESM-2 mean | LR | 0.6127 | 0.6626 | 0.5636 |
| 10 | N2-exact | ESM-2 mean | MLP | 0.6223 | 0.7148 | 0.5479 |
| 11 | N2-nearest | Degree | LR | 0.4885 | 0.3483 | 0.6353 |
| 12 | N2-nearest | AAC+CTD | LR | 0.5708 | 0.5836 | 0.5611 |
| 13 | N2-nearest | AAC+CTD | MLP | 0.5811 | 0.6127 | 0.5582 |
| 14 | N2-nearest | ESM-2 mean | LR | 0.5973 | 0.6454 | 0.5649 |
| 15 | N2-nearest | ESM-2 mean | MLP | 0.6232 | 0.6384 | 0.6140 |

Each matching variant is used in both training and evaluation. Pooled N2 negatives may cross species; within-species results are reported in Tables S41–S43.

**S18 Pooled PIPR benchmark performance**

Degree-only logistic-regression ROC-AUC changed from 0.849 under N0 to 0.535 under N2; ESM-2–MLP changed from 0.981 to 0.698. The pooled estimate does not establish compatibility independent of taxonomic pairing (Tables S37–S38).

**Supplementary Table S35.** Pooled multispecies model performance under N0 and degree-matched N2. Panel A (1 of 3)

| **Row** | **Negative set** | **Representation** | **Model** | **Train pairs** | **Test pairs** | **ROC-AUC** | **PR-AUC** |
| --- | --- | --- | --- | --- | --- | --- | --- |
| 1 | N0 | Degree | LR | 51,282 | 12,821 | 0.8492 | 0.8530 |
| 2 | N0 | AAC+CTD | LR | 51,282 | 12,821 | 0.7987 | 0.8092 |
| 3 | N0 | AAC+CTD | MLP | 51,282 | 12,821 | 0.9613 | 0.9692 |
| 4 | N0 | ESM-2 mean | LR | 51,282 | 12,821 | 0.9373 | 0.9488 |
| 5 | N0 | ESM-2 mean | MLP | 51,282 | 12,821 | 0.9809 | 0.9849 |
| 6 | N2 | Degree | LR | 50,684 | 12,672 | 0.5350 | 0.5469 |
| 7 | N2 | AAC+CTD | LR | 50,684 | 12,672 | 0.6042 | 0.5970 |
| 8 | N2 | AAC+CTD | MLP | 50,684 | 12,672 | 0.6459 | 0.6398 |
| 9 | N2 | ESM-2 mean | LR | 50,684 | 12,672 | 0.6815 | 0.6627 |
| 10 | N2 | ESM-2 mean | MLP | 50,684 | 12,672 | 0.6976 | 0.6833 |

**Supplementary Table S35 (continued).** Pooled multispecies model performance under N0 and degree-matched N2. Panel B (2 of 3)

| **Row** | **Negative set** | **Representation** | **Model** | **F1** | **MCC** | **Balanced accuracy** | **Accuracy** |
| --- | --- | --- | --- | --- | --- | --- | --- |
| 1 | N0 | Degree | LR | 0.7322 | 0.5242 | 0.7576 | 0.7586 |
| 2 | N0 | AAC+CTD | LR | 0.7130 | 0.4532 | 0.7259 | 0.7263 |
| 3 | N0 | AAC+CTD | MLP | 0.9053 | 0.8190 | 0.9086 | 0.9090 |
| 4 | N0 | ESM-2 mean | LR | 0.8704 | 0.7494 | 0.8743 | 0.8745 |
| 5 | N0 | ESM-2 mean | MLP | 0.9417 | 0.8884 | 0.9435 | 0.9438 |
| 6 | N2 | Degree | LR | 0.4526 | 0.0714 | 0.5341 | 0.5341 |
| 7 | N2 | AAC+CTD | LR | 0.5721 | 0.1370 | 0.5685 | 0.5685 |
| 8 | N2 | AAC+CTD | MLP | 0.5770 | 0.1950 | 0.5971 | 0.5971 |
| 9 | N2 | ESM-2 mean | LR | 0.6356 | 0.2486 | 0.6241 | 0.6241 |
| 10 | N2 | ESM-2 mean | MLP | 0.6278 | 0.2588 | 0.6294 | 0.6294 |

**Supplementary Table S35 (continued).** Pooled multispecies model performance under N0 and degree-matched N2. Panel C (3 of 3).

| **Row** | **Negative set** | **Representation** | **Model** | **Precision** | **Recall** | **Specificity** |
| --- | --- | --- | --- | --- | --- | --- |
| 1 | N0 | Degree | LR | 0.8103 | 0.6679 | 0.8472 |
| 2 | N0 | AAC+CTD | LR | 0.7400 | 0.6878 | 0.7639 |
| 3 | N0 | AAC+CTD | MLP | 0.9318 | 0.8802 | 0.9371 |
| 4 | N0 | ESM-2 mean | LR | 0.8887 | 0.8529 | 0.8956 |
| 5 | N0 | ESM-2 mean | MLP | 0.9654 | 0.9192 | 0.9678 |
| 6 | N2 | Degree | LR | 0.5485 | 0.3853 | 0.6829 |
| 7 | N2 | AAC+CTD | LR | 0.5674 | 0.5769 | 0.5601 |
| 8 | N2 | AAC+CTD | MLP | 0.6073 | 0.5496 | 0.6446 |
| 9 | N2 | ESM-2 mean | LR | 0.6167 | 0.6558 | 0.5923 |
| 10 | N2 | ESM-2 mean | MLP | 0.6306 | 0.6250 | 0.6338 |

Each negative definition is used in training and evaluation. Cross-species pairing can contribute to pooled N2 discrimination; see the taxonomy diagnostic in Table S38.

**Supplementary Table S36.** Endpoint-degree balance in the pooled multispecies benchmark. Panel A (1 of 4)

| **Row** | **Comparison** | **Mean difference** | **Median difference** | **Cohen’s d** | **SMD** |
| --- | --- | --- | --- | --- | --- |
| 1 | Positive − N0 | 18.4827 | 12 | 0.8272 | 0.8272 |
| 2 | Positive − N2 | 3.6115 | 1 | 0.1301 | 0.1301 |

**Supplementary Table S36 (continued).** Endpoint-degree balance in the pooled multispecies benchmark. Panel B (2 of 4).

| **Row** | **Comparison** | **\|SMD\| < 0.10** | **\|SMD\| < 0.25** | **KS statistic** | **KS P value** |
| --- | --- | --- | --- | --- | --- |
| 1 | Positive − N0 | No | No | 0.4464 | 0.0000 |
| 2 | Positive − N2 | No | Yes | 0.0590 | 2.13e-96 |

**Supplementary Table S36 (continued).** Endpoint-degree balance in the pooled multispecies benchmark. Panel C (3 of 4)

| **Row** | **Comparison** | **Wasserstein distance** | **Distinct-protein mean difference** | **Distinct-protein 95% CI lower** | **Distinct-protein 95% CI upper** |
| --- | --- | --- | --- | --- | --- |
| 1 | Positive − N0 | 18.4827 | -0.4086 | -1.0035 | 0.1577 |
| 2 | Positive − N2 | 3.6115 | -0.6697 | -0.9639 | -0.3725 |

**Supplementary Table S36 (continued).** Endpoint-degree balance in the pooled multispecies benchmark. Panel D (4 of 4)

| **Row** | **Comparison** | **Distinct-protein CI excludes 0** | **Positive-set proteins** | **Negative-set proteins** |
| --- | --- | --- | --- | --- |
| 1 | Positive − N0 | No | 11,205 | 1,399 |
| 2 | Positive − N2 | Yes | 11,205 | 9,733 |

Differences are positive minus negative. Endpoint-occurrence statistics are descriptive; distinct-protein mean differences use a separate 95% bootstrap interval. These intervals are not component-aware model tests.

**S19 Taxonomy audit and within-species replication**

Of 11,529 input proteins, 30 belonged to other organisms and six remained unresolved. Exclusions removed 79 positive and 511 N0 pairs from the species-specific analyses. Within-species decile-matched ESM-2–MLP ROC-AUC was 0.596 for C. elegans, 0.567 for D. melanogaster, and 0.647 for E. coli. Exact matching omitted proposed negatives for 697, 703, and 1,410 retained positive pairs, respectively. Nearest matching retained one negative per positive. In the reconstructed pooled set, 14,284 of 31,678 N2 negatives crossed species (45.1%); the same-species indicator yielded ROC-AUC 0.729 on 12,642 assigned test pairs. All within-species model results use single random pair splits and permit protein recurrence.

**Supplementary Table S37.** Taxonomic composition of the reconstructed pooled N2 negative set

| **Pair category** | **Negative pairs** | **Percent** |
| --- | --- | --- |
| C. elegans | 640 | 2.0203 |
| Cross-species | 14,284 | 45.0912 |
| D. melanogaster | 15,758 | 49.7443 |
| E. coli | 908 | 2.8663 |
| Other-organism endpoint | 69 | 0.2178 |
| Unresolved endpoint | 19 | 0.0600 |

Percentages use all 31,678 reconstructed pooled N2 negatives as the denominator. Cross-species pairs have endpoints assigned to different species.

**Supplementary Table S38.** Same-species indicator performance on pooled N0 and N2 test pairs

| **Set** | **Test pairs** | **Excluded pairs** | **ROC-AUC** | **PR-AUC** | **MCC** |
| --- | --- | --- | --- | --- | --- |
| N0 | 12,698 | 123 | 0.5000 | 0.4980 | 0.0000 |
| N2 | 12,642 | 30 | 0.7285 | 0.6482 | 0.5443 |

The score is 1 for endpoints from the same target species and 0 otherwise. Excluded pairs have at least one endpoint outside the three resolved target species. No classifier is fitted.

**Supplementary Table S39.** Species-specific dataset sizes after taxonomy-based separation

| **Species** | **Proteins** | **Positive pairs** | **N0 pairs** | **Total pairs** | **Zero-degree proteins** |
| --- | --- | --- | --- | --- | --- |
| C. elegans | 2,629 | 3,950 | 3,914 | 7,864 | 11 |
| D. melanogaster | 7,041 | 21,742 | 21,046 | 42,788 | 14 |
| E. coli | 1,823 | 5,907 | 6,954 | 12,861 | 303 |

Zero degree means no partner in the observed within-species positive graph. Counts include only retained within-species pairs.

**Supplementary Table S40.** Retained and excluded pair counts in the combined benchmark

| **Pair category** | **Negative pairs** | **Positive pairs** |
| --- | --- | --- |
| C. elegans | 3,914 | 3,950 |
| D. melanogaster | 21,046 | 21,742 |
| E. coli | 6,954 | 5,907 |
| Other-organism endpoint | 511 | 61 |
| Unresolved endpoint | 0.0000 | 18 |

Pair categories account for all positive and original-negative pairs in the cleaned combined input. Excluded pairs contain an endpoint outside the target species or an unresolved endpoint.

**Supplementary Table S41.** C. elegans model performance under original and degree-matched negatives

| **Set** | **Representation** | **Model** | **Train pairs** | **Test pairs** | **ROC-AUC** | **PR-AUC** | **MCC** |
| --- | --- | --- | --- | --- | --- | --- | --- |
| N0 | Degree | LR | 6,291 | 1,573 | 0.8680 | 0.8814 | 0.5959 |
| N0 | AAC+CTD | LR | 6,291 | 1,573 | 0.9245 | 0.9395 | 0.7250 |
| N0 | AAC+CTD | MLP | 6,291 | 1,573 | 0.9786 | 0.9847 | 0.8933 |
| N0 | ESM-2 mean | LR | 6,291 | 1,573 | 0.9836 | 0.9869 | 0.8846 |
| N0 | ESM-2 mean | MLP | 6,291 | 1,573 | 0.9828 | 0.9860 | 0.8912 |
| N2-decile | Degree | LR | 6,320 | 1,580 | 0.5831 | 0.5879 | 0.1572 |
| N2-decile | AAC+CTD | LR | 6,320 | 1,580 | 0.5494 | 0.5499 | 0.0646 |
| N2-decile | AAC+CTD | MLP | 6,320 | 1,580 | 0.5346 | 0.5499 | 0.0422 |
| N2-decile | ESM-2 mean | LR | 6,320 | 1,580 | 0.5310 | 0.5476 | 0.0380 |
| N2-decile | ESM-2 mean | MLP | 6,320 | 1,580 | 0.5964 | 0.6076 | 0.1456 |
| N2-exact | Degree | LR | 5,762 | 1,441 | 0.5835 | 0.6358 | 0.0245 |
| N2-exact | AAC+CTD | LR | 5,762 | 1,441 | 0.5379 | 0.6009 | 0.0321 |
| N2-exact | AAC+CTD | MLP | 5,762 | 1,441 | 0.5413 | 0.6033 | 0.0601 |
| N2-exact | ESM-2 mean | LR | 5,762 | 1,441 | 0.5230 | 0.5807 | 0.0420 |
| N2-exact | ESM-2 mean | MLP | 5,762 | 1,441 | 0.5736 | 0.6326 | 0.0693 |
| N2-nearest | Degree | LR | 6,320 | 1,580 | 0.5023 | 0.5282 | -0.0013 |
| N2-nearest | AAC+CTD | LR | 6,320 | 1,580 | 0.5591 | 0.5647 | 0.0810 |
| N2-nearest | AAC+CTD | MLP | 6,320 | 1,580 | 0.5601 | 0.5644 | 0.0650 |
| N2-nearest | ESM-2 mean | LR | 6,320 | 1,580 | 0.5285 | 0.5231 | 0.0494 |
| N2-nearest | ESM-2 mean | MLP | 6,320 | 1,580 | 0.5704 | 0.5571 | 0.1114 |

Results use one 80/20 random pair split (seed 42) with within-species degree matching. Counts reflect matching omissions; train/test proteins may recur.

**Supplementary Table S42. D.** melanogaster model performance under original and degree-matched negatives

| **Set** | **Representation** | **Model** | **Train pairs** | **Test pairs** | **ROC-AUC** | **PR-AUC** | **MCC** |
| --- | --- | --- | --- | --- | --- | --- | --- |
| N0 | Degree | LR | 34,229 | 8,559 | 0.8295 | 0.8308 | 0.4927 |
| N0 | AAC+CTD | LR | 34,229 | 8,559 | 0.8343 | 0.8590 | 0.5113 |
| N0 | AAC+CTD | MLP | 34,229 | 8,559 | 0.9755 | 0.9813 | 0.8611 |
| N0 | ESM-2 mean | LR | 34,229 | 8,559 | 0.9683 | 0.9763 | 0.8353 |
| N0 | ESM-2 mean | MLP | 34,229 | 8,559 | 0.9853 | 0.9891 | 0.9001 |
| N2-decile | Degree | LR | 34,786 | 8,698 | 0.5357 | 0.5386 | 0.0649 |
| N2-decile | AAC+CTD | LR | 34,786 | 8,698 | 0.5546 | 0.5460 | 0.0722 |
| N2-decile | AAC+CTD | MLP | 34,786 | 8,698 | 0.5328 | 0.5279 | 0.0397 |
| N2-decile | ESM-2 mean | LR | 34,786 | 8,698 | 0.5562 | 0.5552 | 0.0784 |
| N2-decile | ESM-2 mean | MLP | 34,786 | 8,698 | 0.5668 | 0.5768 | 0.0935 |
| N2-exact | Degree | LR | 34,224 | 8,557 | 0.5196 | 0.5302 | 0.0222 |
| N2-exact | AAC+CTD | LR | 34,224 | 8,557 | 0.5458 | 0.5523 | 0.0636 |
| N2-exact | AAC+CTD | MLP | 34,224 | 8,557 | 0.5320 | 0.5431 | 0.0409 |
| N2-exact | ESM-2 mean | LR | 34,224 | 8,557 | 0.5412 | 0.5577 | 0.0576 |
| N2-exact | ESM-2 mean | MLP | 34,224 | 8,557 | 0.5598 | 0.5757 | 0.0729 |
| N2-nearest | Degree | LR | 34,786 | 8,698 | 0.4953 | 0.4945 | -0.0112 |
| N2-nearest | AAC+CTD | LR | 34,786 | 8,698 | 0.5526 | 0.5516 | 0.0563 |
| N2-nearest | AAC+CTD | MLP | 34,786 | 8,698 | 0.5459 | 0.5448 | 0.0619 |
| N2-nearest | ESM-2 mean | LR | 34,786 | 8,698 | 0.5411 | 0.5412 | 0.0529 |
| N2-nearest | ESM-2 mean | MLP | 34,786 | 8,698 | 0.5591 | 0.5612 | 0.0895 |

Results use one 80/20 random pair split (seed 42) with within-species degree matching. Counts reflect matching omissions; train/test proteins may recur.

**Supplementary Table S43.** E. coli model performance under original and degree-matched negatives

| **Set** | **Representation** | **Model** | **Train pairs** | **Test pairs** | **ROC-AUC** | **PR-AUC** | **MCC** |
| --- | --- | --- | --- | --- | --- | --- | --- |
| N0 | Degree | LR | 10,288 | 2,573 | 0.9511 | 0.9536 | 0.7866 |
| N0 | AAC+CTD | LR | 10,288 | 2,573 | 0.9357 | 0.9443 | 0.7399 |
| N0 | AAC+CTD | MLP | 10,288 | 2,573 | 0.9659 | 0.9729 | 0.8589 |
| N0 | ESM-2 mean | LR | 10,288 | 2,573 | 0.9682 | 0.9725 | 0.8411 |
| N0 | ESM-2 mean | MLP | 10,288 | 2,573 | 0.9773 | 0.9807 | 0.8859 |
| N2-decile | Degree | LR | 9,450 | 2,364 | 0.5832 | 0.5686 | 0.1559 |
| N2-decile | AAC+CTD | LR | 9,450 | 2,364 | 0.5661 | 0.5484 | 0.1049 |
| N2-decile | AAC+CTD | MLP | 9,450 | 2,364 | 0.5889 | 0.5651 | 0.1456 |
| N2-decile | ESM-2 mean | LR | 9,450 | 2,364 | 0.5761 | 0.5702 | 0.1142 |
| N2-decile | ESM-2 mean | MLP | 9,450 | 2,364 | 0.6468 | 0.6355 | 0.2174 |
| N2-exact | Degree | LR | 8,322 | 2,082 | 0.5805 | 0.6453 | 0.0604 |
| N2-exact | AAC+CTD | LR | 8,322 | 2,082 | 0.5658 | 0.6282 | 0.0523 |
| N2-exact | AAC+CTD | MLP | 8,322 | 2,082 | 0.5720 | 0.6409 | 0.1073 |
| N2-exact | ESM-2 mean | LR | 8,322 | 2,082 | 0.5566 | 0.6067 | 0.0824 |
| N2-exact | ESM-2 mean | MLP | 8,322 | 2,082 | 0.6122 | 0.6787 | 0.1500 |
| N2-nearest | Degree | LR | 9,450 | 2,364 | 0.5075 | 0.5097 | 0.0042 |
| N2-nearest | AAC+CTD | LR | 9,450 | 2,364 | 0.5258 | 0.5193 | 0.0321 |
| N2-nearest | AAC+CTD | MLP | 9,450 | 2,364 | 0.5611 | 0.5402 | 0.0986 |
| N2-nearest | ESM-2 mean | LR | 9,450 | 2,364 | 0.5174 | 0.5144 | 0.0389 |
| N2-nearest | ESM-2 mean | MLP | 9,450 | 2,364 | 0.5959 | 0.5967 | 0.1396 |

Results use one 80/20 random pair split (seed 42) with within-species degree matching. Counts reflect matching omissions; train/test proteins may recur.

**Supplementary Table S44.** Within-species degree balance and matching omissions

| **Species** | **Set** | **Negatives** | **Skipped substitutions** | **SMD** | **KS distance** | **Wasserstein distance** |
| --- | --- | --- | --- | --- | --- | --- |
| C. elegans | N0 | 3,914 | 0.0000 | 0.6579 | 0.3988 | 16.3521 |
| C. elegans | N2-decile | 3,950 | 0.0000 | 0.1775 | 0.0847 | 5.5292 |
| C. elegans | N2-exact | 3,253 | 697 | 0.1602 | 0.0644 | 5.0728 |
| C. elegans | N2-nearest | 3,950 | 0.0000 | 0.0391 | 0.0128 | 1.2801 |
| D. melanogaster | N0 | 21,046 | 0.0000 | 0.7352 | 0.3970 | 13.5718 |
| D. melanogaster | N2-decile | 21,742 | 0.0000 | 0.0981 | 0.0432 | 2.1663 |
| D. melanogaster | N2-exact | 21,039 | 703 | 0.0573 | 0.0144 | 1.2842 |
| D. melanogaster | N2-nearest | 21,742 | 0.0000 | 0.0053 | 0.0019 | 0.1400 |
| E. coli | N0 | 6,954 | 0.0000 | 1.3407 | 0.6425 | 37.5447 |
| E. coli | N2-decile | 5,907 | 0.0000 | 0.1915 | 0.0968 | 7.0403 |
| E. coli | N2-exact | 4,497 | 1,410 | 0.1959 | 0.0896 | 7.3735 |
| E. coli | N2-nearest | 5,907 | 0.0000 | 0.0192 | 0.0112 | 0.7423 |

Skipped substitutions omit a proposed negative, while the corresponding positive remains. SMD, KS, and Wasserstein distance compare positive and negative endpoint-degree distributions.

**Supplementary Table S45.** Species-specific train/test class counts and protein overlap

| **Species** | **Set** | **Train positives** | **Train negatives** | **Test positives** | **Test negatives** | **Train/test shared proteins** |
| --- | --- | --- | --- | --- | --- | --- |
| C. elegans | N0 | 3,160 | 3,131 | 790 | 783 | 711 |
| C. elegans | N2-decile | 3,160 | 3,160 | 790 | 790 | 1,226 |
| C. elegans | N2-exact | 3,160 | 2,602 | 790 | 651 | 1,130 |
| C. elegans | N2-nearest | 3,160 | 3,160 | 790 | 790 | 1,211 |
| D. melanogaster | N0 | 17,393 | 16,836 | 4,349 | 4,210 | 3,440 |
| D. melanogaster | N2-decile | 17,393 | 17,393 | 4,349 | 4,349 | 4,558 |
| D. melanogaster | N2-exact | 17,393 | 16,831 | 4,349 | 4,208 | 4,530 |
| D. melanogaster | N2-nearest | 17,393 | 17,393 | 4,349 | 4,349 | 4,553 |
| E. coli | N0 | 4,725 | 5,563 | 1,182 | 1,391 | 961 |
| E. coli | N2-decile | 4,725 | 4,725 | 1,182 | 1,182 | 870 |
| E. coli | N2-exact | 4,725 | 3,597 | 1,182 | 900 | 857 |
| E. coli | N2-nearest | 4,725 | 4,725 | 1,182 | 1,182 | 851 |

Positive and negative counts refer to pairs. Shared proteins occur in both training and test sets.

Supplementary Figure S6 provides a compact cross-analysis view of four central comparisons: the matched protein-novelty curve, the independently generated sequence-cluster-controlled R3 sensitivities, Guo degree-only discrimination under N0 versus N2, and pooled PIPR ESM-2–MLP discrimination under N0 versus N2. It is a visual summary of results reported in Supplementary Tables S35–S36, not an additional statistical analysis.

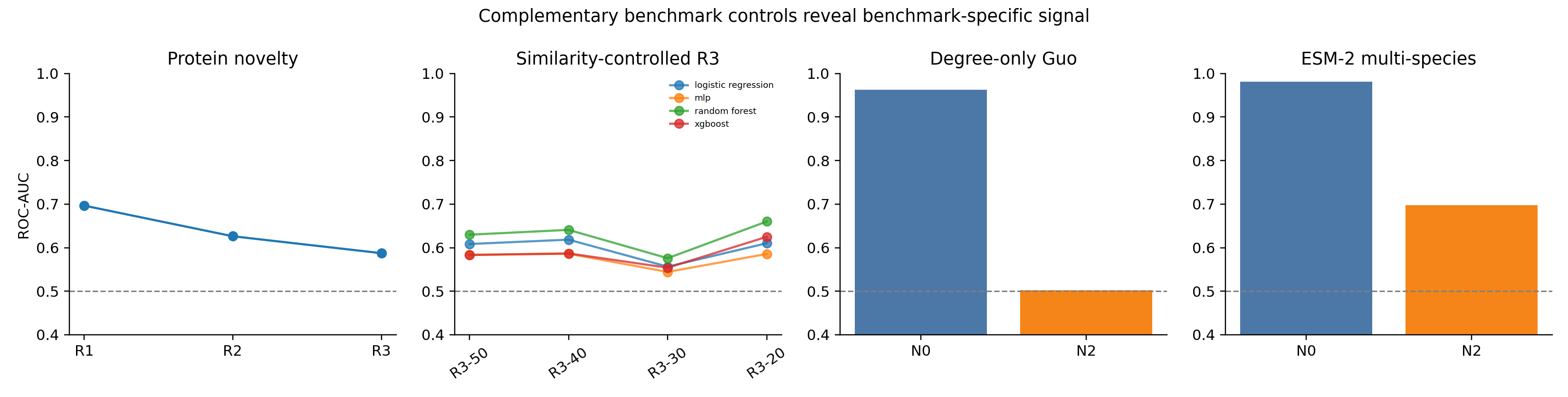

**Supplementary Figure S6.** Integrated central findings. (A) Matched Guo protein-novelty curve. (B) Independently generated sequence-cluster-controlled R3 variants. (C) Guo degree-only discrimination under N0 and N2. (D) Pooled multispecies ESM-2–MLP discrimination under N0 and N2. Corresponding values are reported in Supplementary Tables S35–S36.
